# Seasonal patterns of environmental DNA detection for freshwater unionid mussels

**DOI:** 10.64898/2026.02.19.706871

**Authors:** Nathaniel Marshall, Dan Symonds, Charlie Allen, Noah Berg, Cheryl Dean, Mae Sierra, W. Cody Fleece

## Abstract

Environmental DNA (eDNA) provides a powerful non-invasive tool for monitoring freshwater mussel assemblages, yet detection probabilities can be influenced by reproductive behaviors, seasonal vertical migration, and hydrological conditions. This study assessed eDNA detection from April through October across two diverse mussel beds in Ohio, encompassing species with both tachytictic (short-term brooders) and bradytictic (long-term brooders) reproductive strategies. Mussel DNA was consistently detected across seasons, with detection patterns generally aligning with species observed through a visual tactile survey. Overall, the eDNA sequence abundance was positively correlated with tactile mussel counts, however congruence between the two surveys was strongest during low discharge and when the surveys occurred in close temporal proximity to one another. This study finds that eDNA sampling for freshwater mussels performs adequately within the currently prescribed survey window for visual surveys. However, seasonal factors such as endobenthic burial behavior and high discharge events may have reduced detection efficiency, particularly in Killbuck Creek, where species richness was lowest during periods of high flow in early spring. Therefore, decisions made regarding the timing of eDNA surveys should consider local environmental conditions (e.g., temperature and flow) to achieve optimal results.

## Introduction

Of the 303 currently recognized mussel species within the U.S. (FMCS 2021), nearly one third are federally endangered (77 species) or federally threatened (18 species) (USFWS 2023). Due to their rarity and the fact that unionids burrow into the benthic substrate, traditional visual surveys are time-consuming and require expert taxonomists. Therefore, novel methods to improve the efficiency and accuracy of surveys could be beneficial for population management.

The analysis of environmental DNA (eDNA - genetic material released from urine, waste, mucus, or sloughed cells) is increasingly integrated into natural resource surveys designed to detect the presence of special status species and to describe entire assemblages (Beng & Corlett 2020, Deiner et al. 2021).

Environmental DNA surveys can be more sensitive, less costly, less intrusive on the environment, and provide improved sampling capabilities for challenging and remote habitats (Jerde 2021, Sternhagen et al. 2024). Therefore, eDNA metabarcoding, a technique to simultaneously identify multiple species within a single sample, is being explored as a way to assess mussel assemblage characteristics in river systems (Marshall et al. 2022, Marshall & Fleece 2025, Prié et al. 2025).

While the benefits of eDNA have been widely documented (Beng & Corlett 2020, Fediajevaite et al. 2021), direct uptake and implementation of eDNA for environmental consultation and permitting purposes has been rare within the U.S. (Laschever et al. 2023). One current limitation for wider adoption of eDNA for surveying freshwater mussels is the lack of understanding in how eDNA performs compared to visual tactile protocols. Uncertainties and limitations associated with visual mussel surveys are often outlined within prescribed protocols (see Mussel Survey Guidelines and Protocols – https://molluskconservation.org/Mussel_Protocols.html); and while several protocols for best practices broadly exist for eDNA sampling (e.g., Taberlet et al. 2018, Sellers et al. 2024), specific documentation detailing potential limitations and uncertainties pertaining to the detection of freshwater mussels with eDNA are not available.

Several factors contribute to imperfect detection of freshwater mussels during visual searches. Large-bodied species and species with sculptured shells are typically detected at higher rates than small-bodied ones (Hornbach & Deneka 1996, Vaughn et al. 1997, Sanchez & Schwalb 2019). Visual detectability can also be influenced by vertical migration, as epibenthic species are easier to find than endobenthic ones (Amyot & Downing 1991, Sanchez & Schwalb 2019). Mussels can exhibit seasonal patterns in vertical migration associated with day length, water temperature, flow discharge, and reproductive behavior (Amyot & Downing 1991, Watters et al. 2001, Perles et al. 2003, Schwalb & Pusch 2007), which will influence seasonal detectability during a visual search (Villella et al. 2004). Several biological factors such as gender and age demographics and reproductive activity can also influence detection (Smith et al. 2001). Environmental factors and habitat conditions such as substrate type, water clarity, flow, and depth, can also influence the visibility of freshwater mussels and affect the ease at which individuals are recovered from substrate (Smith 2006, Wisniewski et al. 2013, Sanchez & Schwalb 2019). In addition to biological and environmental factors, survey experience and fatigue of the observers can affect freshwater mussel detectability (Wisniewski et al. 2013, Reid 2016). While these factors influencing visual detection probability for freshwater mussels are well documented, detailed understanding of factors pertaining to the detection of freshwater mussel eDNA have been less intensively investigated.

Environmental DNA concentrations may fluctuate seasonally due to reproductive and life history behavioral responses as well as environmental factors. For example, eDNA detection probability of *Necturus alabamensis* (Black Warrior Waterdog) was found to be greatest in colder seasons when they were more active, while eDNA detection probability of *Sternotherus depressus (*Flattened Musk Turtle) was found to be greatest in warm seasons when this species is more active (de Souza et al. 2016).

Environmental DNA concentrations for *Austropotamobius pallipes* (white-clawed crayfish) can vary seasonally, with highest concentrations corresponding to heightened activity during reproduction and juvenile release, and the lowest concentrations associated with colder months during reduced activity (Troth et al. 2021). Two submerged macrophytes (*Potamogeton pusillus* and *Stuckenia pectinata*) had the greatest eDNA concentrations during peak growth associated with warm Summer months (Takahara et al. 2025). Additionally, seasonal life-history behaviors such as spawning can largely influence detection patterns from eDNA. For instance, eDNA concentrations of *Oncorhynchus nerka* (Sockeye Salmon) were found to significantly increase during spawning events (Tilliotson et al. 2018).

Previous studies have suggested bivalve eDNA shedding rates, and its detection may also fluctuate with environmental and seasonal dynamics. From mesocosm experiments, eDNA was shed at significantly higher rates from mussels collected in September compared to November for *Alasmidonta heterodon* (Dwarf Wedgemussel) in New Jersey, suggesting lower metabolic activity and less DNA release associated with cooler temperatures in autumn (Schill & Galbraith 2019). Additionally, eDNA for the freshwater mussel *Margaritifera margaritifera* (freshwater pearl mussel) was greater in late summer aligning with the release of glochidia compared to samples taken in late spring or early summer during higher flow regimes (Wacker et al. 2019). Environmental DNA concentration for invasive *Corbicula fluminea* (Chinese Basket Clam) was found to be higher during low flow events in summer months compared to higher discharge during autumn months leading to eDNA dilution (Curtis et al. 2020).

Additionally, freshwater mussels are often categorized by their reproductive behavior into two broad groups, as either short-term brooders (tachytictic) or long-term brooders (bradytictic) (Haag 2012). Short-term brooders typically spawn in the late winter and early spring, with females brooding the larvae for a short period (2–8 weeks); whereas long-term brooders typically spawn in the late summer and autumn, with females brooding through the winter (Watters et al. 2009, Haag 2012). These species-specific complex reproductive behaviors are likely to influence the shedding rates of eDNA material, and as a result, freshwater mussel eDNA detections may vary across reproductive seasons (e.g., spring to autumn).

This study assessed the influence of seasonal sampling on the eDNA detection of two diverse freshwater mussel beds in central Ohio streams. Current freshwater mussel protocols for visual tactile searches for surveys supporting regulatory determinations in the state of Ohio restrict freshwater mussel surveys to the period between May and October (ODNR & USFWS 2024), mainly due to an assumption regarding endobenthic burying behavior associated with colder temperatures and a reduced level of metabolic activity. Therefore, eDNA sampling was conducted within a similar sampling period, to assess the detection capabilities of eDNA within a currently prescribed sampling season. This report assesses the ability of eDNA to (1) describe mussel assemblages and estimate relative abundances, (2) detect rare and federally listed species, and (3) detect the extent of a mussel assemblage across a full sampling season.

## Methods

### Site Selection

One sub-objective of this project was to assess eDNA detection of *Epioblasma obliquata* (Purple Cat’s Paw). *Epioblasma obliquata* is a federally endangered species that has been the focus of propagation and reintroduction programs. Specifically, a reintroduction program has occurred in the Walhonding River with propagated mussel releases in 2017, 2018, 2022, and 2023 totaling approximately 1,040 individuals (A. Boyer pers. comm.). Additionally, a population persists within Killbuck Creek (A. Boyer pers. comm.). This study leveraged these two sites as the focus of the eDNA survey (Table 1, Figure 1).

**Figure 1.**
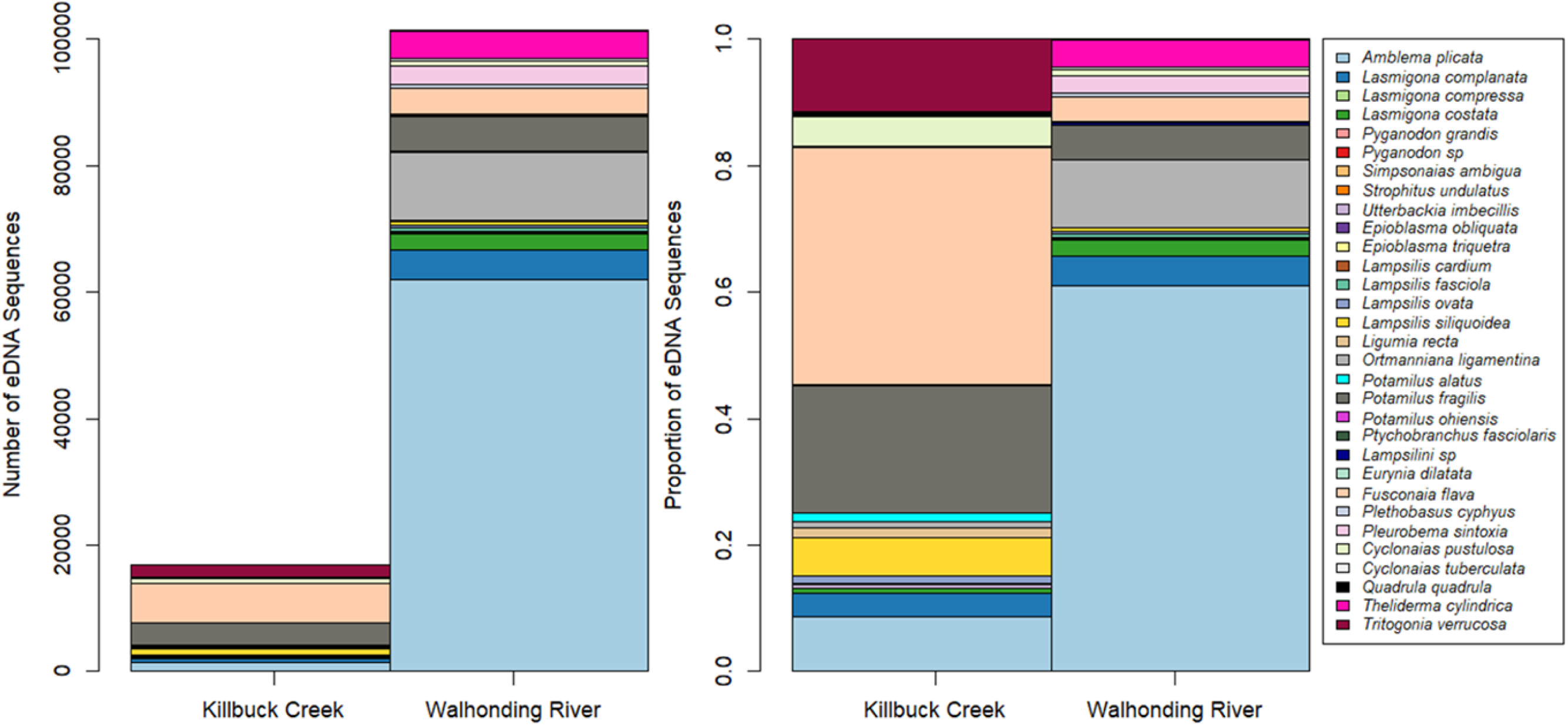
(A) Mean number of freshwater mussel environmental DNA sequences per sample and (B) proportion of freshwater mussel environmental DNA sequences detected in Killbuck Creek and Walhonding River.

**Table 1.** Survey site information for eDNA sampling conducted in Killbuck Creek and Walhonding River.

| **Sampling Site** | **Latitude** | **Longitude** | **County** | **Drainage**<br>**(sq. miles)** | **USGS**<br>**Gage** | **Site Distance to**<br>**Gage** |
| --- | --- | --- | --- | --- | --- | --- |
| Killbuck Cr. | 40.40056 | -81.9575 | Holmes | 582 | 03139000 | 19 KM downstream |
| Walhonding R. | 40.32880 | -81.9896 | Coshocton | 1570 | 03139850 | 9 KM upstream |

In addition to eDNA surveys, a data was obtained from USFWS for a visual assessment at the Walhonding River site from a quantitative tactile search conducted on September 5-6, 2024 (EDGE 2025). This provided an opportunity to compare eDNA and visual search results. In brief, the mussel assemblage at the Walhonding River site was quantitatively assessed where substrates were sampled by randomly placing 400 0.25 m^2^ quadrats across a delineated 566 m^2^ survey area that was previously designated as habitat for *E. obliquata* reintroductions. The 400 quadrats accounted for a total of 100 m^2^, which represented 17% of the delineated survey area. Each of the 400 quadrats was searched at the surface of the sediment with visual/tactile methods, while 266 of the 400 quadrats were further surveyed with excavation of sediment to approximately 10 cm.

### eDNA Sample Collection

A total of ten sampling events occurred at the Killbuck Creek site, and nine total sampling events occurred at the Walhonding River site (Supplementary Table 1). The initial sampling event started on April 25, 2024 (116^th^ Julian Day of Year) in Killbuck Creek and on May 09, 2024 (130^th^ Julian Day of Year) in Walhonding River. The initial sampling date was delayed in Walhonding River as high river discharge and fast flows prevented staff from sampling in April. At both sites, the final sampling date was October 22, 2024 (296^th^ Julian Day of Year) (Supplementary Table 1).

On each sampling date, three replicate water samples were collected at Killbuck Creek and Walhonding River, with the exception of the August 29, 2024 sampling at Walhonding River which consisted of nine total water replicates (Supplementary Table 1). These nine water replicates were collected along the entire perimeter of the central island (see Figure 1). The 10 sampling events in Killbuck Creek resulted in 30 total samples, while the nine sampling events in the Walhonding River resulted in 33 total samples (Supplementary Table 1).

Water samples were collected from near the benthos using a peristaltic pump with a polypropylene filter holder attached to a painter’s pole. A new polypropylene filter holder was used between all samples collected on each day. Environmental DNA was collected along a transect spanning the width of the river for each replicate sample. Once the first sample was collected, the surveyor took the next sample ∼2-3m upstream, and continued this for three replicates per site. Samples were filtered on a 47-mm-diameter GF/C. Each sample consisted of filtering 1000mL of river water. Filters were placed into separate coin envelopes, which were then placed in Ziploc bags with silicone desiccant beads and stored in a freezer.

On each of the 10 days of sampling, a negative field control was collected consisting of 500mL of distilled water being poured into a clean Nalgene in the field and processed along with field samples. Sampling was always conducted in the upstream direction, to avoid potential contamination and sediment disturbance. Field staff used new gloves between each sample, and all equipment washed with 30% bleach following each day of field collection.

### Laboratory Processing

All filters were shipped on ice to Cramer Fish Science Genidaqs laboratory (Sacramento, California, USA, https://genidaqs.com) for DNA extraction and metabarcoding using Illumina MiSeq processing.

The DNA was extracted from each filter using a modified Qiagen DNeasy® Blood and Tissue Kit protocol. Each filter was processed overnight at 56 °C in 540 µl ATL and 60 µl Proteinase K. The resulting supernatant was passed through a QIAshredder spin column, mixed with 600 µl AL and incubated at 70 °C for 10 min. After adding 600 µl ethanol, the resulting mixture was loaded onto a DNeasy Spin column following manufacture’s protocol, with a final elution volume of 100 µl. The DNA was further processed with a Zymo Research One Step PCR Inhibitor Removal kit (Zymo Research, Irvine, California, USA). A negative control was simultaneously extracted to test for possible laboratory contamination.

Each water sample was amplified for a ∼175 base pair (bp) fragment of the mitochondrial 16S gene region which has previously been designed and validated for detection of freshwater mussels from eDNA samples (Prié et al. 2021; Marshall et al. 2022). Mussel eDNA was sequenced with MiSeq Illumina metabarcoding as previously described in Marshall et al. (2022). Library preparation followed a three step PCR described in O’Donnell et al. (2016). Each sample was amplified for a ∼175 bp fragment of the 16S gene region which has previously been tested for amplification of unionid mussels from eDNA samples (Marshall et al. 2022, Prié et al. 2021). Initial PCR amplification was completed for each sample in triplicate with 10 µl PCR reactions containing 4 µl extracted eDNA, 0.4µM primer, and Applied Biosystems™ TaqMan™ Environmental Master Mix 2.0. The amplifications started with an initial denaturation at 95°C for 5 min, followed by 35 cycles of 95°C for 15s, 5% ramp down to 55°C for 30s, and 72°C for 30s. Triplicate PCR products were diluted 1:10 then pooled prior to starting the Illumina adaptor and barcoding PCR processes.

The MiSeq library dual indexed paired-end sample preparation was adapted as described in Miya et al. (2015) from ‘16S metagenomic sequencing library preparation: preparing 16S ribosomal gene amplicon for the Illumina MiSeq system (Illumina part no. 15044223 Rev. B, San Diego, California, USA). A PCR process initiated the incorporation of Illumina adaptors and multiplexing barcodes using Prié et al. (2021) forward and reverse primers containing 33 or 34 base pairs of 5’ Illumina hanging tails to provide a priming site for a final PCR to incorporate barcodes and remaining base pairs of Illumina adaptors. The 12 µl PCR reaction contained 2 µl diluted pooled PCR product, 0.3 µM Illumina adaptor primers and 6 µl 1X Qiagen Plus Multiplex Master Mix. The PCR process denatured for 95°C for 5 min, 5 cycles of 98°C for 20s, 1% ramp down to 65°C for 15s, and 72°C for 15s., followed by 7 cycles of 98°C for 20s, 5% ramp down to 65°C for 15s, and 72°C for 15s. PCR product was diluted 1:10 prior to use in the barcode adaptor PCR process.

The final PCR incorporated paired-end dual indices (eight base pair barcodes) that allowed samples to be identified in the raw read data, and the p5/p7 adaptor sequences to allow the sample to bind onto the Illumina MiSeq flow cell. This final 12μl PCR reaction contained 1μl diluted product from the previous PCR, 0.3 µM forward and reverse indexed primer and 6ul 1X KAPA HiFi HotStart Ready Mix PCR Kit (Roche Diagnostics, Indianapolis, Indiana, USA). Conditions were 3 minutes of initial denaturation at 95°C, followed by 10 cycles at 98°C for 20 s, 5% ramp down to 72°C for 15 s, with a final 5 min 72°C extension. All PCRs were completed on Bio-Rad C1000 Touch Thermal Cyclers. Illumina adapted PCR products were pooled with equal volumes, then size selected (target ∼319bp) using 2% agarose gel electrophoresis. The final pool was sequenced with 2× 300 nt V3 Illumina MiSeq chemistry by loading 6.4 pmol library. An additional 20% PhiX DNA spike-in control was added to improve data quality of low base pair diversity samples. Additionally, a PCR no-template negative control was run for each library preparation step.

#### Bioinformatic Processing and Taxonomic Identification

The data was processed following a bioinformatic pipeline previously described in Marshall et al. (2022). The forward and reverse primer sequences were removed from the demultiplexed sequences using the cutadapt (Martin et al. 2011) plugin within QIIME 2 (Bolyen et al. 2019). Next, sequence reads were filtered and trimmed using the denoising DADA2 (Callahan et al. 2016) plugin within QIIME 2. Based on the quality scores from the forward and reverse read files, a “truncLen” was set to 120 for the forward and 110 for the reverse read files. Using DADA2, error rates were estimated, sequences were merged and dereplicated, and any erroneous or chimeric sequences were removed. Unique sequences were then clustered into Molecular Operational Taxonomic Units (MOTUs) using the QIIME 2 vsearch de-novo with a 97.5% similarity threshold (Marshall et al. 2022). MOTUs from unionid taxa were identified to the species-level using the Basic Local Alignment Search Tool (BLAST+, https://blast.ncbi.nlm.nih.gov/Blast.cgi; Camacho et al. 2009) against our custom database of both in-lab generated sequences and mt-16S sequences downloaded from NCBI GenBank. These MOTUs were further validated with comparisons against the complete NCBI nr database, to investigate alignment to mis-labeled sequences or species not historically within the sampling region. MOTUs that did not return a sequence match from the BLAST search were excluded, as they were considered not from unionid taxa.

Sequences were assigned to a species if they met a threshold of >97.5% identity and 100% query coverage. Furthermore, sequences assigned to multiple species with the same BLAST e-value score were inspected and a final decision was made based on known species distribution and presence within the sampled drainages. Additionally, if multiple sequences were assigned to the same taxonomy, they were inspected and removed or collapsed into a single MOTU to obtain a final matrix of read counts per taxa. Freshwater mussels display a unique form of mitochondrial inheritance, termed doubly uniparental inheritance (DUI), in which males possess a paternal mitochondrial mitotype that is restricted to male gonads and gametes (Gusman et al. 2016). As the male mitotype is genetically distinct from the female mitotype (Curole & Kocher 2005), we only retained sequences determined to be the female mitotype for direct evaluation of eDNA detection probabilities and seasonal assemblage assessments.

In metabarcoding analysis, the sequencing process can introduce a form of sample cross-contamination in which sequences from one sample are falsely detected within another sample, often termed ‘critical mistags’, ‘tag jumps’, or ‘index hopping’ (Esling et al. 2015, Richardson 2022). These sequencing artifacts generally represent a small fraction of the total sequences (Esling et al. 2015), yet they can lead to false detections in some circumstances. Therefore, the final processing of the MOTUs in this study consisted of estimating and removing mistags from the dataset following the framework outlined by Richardson (2022). Based on the number of reads per MOTU within the field and laboratory negative controls, the mistag rate was estimated for all dual-index combinations. Then this mistag rate was applied to all field samples to identify and remove potential cases of erroneous detection.

### Data Analysis

#### eDNA Detection Index

To assess the repeatability of eDNA detection, an eDNA Detection Index was developed and applied to all species detections at both sites within each of the sampling events. The index levels are described as:

- **<u>Non-detection</u>** occurs when a species is not detected in any water sample replicates collected within a single site.
- **<u>Low repeatability</u>** occurs when a species is detected in **≤ 33%** of water replicates collected within a single site.
- **<u>Moderate repeatability</u>** occurs when a species is detected in **≤ 67% and > 33%** of water replicates collected within a single site.
- **<u>High repeatability</u>** occurs when a species is detected in **> 67%** of water replicates collected within a single site.

#### Mussel Assemblage

Community assemblages were visualized with bubble plots to show the amount of eDNA detected for each species (i.e., measured as the number of sequence reads) on each sampling event from the two sites. For the bubble plots, the amount of eDNA detected for each species on each sampling event was averaged across the water replicate samples collected at that site and on that date. The eDNA Detection Index was used within the bubble plots to illustrate detection across replicate water samples.

Next, the amount of eDNA per sampling event was normalized by calculating the relative abundance (i.e., species proportion) on each sampling date. The relative abundances were calculated using the mean amount of eDNA for each species across the water replicate samples divided by the total unioinid sequences. Barplots were used to visualize the assemblage proportions across the sampling season within each of the two sites.

Non-metric multidimensional scaling was used to assess similarity in species detections across water sample replicates. Non-phylogenetic beta diversities were estimated using (a) a qualitative metric (calculated from presence/absence) – Jaccard dissimilarity) and (b) a quantitative metric (calculated from the assemblage proportions – Bray-Curtis Dissimilarity). These beta diversity metrics were used to assess the amount of variation in the detected mussel community across the sampling season. PerMANOVAs (vegan function adonis2) were used to test for differences in the amount of seasonal variation within Killbuck Creek replicates compared to the amount of seasonal variation within Walhonding River replicates. With homogeneity of multivariate dispersions calculated using the betadisper function and comparisons statistically tested with ANOVA.

#### Seasonal Trends

The amount of eDNA detected for each species at the two sites was plotted across sampling dates. The seasonal trend in the amount of eDNA and the proportion of mussel assemblage for each species was assessed using a regression spline in R with the *ss* function. The amount of eDNA for each species on each sampling date was plotted against the gage height of the nearest USGS gage (see Table 1).

Patterns in species richness and community assemblage (diversity metrics) were analyzed to assess the influence of discharge and sampling date (i.e., Julian Day). Because both USGS gage stations do not record discharge directly (flow cfs), the flow exceedance probability was used as a surrogate of discharge. A measure of flow exceedance was generated by calculating the exceedance probability on each sampling data, where the exceedance probability is the total number of days in which the gage height was higher than that on the sampling date divided by the total number of days on record for that gage. This flow exceedance probability was calculated using daily average values and using the entire dataset of record. At both sites, the flow exceedance decreased across the sampling period (Supplementary Figure 1). For example, initial sampling in April at Killbuck Creek occurred at a flow exceedance as high as 0.67 and the final sampling in October occurred at a flow exceedance of only 0.02, with similar trends observed in Walhonding River (Supplementary Figure 1). Therefore, a strong correlation between time of year and flow exceedance made it not possible to assess eDNA detection patterns in response to these two factors.

#### Probability of eDNA Detection

To assess how eDNA concentration influences detection repeatability, the repeatability of eDNA detection (i.e., eDNA Detection Index) was compared to the amount of eDNA collected for each detection.

To estimate detection probability and to assess the eDNA survey level of effort, Bayesian integrated occupancy models were developed in the ‘spOccupancy’ R package (Doser et al. 2022). To accomplish this, the eDNA relative-abundance data was transformed into binary detection/non-detection data for all species detected with eDNA across the two sites. Occupancy models leverage repeated observations of detection/non-detection data to jointly estimate detectability and probability of occurrence (MacKenzie et al., 2002). A multispecies occupancy model was evaluated for all species to estimate the mean detection probability (p) (i.e., the probability of successful eDNA detection of a species within a replicate environmental sample). The survey design was evaluated by calculating the detection probability (p*) for each species: p* = 1-(1-p)^n^ where p is the estimated detection for a single replicate environmental sample and n is the total number of replicates. The detection probabilities for each species were estimated for both sites.

The occupancy models were further evaluated to assess how seasonal sampling (day of the year) or river discharge may have influenced detection probability. To accomplish this, two additional occupancy models were analyzed: (1) using site-level flow exceedances as a detection covariate and (2) using the site-level Julian day of year as a detection covariate. By keeping these detection covariates at the site-level (Walhonding River vs Killbuck Creek), we are able to compare how detection patterns differed between the two sites. To evaluate factors influencing detectability across mussel species, we also included species-level attributes: (1) whether the species was visually observed at the site, (2) short-term versus long-term brooding behavior, and (3) presence or absence of shell sculpturing.

### Comparisons between eDNA and Visual Survey

The mussel assemblage detected in Walhonding River with eDNA on each sampling date was compared to the visually observed mussel assemblage. The mean eDNA abundance across the entire sampling period was compared to the visually observed mussel abundance using linear regression. To assess seasonal influences on the detected mussel assemblage, linear regressions were also used to compare the eDNA abundance on each individual sampling event to the visually observed mussel abundance.

Additionally, beta diversities were estimated using a quantitative metric (calculated from the assemblage proportions– Bray-Curtis Dissimilarity) for each eDNA sampling event and the visual mussel survey.

These beta diversities were compared across time between an eDNA sampling event and the visual survey (measured as days between the two surveys).

## Results

The metabarcoding processing resulted in 10,022,412 raw sequence reads, of which a total of 3,853,519 reads were classified as a unionid female mitotype 16S sequence, with an average of 61,165.1 ± 57,497.4 (standard deviation) unionid female mitotype reads per sample. The total number of species detected in a sample was positively correlated with the number of sequences obtained in the sample (Supplementary Figure 2).

Across the 10 field controls and the four laboratory controls, the number of female mussel sequences ranged from 0 to 103 total reads per sample. A total of five MOTUs occurred within the control samples, ranging from 3 to 103 total sequence reads per MOTU. The control samples consisted of 13 total occurrences of a MOTU (mean reads per occurrence = 19.5 ± 28.2 std). The mistag analysis removed all but one occurrence from the control samples. The mistag analysis removed a total of 10 MOTU occurrences from the 63 field eDNA samples.

### Species Detection

In total, 33 MOTUs were classified as unionid female mitotype. *Quadrula quadrula* (Mapleleaf) and *Ortmanniana ligamentina* (Mucket) each had two MOTUs with greater than 97% genetic identity to a reference sequence, and thus these MOTUs were considered intraspecific population variation and were ultimately consolidated as a single *Q. quadrula* MOTU and a single *O. ligamentina* MOTU for further reporting. This reduced the number of MOTUs from 33 to 31 for this analysis. Of these 31 MOTUs, 29 were identified to a species-level, ranging in percent of genetic identity from 99.25 to 100.00 (Table 2).

**Table 2.** Species detected and their environmental DNA sequence abundance from Killbuck Creek and Walhonding River. The visual mussel abundance is listed for Walhonding River.

| Tribe | Species | Common Name | Percent Genetic Identity | Killbuck Cr. | Walhonding R. |  |
| --- | --- | --- | --- | --- | --- | --- |
|  |  |  |  | eDNA | eDNA | Visual |
| Amblemini | *Amblema plicata* | Threeridge | 100 | 44288 | 2043499 | 600 |
| Anodontini | *Lasmigona complanata* | White Heelsplitter | 100 | 17882 | 153488 | 11 |
|  | *Lasmigona compressa* | Creek Heelsplitter | 100 | 324 | 949 | 0 |
|  | *Lasmigona costata* | Flutedshell | 100 | 3663 | 86263 | 24 |
|  | *Pyganodon grandis* | Giant Floater | 100 | 252 | 0 | 0 |
|  | *Pyganodon sp.* | - | 100 | 22 | 57 | - |
|  | *Simpsonaias ambigua*<sup>PFE</sup> | Salamander Mussel | 100 | 0 | 240 | 0 |
|  | *Strophitus undulatus* | Creeper | 100 | 91 | 355 | 2 |
|  | *Utterbackia imbecillis* | Paper Pondshell | 100 | 2919 | 2999 | 0 |
| Lampsilini | *Epioblasma obliquata*<sup>FE</sup> | Purple Cataspaw | 100 | 494 | 171 | 16 |
|  | *Epioblasma triquetra*<sup>FE</sup> | Snuffbox | 100 | 103 | 0 | 0 |
|  | *Lampsilis cardium* | Plain Pocketbook | 100 | 351 | 7861 | 9* |
|  | *Lampsilis fasciola* | Fatmucket | 100 | 0 | 19404 | 5 |
|  | *Lampsilis ovata* | Pocketbook | 99.25 | 5744 | 13329 | 0* |
|  | *Lampsilis siliquoidea* | Wavy Rayed Lampmussel | 100 | 30733 | 21655 | 1 |
|  | *Ligumia recta* | Black Sandshell | 100 | 7710 | 2263 | 0 |
|  | *Obliquaria reflexa* | Threehorn Wartyback | - | 0 | 0 | 1 |
|  | *Ortmanniana ligamentina* | Mucket | 100 | 5006 | 355546 | 259 |
|  | *Potamilus alatus* | Pink Heelsplitter | 100 | 6866 | 3622 | 0 |
|  | *Potamilus fragilis* | Fragile Papershell | 100 | 101855 | 181010 | 11 |
|  | *Potamilus ohioensis* | Pink Papershell | 100 | 0 | 431 | 0 |
|  | *Ptychobranhus fasciolaris* | Kidneyshell | 100 | 141 | 323 | 0 |
|  | *Lampsilini sp.* | - | 90.51 | 203 | 10244 | - |
| Pleurobemini | *Eurynia dilatata* | Spike | 99.25 | 131 | 7477 | 0 |
|  | *Fusconaia flava* | Wabash Pigtoe | 100 | 189671 | 133050 | 53 |
|  | *Plethobasus cyphus*<sup>FE</sup> | Sheepnose | 100 | 0 | 17568 | 11 |
|  | *Pleurobema sintoxia* | Round Pigtoe | 100 | 921 | 94788 | 69 |
| Quadrulini | *Cyclonaias pustulosa* | Pimpleback | 100 | 23698 | 30151 | 45 |
|  | *Cyclonaias tuberculata* | Purple Wartyback | 100 | 416 | 12495 | 21 |
|  | *Quadrula quadrula* | Mapleleaf | 100 | 2943 | 64 | 0 |
|  | *Theliderma cylindrica*<sup>FT</sup> | Rabbitsfoot | 100 | 0 | 141706 | 28 |
|  | *Tritogonia verrucosa* | Pistolgrip | 100 | 58425 | 7659 | 46 |

PFE = Proposed Federally Endangered, FE = Federally Endangered, FT = Federally Threatened
\*Morphological distinctions between *Lampsilis cardium* and *L. ovata* are very difficult, and previous eDNA sampling has identified both species present within the Walhonding River. Further comparisons between eDNA and visual data combine these two species.

One MOTU was only identified to the genus level as *Pyganodon* sp. (Table 2). While the scientific species name is incomplete, this sequence is 100% genetic identity to a previously discovered, but unnamed, cryptic *Pyganodon* from the Great Lakes region in northeastern North America (Cyr et al., 2007). This sequence was previously detected in the Walhonding River during eDNA sampling at the Six Mile dam (Marshall et al. 2022). The other MOTU not identified to the species level was classified only to the tribe level as Lampsilini sp. (Table 2). While this sequence does group within the Lampsilini tribe (Supplementary Figure 3), it is less than 91% genetic identity to any known genetic sequence, and thus cannot definitively be identified to the species or genus level.

A total of five federally listed and/or proposed for listing species were detected. This included federally endangered *Epioblasma obliquata*, *E. triquetra* (Snuffbox), and *Plethobasus cyphyus* (Sheepnose), federally threatened *Theliderma cylindrica* (Rabbitsfoot), and proposed for listing *Simpsonaias ambigua* (Salamander Mussel) (Table 2). *Epioblasma obliquata* was detected at both sites, while *E. triquetra* was detected at Killbuck Creek only and *P. cyphyus, T. cylindrica, S, ambigua* were detected at Walhonding River only. Both *Epioblasma* spp. had relatively low sequence reads across the study, with *E. triquetra* and *E. obliquata* eDNA accounting for only 0.02% (103 of 504,852 sequence reads) and 0.1% (494 of 504,852 sequence reads) of all mussel eDNA in Killbuck Creek and *E. obliquata* eDNA accounting for only 0.005% (171 of 3,348,667 sequence reads) of all mussel eDNA in Walhonding River (Table 2).

Of the 31 MOTU sequences detected across the study, 26 were detected within Killbuck Creek and 29 were detected within Walhonding River (Table 2). The number of sequence reads pre MOTU greatly varied across species, ranging from 22–189,671 in Killbuck Creek and 57–2,043,499 in Walhonding River (Table 2). The majority of the eDNA sequence reads were attributed to samples collected at Walhonding River (3,348,667, 87% of total sequence reads) compared to Killbuck Creek (504,852, 13% of total sequence reads) (Figure 2). The species assemblage was relatively different between the two sites, with Killbuck Creek dominated by *Fusconaia flava* (Wabash Pigtoe) and *Potamilus fragilis* (Fragile Papershell), while Walhonding River was dominated by *Amblema plicata* (Threeridge) (Table 2, Figure 2).

**Figure 2.**
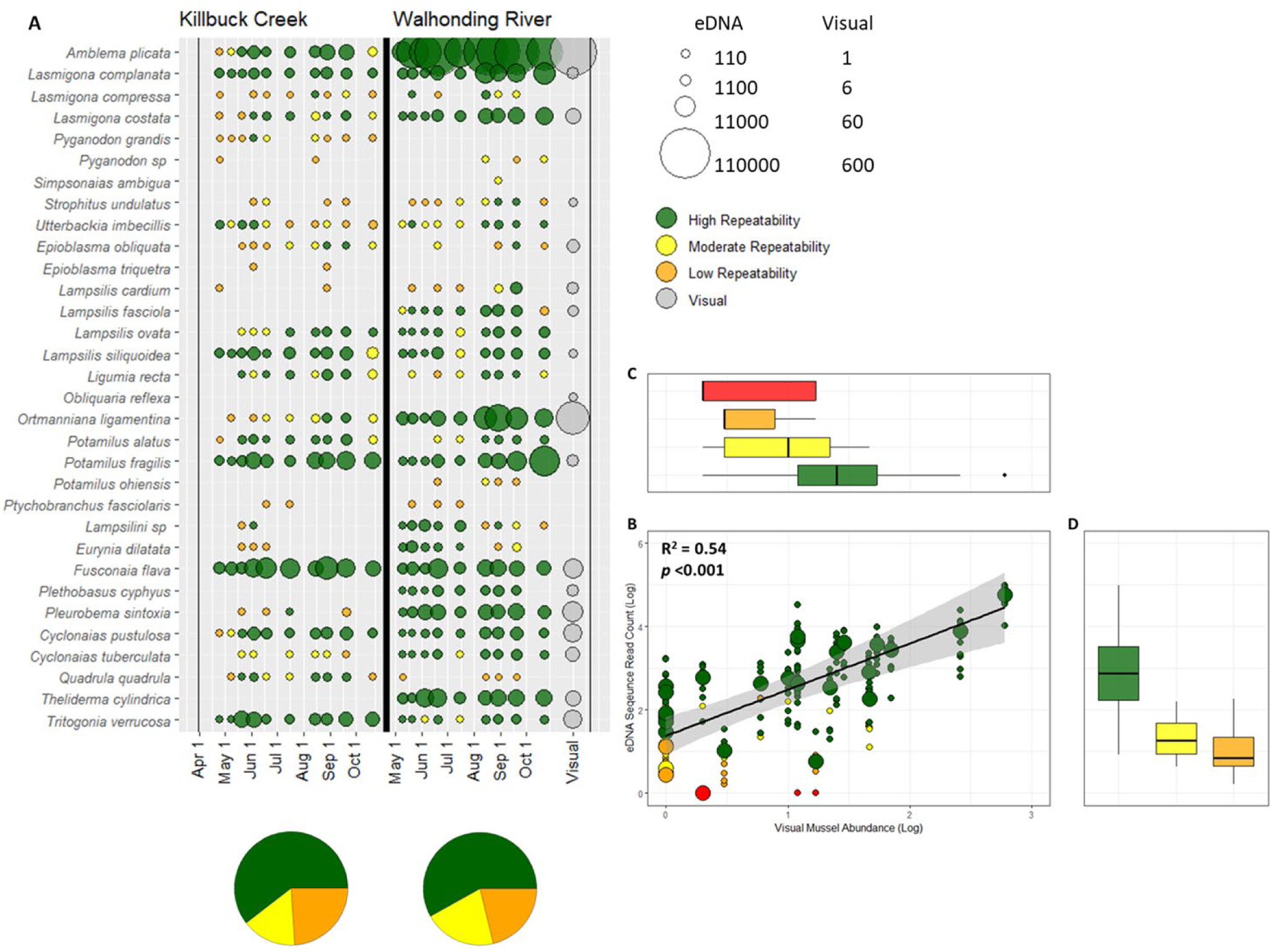
(A) Bubble plot of seasonal environmental DNA detections for freshwater mussels in Killbuck Creek and Walhonding River. Size of bubbles represent the mean eDNA sequence and colors represent repeatability of detection across replicates. Visual mussel abundance found in the Walhonding River is shown in grey. (B) Linear regression comparing the mean environmental DNA abundance (i.e., sequence read count) across all sampling events to the visually observed mussel abundance. Large circles represent the mean eDNA abundance, while small circles represent the eDNA abundance on each individual sampling event. Boxplots represent variation in (C) visual mussel abundance or (D) eDNA sequence read count across the eDNA detection index

#### eDNA Detection Index

The majority of detections in both Killbuck Creek and Walhonding River were classified as high repeatability (Figure 3). A greater number of high repeatability detections occurred in Walhonding River (156, 74% of detections) than Killbuck Creek (102, 56% of detections) (Figure 3). There were significant differences between the amount of DNA sequence reads obtained at the three levels of the eDNA detection index, where detections occurring at high repeatability had on average the greatest amount of sequence reads, and detections at moderate and low repeatability had on average similar amounts of sequence reads (ANOVA: F-value = 201.6, *p* < 0.001) (Figure 3).

**Figure 3.**
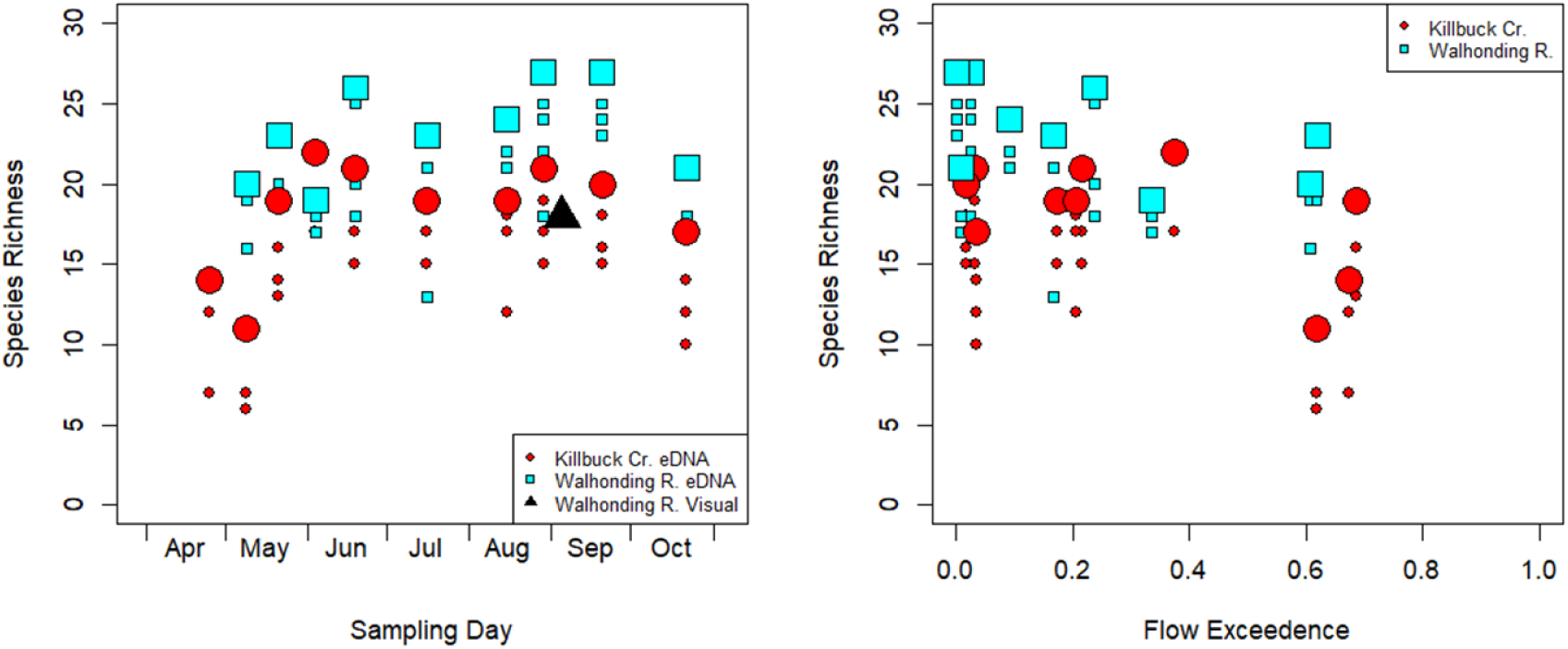
Species richness detected with eDNA within Killbuck Creek and Walhonding River (A) across the sampling time points and (B) across flow exceedance at time of sampling. Large symbols represent the total richness detected on that sampling date and small symbols represent the richness detected within each eDNA sample replicate.

### Mussel Assemblage and Sampling Season

The detected species richness with eDNA varied across the sampling time points, with Killbuck Creek ranging from 11 to 21 species and Walhonding River ranging from 20 to 27 species (Figure 4). Generally, the species richness increased across the sampling time points and decreased with increasing flow exceedance (Figure 4). For example, the two lowest richness time points in Killbuck Creek occurred in the first two sampling events in April and May, coinciding with the greatest flow exceedance (Figure 4).

**Figure 4.**
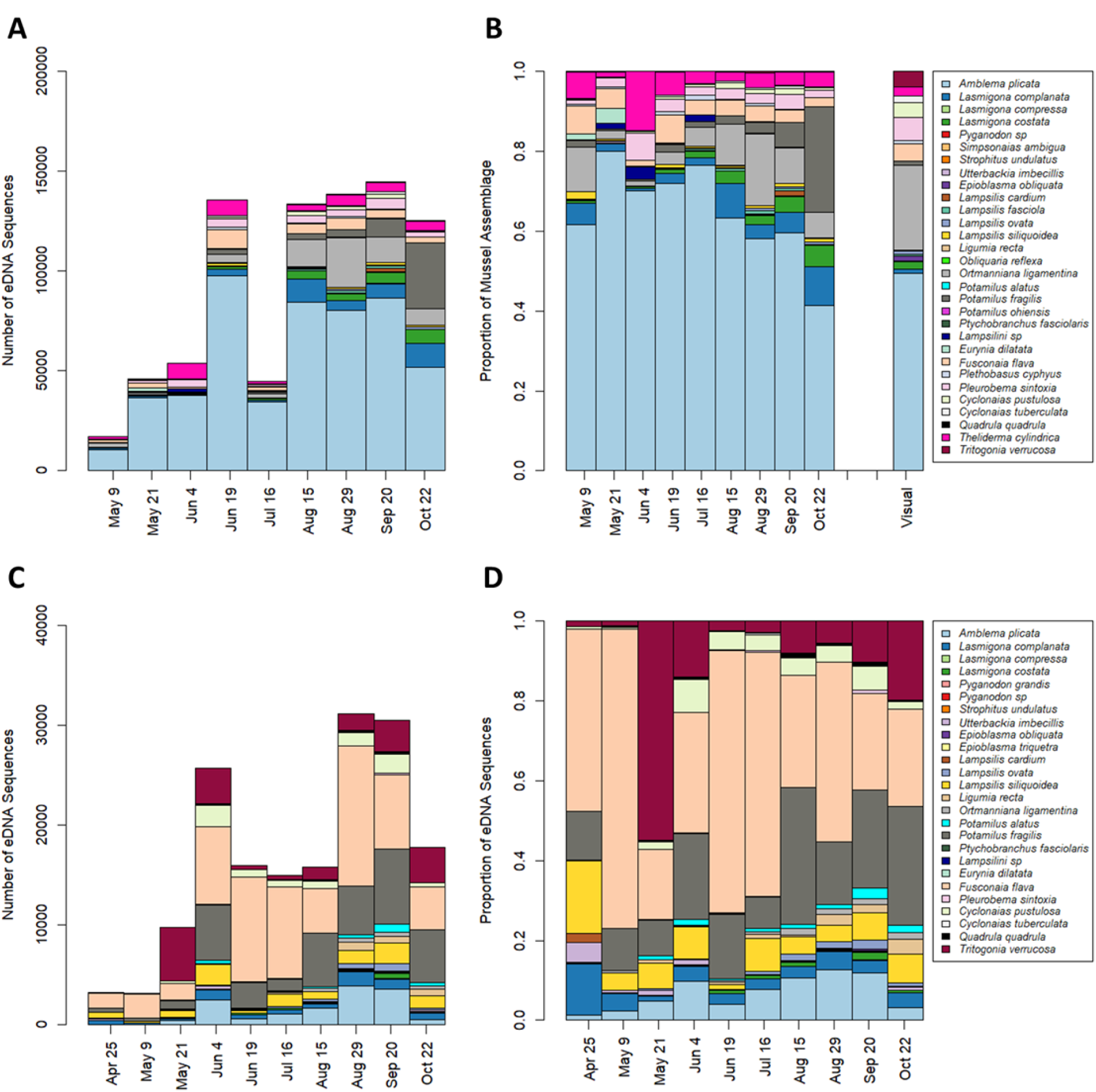
(A) Number of freshwater mussel environmental DNA sequences and (B) proportion of freshwater mussel environmental DNA sequences detected across sampling events in Walhonding River. (C) Number of freshwater mussel environmental DNA sequences and (D) proportion of freshwater mussel environmental DNA sequences detected across sampling events in Killbuck Creek. The proportion of freshwater mussel abundance recorded during a visual survey in Walhonding River is shown in B.

The relative abundance estimates from eDNA were relatively constant across the sampling season at both sites, with a few exceptions (Figure 5). Seasonal trends in sequence read abundance can be found in Supplementary Figure 4 & 5. Community assemblage displayed generally lower fluctuations in Walhonding River (Figure 5, Supplementary Figure 6). Notably, within Killbuck Creek, *Tritogonia verrucosa* (Pistolgrip) displayed the largest fluctuation, with a particularly high spike occurring on May 21, 2024 (Figure 5).

**Figure 5.**
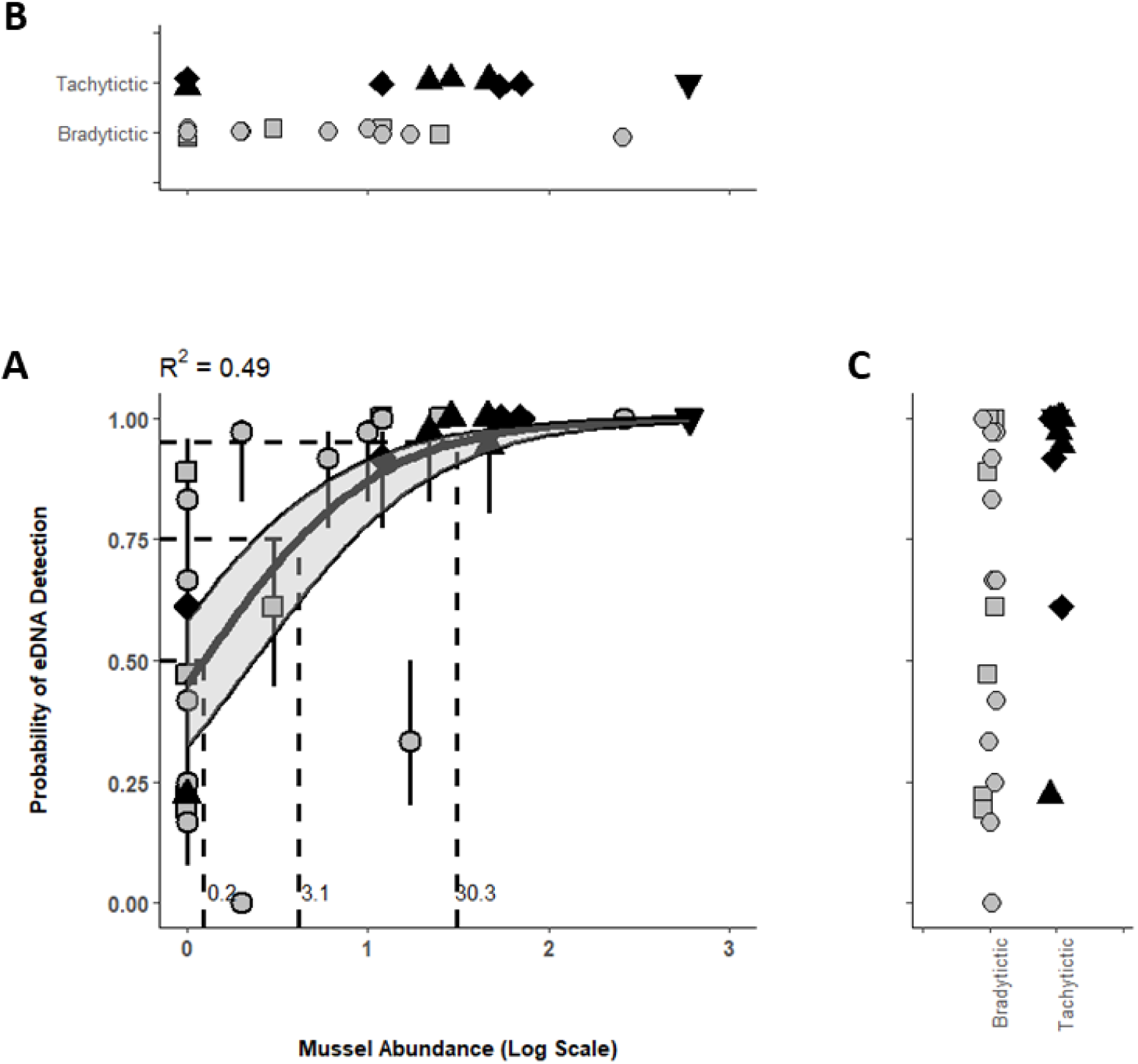
(A) Logistic regression model depicting how mussel abundance influences eDNA probability of detection. Individual error bars represent credibility intervals for probability of detection estimates. Thick regression curve represents median probability of detection estimates, while grey shading represents logistic regression from upper and lower probability of detection estimates. The dashed lines indicate the threshold abundance for detection at 0.50, 0.75, and 0.95 detection probabilities. Scatterplots depict variation in (B) mussel abundance or (C) eDNA detection probability based on reproductive behavior (tachytictic (black) or bradytictic (grey)) and mussel tribe (Amblemini (upside down triangle), Anodontini (square), Lampsilini (circle), Pleurobemini (triangle), Quadrulini (diamond)).

Beta diversity estimates (i.e., Bray Curtis Dissimilarity and Jaccard Dissimilarity) and NMDS analysis were used to further assess fluctuations in community assemblage over the sampling period. Patterns within both beta diversity metrics were similar, where the samples clustered within their respective sampling site (i.e., Killbuck Creek vs Walhonding River) (Supplementary Figure 6, circles vs squares). Additionally, samples showed low dissimilarity across sampling replicates collected on the same sampling date (Supplementary Figures 6, 7, & 8). To further assess seasonal fluctuations in the detected assemblage, the beta diversity metrics were assessed against the amount of time between the sampling events. In both Killbuck Creek and Walhonding River, the community dissimilarity increased with increasing number of days between sampling events (Supplementary Figure 8).

Species detected in Walhonding River generally displayed a greater probability of eDNA detection than those detected in Killbuck Creek (Walhonding River mean POD = 0.73; Killbuck Creek mean POD = 0.62) (Supplementary Figure 9). In Killbuck Creek, species-specific detection probabilities were more frequently negatively influenced by river discharge than by day of year. Discharge negatively affected detection for 15 species, whereas only six species showed negative effects of day of year (Supplementary Figures 10-13). In contrast, detection probabilities in the Walhonding River were relatively stable across both covariates, with only one species negatively affected by discharge and three species negatively influenced by day of year (Supplementary Figures 10-13).

### Comparison of eDNA and Visual Surveys

The eDNA data collected in the Walhonding River was compared to that collected from quadrat surveys by EDGE on September 5-6, 2024 (EDGE 2025). The visual survey in Walhonding River observed 18 species, while the eDNA surveys detected 28 MOTUs with 26 of those being identified to a species (Table 2, Figure 3). Across all sampling timepoints, eDNA detected greater species richness than that observed with the visual survey (Figure 4). Species found to be abundant during the visual survey were typically detected as a large proportion of the mussel eDNA (e.g., *A. plicata* and *O. ligamentina*) (Figures 3 & 5B). Of the 18 species observed during visual surveys, 16 were detected on at least eight of the nine sampling occurrences (Table 2, Figure 3). *Obliquaria reflexa* (Threehorn Wartyback) was observed as a single individual and was never detected with eDNA, while *E. obliquata* was observed as 16 individuals and detected with eDNA on four occurrences (Table 2, Figure 3).

Species that were observed with the visual survey were typically detected across all eDNA sampling events, regardless of sampling season or flow exceedance (Supplementary Figure 14, Table 2, Figure 3). However, for species detected with only eDNA (i.e., species not observed during the visual survey), the number of eDNA-only species increased across the sampling season and with decreasing flow exceedance (Supplementary Figure 14, Figure 3). For example, during high flow exceedance in the spring, only three or four eDNA-only species were detected, compared to 10 eDNA-only species detected during low flow exceedance in late summer (Supplementary Figure 14).

The mean eDNA abundance across all sampling events (i.e., the mean sequence read count across all samples) was positively correlated with the observed visual abundance (R^2^ = 0.54, *p* < 0.001) (Figure 3). Species observed with the visual survey were likely to be detected with eDNA at high repeatability (Figure 3). Similarly, the eDNA sequence abundance increased as the repeatability of eDNA detection increased (Figure 3).

While the mean correlation between eDNA abundance and visual abundance was R^2^ = 0.54, this correlation varied across the nine sampling events, ranging from 0.22 – 0.61 (Supplementary Figure 15). The highest correlation was found when the eDNA survey was conducted in close temporal proximity to the visual survey (i.e., within ±20 days of the visual survey; Supplementary Figure 16). Similarly, comparisons of community assemblage between eDNA and the visual survey found that the best congruence between survey methods (measured as Bray-Curtis Dissimilarity) occurred when the eDNA survey was conducted in close temporal proximity to the visual survey (i.e., within ±20 days of the visual survey; Supplementary Figure 17).

Logistic regression assessed the probability of eDNA detection compared to the visually observed mussel abundance, with detection probability increasing with mussel abundance (Figure 6A). Mussels that are broadly categorized as tachytictic generally had a higher probability of eDNA detection than mussels categorized as bradytictic; however, tachytictic mussels were generally more abundant than bradytictic mussels (Figure 6B &C). Species detectability was generally higher when a mussel species was visually confirmed at the sampling site and when the species possessed a sculptured shell, whereas brooding strategy (short-term vs. long-term) showed no clear influence on detection probability (Supplementary Figure 18).

## Discussion

The reproductive behaviors of a species can significantly alter their seasonal DNA shedding rates and ultimately influence eDNA detections across reproductive and life history stages (de Souza et al. 2016, Tilliotson et al. 2018, Troth et al. 2021). This is likely true for freshwater mussels, which display seasonal changes in vertical migration within the sediment and behaviors associated with spawning, glochidia release, and host parasitization (Amyot & Downing 1991, Watters et al. 2001, Sanchez & Schwalb 2019). Yet, this study consistently detected mussel eDNA from April through September from diverse mussel beds that consisted of species grouped into the two broad reproductive strategies for freshwater mussels (Haag 2012). Broadly speaking, tachytictic mussels, such as those in the Pleurobemini and Quadrulini tribes, are expected to have short term brooding behaviors with spawning occurring in early spring and glochidia release in late summer or early autumn. Whereas bradytictic mussels, such as those in the Anodontini and Lampsilini tribes, are expected to have long term brooding behaviors with spawning occurring in late summer or early autumn and glochidia release in early to late spring (Watters et al. 2009, Haag 2012). Even with these diverse reproductive behaviors, the mussels known to occupy the Walhonding River site were consistently detected with eDNA across sampling events.

However, the species richness detected in Killbuck Creek ranged from 11 to 21 species depending on the sampling date, with the lowest richness coinciding with low temperatures and high flow conditions in April and May. Many mussel species exhibit endobenthic behavior during winter and may not emerge from burying until sometime in early to mid-spring (Watters et al. 2001, Amyot & Downing 1997). These seasonal endobenthic burying behaviors may limit detection when sampling occurs prior to spring re-emergence. Additionally, high discharge events can confound eDNA results by diluting eDNA and reducing detection rates (Curtis et al. 2020). The sampling dates with the lowest species richness occurred during discharge exceeding 60% of historically flows in Killbuck Creek, suggesting high flows may have reduced detection efficiency. However, the true mussel assemblage present at the Killbuck Creek site is unknown, and thus it is not possible to fully assess seasonal eDNA detection trends of the local fauna. Nonetheless, these results do suggest caution should be taken about sampling eDNA in early spring during high discharge events or before mussel re-emergence from the benthos.

A seasonal assessment of vertical migration of eight species in Ohio suggested mussels may follow two distinct migration patterns (Watters et al. 2001). One group displayed a unimodal annual pattern, with most of the population coming to the surface in the spring and remaining there until late autumn. Whereas the second group displayed a bimodal pattern, with the population coming to the surface in the spring, re-burying in mid-summer before re-emerging at the surface in early autumn and remaining until late autumn. Three of the previously studied bimodal species were detected with eDNA in the current study (*A. plicata*, *C. pustulosa*, and *F. flava*), with all three species being detected on all sampling dates from both study sites. Thus, the summer burial behavior previously observed in these species did not appear to affect seasonal eDNA detection, although it remains uncertain whether the current studied populations followed the same vertical migration patterns described by Watters et al. (2001).

Although most visually observed species in the Walhonding River were routinely detected with eDNA, an exception to this occurred for *O. reflexa* and *E. obliquata*. *Obliquaria reflexa* was visually observed as only a single individual and was subsequently never detected on any sampling date with eDNA. This species was observed as the least abundant, and thus the collection of three eDNA samples per sampling event may not have been sufficient to detect a species this rare (Marshall & Fleece 2025). However, the next three least abundant species (*L. siliquoidea* (n = 1 individual), *S. undulatus* (n = 2 individuals), and *L. fasciola* (n = 5 individuals)) were all detected with eDNA on at least eight of the nine sampling events in the Walhonding River. Therefore, similarly rare species were consistently detectable with the current level of survey effort.

Conversely, *E. obliquata* was more abundant (n = 16 individuals) but only detected on four of the nine sampling events, and only once occurred as a high repeatability detection. *Epioblasma* species often exhibit endobenthic behavior, which might limit their detection with eDNA. For example, Smith et al. (2001) found only 52% of *Epioblasma torulosa rangiana* (Northern Riffleshell) were recovered from the substrate surface, whereas most species were recovered at >70% at the substrate surface. Similarly, the quadrat excavation survey conducted at the site recovered 75% of the *E. obliquata* from the subsurface (EDGE 2025), suggesting this species was more likely to be buried than at the surface. Furthermore, *E. obliquata* was an outlier within the abundance based eDNA detection model, with a detection probability lower than expected based on its abundance.

Another *Epioblasma*, *E. triquetra,* had the least number of detections out of all the species in Killbuck Creek. However, while extant populations of *E. triquetra* occur within the system, it is unknown the density and proximity to the current eDNA sampling site, and thus it is difficult to ascertain if low eDNA detection rates are a result of low mussel density, far proximity to sample collection, or seasonal burying behavior. *Epioblasma obliquata* was estimated to have nearly double the detection probability within Killbuck Creek compared to Walhonding River, where it was detected on nearly every sampling event in Killbuck Creek. It is unknown how the species density in Killbuck Creek compares to Walhonding River, and how that may have influenced detection patterns between the two sites.

Nevertheless, at both sites, the eDNA sequence abundance of *Epioblasma* spp. was consistently low across sampling events, never accounting for greater than 1% of the total mussel DNA. This may suggest that DNA is less likely to transfer to the water column for endobenthic species such as those in the *Epioblasma* genus. However, relatively high detection rates for *E. rangiana* and *E. triquetra* have previously been recorded (Waits et al. 2025). The sampling season in this previous work was not reported, and thus it is not possible to put it into context of seasonal influences on detection for these species. *Epioblasma triquetra* was found to have varying eDNA detection estimates across sampling sites (Waits et al. 2025), suggesting site-specific abundance likely influences species detection patterns.

In the current study, the estimates for probability of eDNA detection were correlated with abundances, similar to previous studies where eDNA detection increases as a species becomes more abundant (Marshall et al. 2022, Marshall & Fleece 2025). Most species observed during the visual surveys were detected with high repeatability and were estimated to have eDNA detection probabilities exceeding 0.90. However, mussel surveys are often implemented to assess the presence/ probable absence of rare and federally listed species, and therefore eDNA needs to be assessed for its ability to detect rare species at a site. In the current study, eDNA generally succeeded at detecting rare species, with four of the five species visually observed as less than 10 individuals being detected on at least eight sampling events and typically detected at high repeatability. Sampling greater than three samples per sampling event would additionally increase the number of detection events for rare species (Marshall & Fleece 2025).

Abundance assessments are difficult to obtain from an eDNA metabarcoding survey due to differences in taxa-specific eDNA shedding rates (e.g., differences in size and surface area, life-histories, spawning times, metabolic activity, biological behavior), habitat differences, and PCR-based biases (e.g., differential primer annealing and amplification) (Ruppert et al. 2019). However, it has been shown that eDNA sequence read counts generally correlate to the relative abundances of the mussels at a site (Marshall et al. 2022, Marshall & Fleece 2025). In the current study, eDNA had the highest congruence with the visual survey when sampled in close temporal proximity (within ± 20 days) and during low discharges. As we are unable to differentiate seasonality from discharge, the higher congruence between eDNA and the visual survey during the same sampling month may be explained by a combination of factors, such as (1) lower dilution and transport effects occurred during low discharge and/or (2) temporal shedding rates accurately corresponded to the active epibenthic mussel community that was readily recovered by tactile searchers.

In addition to abundance, the multispecies occupancy model at the Walhonding River revealed that two species-level attributes strongly influenced detection probability: the presence of shell sculpturing and whether a species was visually observed at the site. Species with sculptured shells had consistently higher detection probabilities than unsculptured species, potentially suggesting sculptured species are more likely to be at the sediment surface where eDNA can be released into the water column. Yet the finding that species that were visually confirmed during tactile surveys were more likely to be detected with eDNA suggests that true local presence strongly increases eDNA detection likelihood, even for species with low abundance. These findings highlight that morphological traits and confirmed site occupancy can meaningfully shape species-specific eDNA detection patterns.

Generally, the total eDNA sequence abundance and the detected species richness was lowest during spring sampling at both sites, coinciding with the highest flow regimes. While the potential transport distance of eDNA typically increases with river discharge (Wilcox et al. 2016, Van Driessche et al. 2022), and bivalve eDNA has been reported to transport several kilometers in some river systems (Deiner & Altermatt 2014, Shogren et al. 2019, Stoeckle et al. 2021), the current study found high discharges reduced species richness. The effective source area (i.e., the stretch of the river contributing to the eDNA pool) is likely to be greater during high flow conditions, however, these high flow conditions may also dilute eDNA signal from rare species (Curtis et al. 2020). High flows generally resulted in lower detection repeatability for many species, and therefore, surveys conducted in early spring likely require greater sampling effort for detection of rare mussels.

The detection repeatability of eDNA may be useful for interpreting a species’ proximity to the sampling location. For example, majority of eDNA detections occurring at high repeatability in the Walhonding River were positively confirmed with visual observations. Whereas species not visually observed were sporadically detected across sampling events and typically at low to moderate repeatability. This suggests eDNA detections occurring at high repeatability have high likelihood that their presence would be later confirmed from a visual survey, while those at low to moderate repeatability may be regionally-present but not local to the sampling site. However, low to moderate repeatability detections sometimes do occur for rare species that are truly present at the site (e.g., *E. obliquata*), and thus caution needs to be used to appropriately interpret eDNA data based on environmental conditions (e.g., flow) and detection strength (e.g., eDNA abundance and repeatability).

Current protocols for freshwater mussel visual tactile surveys describe methodological objectives, such as using timed searches to detect visible, larger mussels, while using transects and adaptive cluster methods to better capture buried or cryptic individuals (Smith 2006). In a similar sense, freshwater mussel eDNA surveys need to be designed based on objectives and goals, with projects focused on local mussel assemblages (site-level) and the detection of rare species requiring greater sampling rigor (i.e., number of samples and replicates) and careful planning around environmental conditions (e.g., flow) and biological behavior (e.g., reproductive activity). The effective source area being sampled during an eDNA survey is likely to fluctuate with these environmental and biological factors, yet this study found eDNA provided adequate mussel detection within the seasonal survey window currently prescribed for tactile searches.

## Acknowledgments

This project was supported through field support during eDNA surveys by Zoe True and Ashley Hansen from Stantec Consulting.

Data Availability Statement: Supplementary data files are provided to Open Science Framework at https://doi.org/XXXX. Raw sequence data are deposited in the National Center for Biotechnology Information (NCBI) Sequence Read Archive (SRA) under the BioProject accession XXXXX.

## Ethics and Permit Approval Statement

No permitting was required for eDNA surveying for this project.

## Conflict of Interest Disclosure

The authors declare no conflicts of interest.

Permission to Reproduce Material from other sources: All material is original to this manuscript. Data from the visual survey was acquired from USFWS.

---

**Figure 1.**
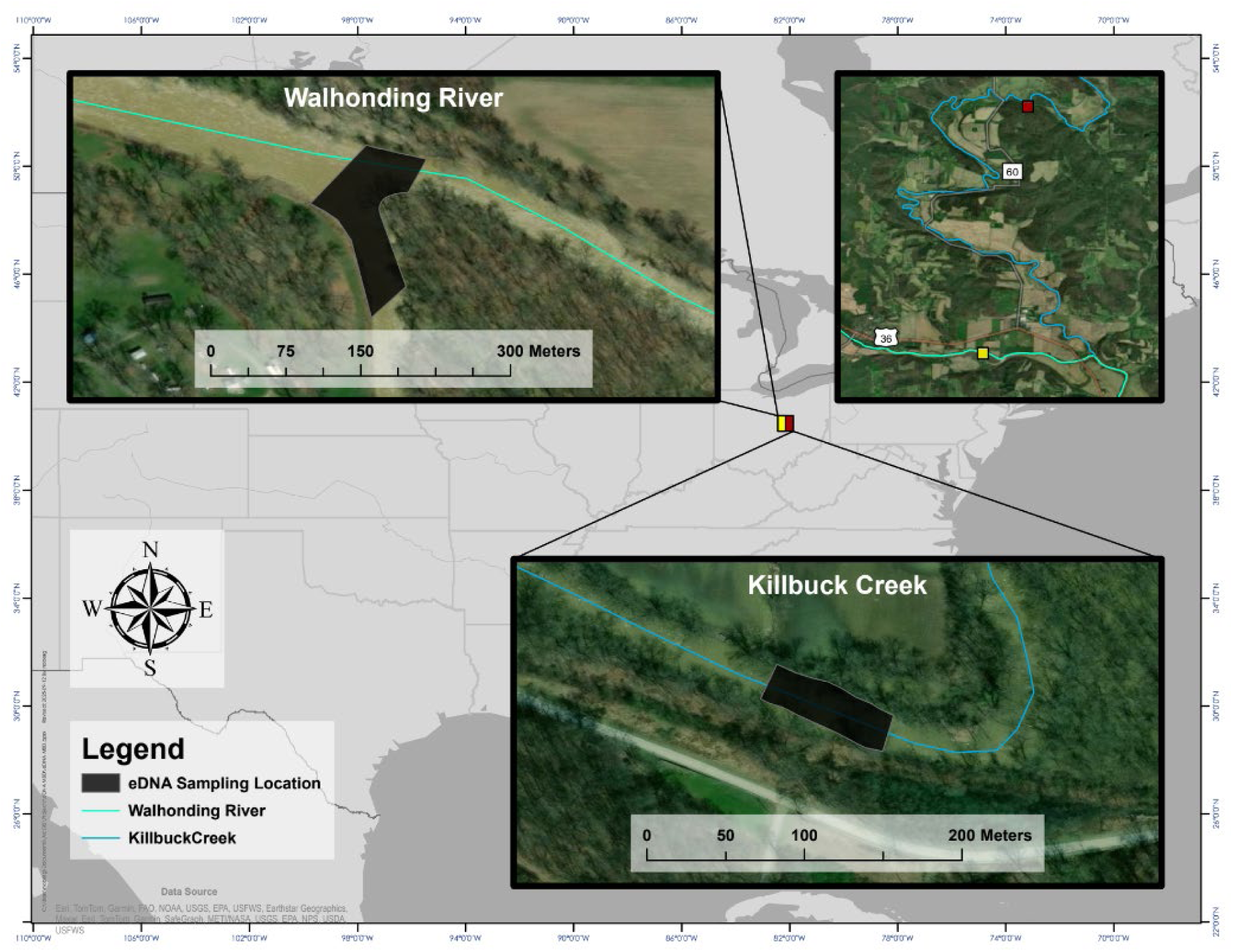
Map of eDNA survey sites sampled in the Walhonding River and Killbuck Creek.

**Supplementary Figure 1.**
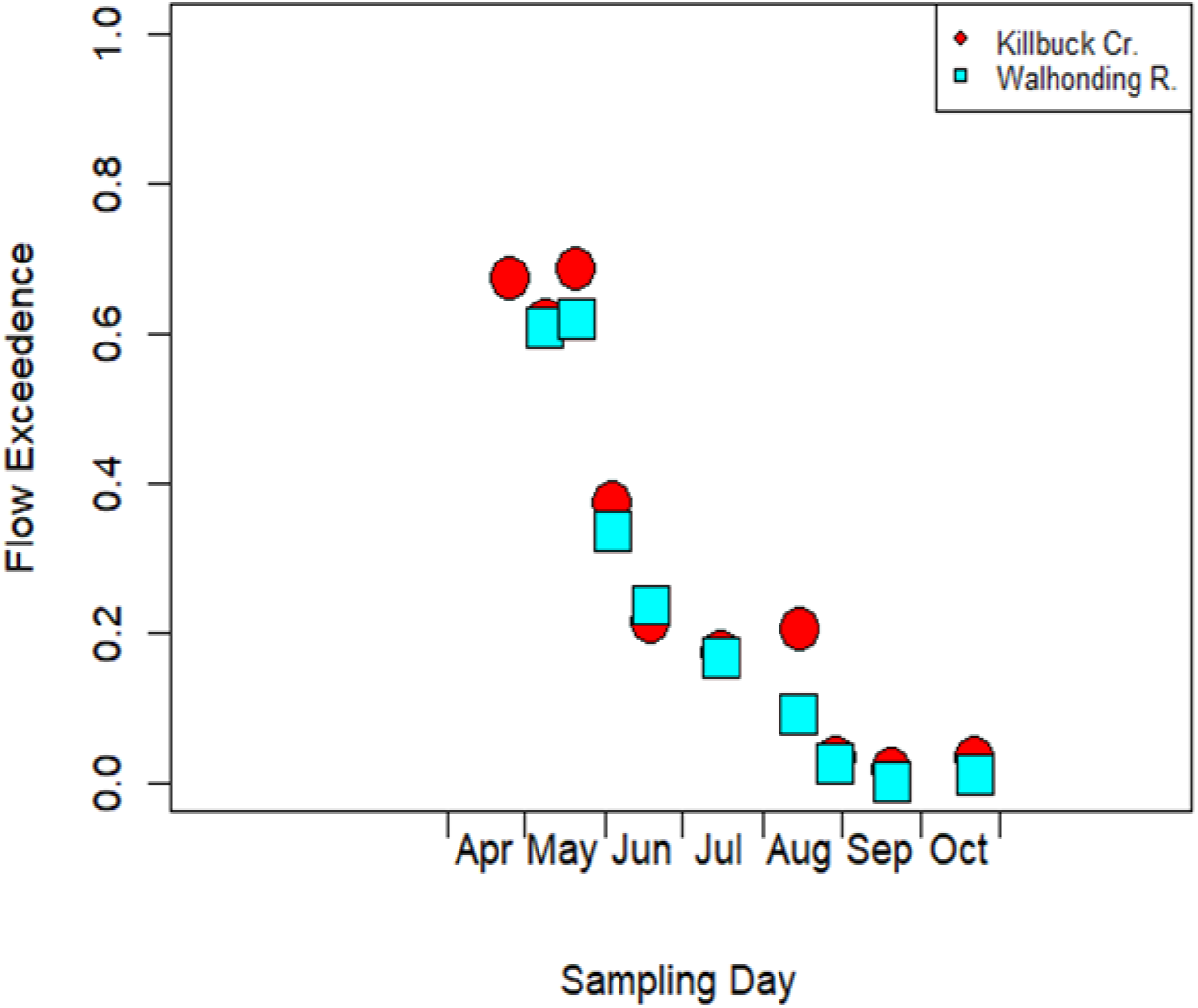
Flow exceedance trends across sampling events in Killbuck Creek and Walhonding River.

**Supplementary Figure 2.**
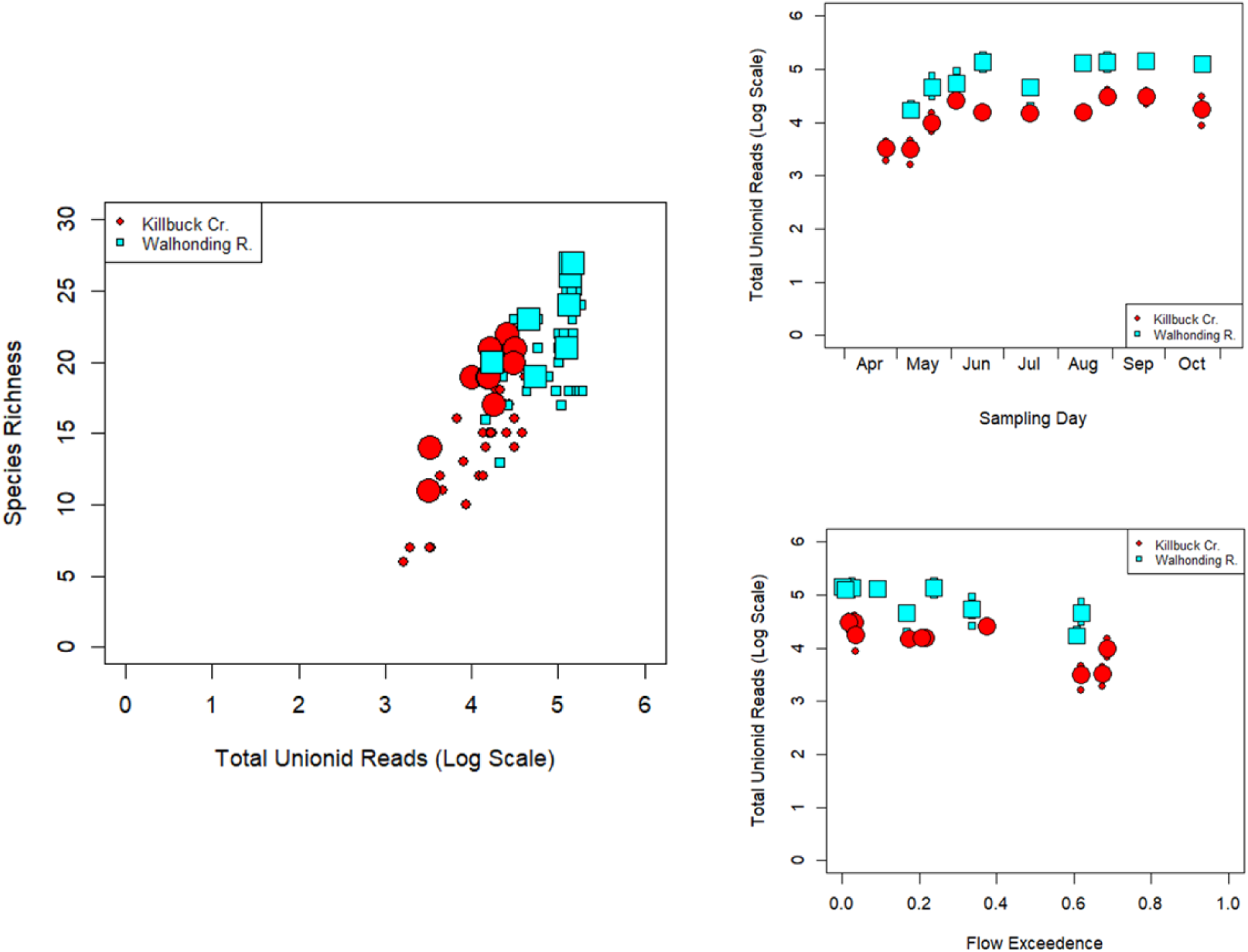
Total mussel sequence read abundance across sampling events and flow exceedances in Killbuck Creek and Walhonding River.

**Supplementary Figure 3.**
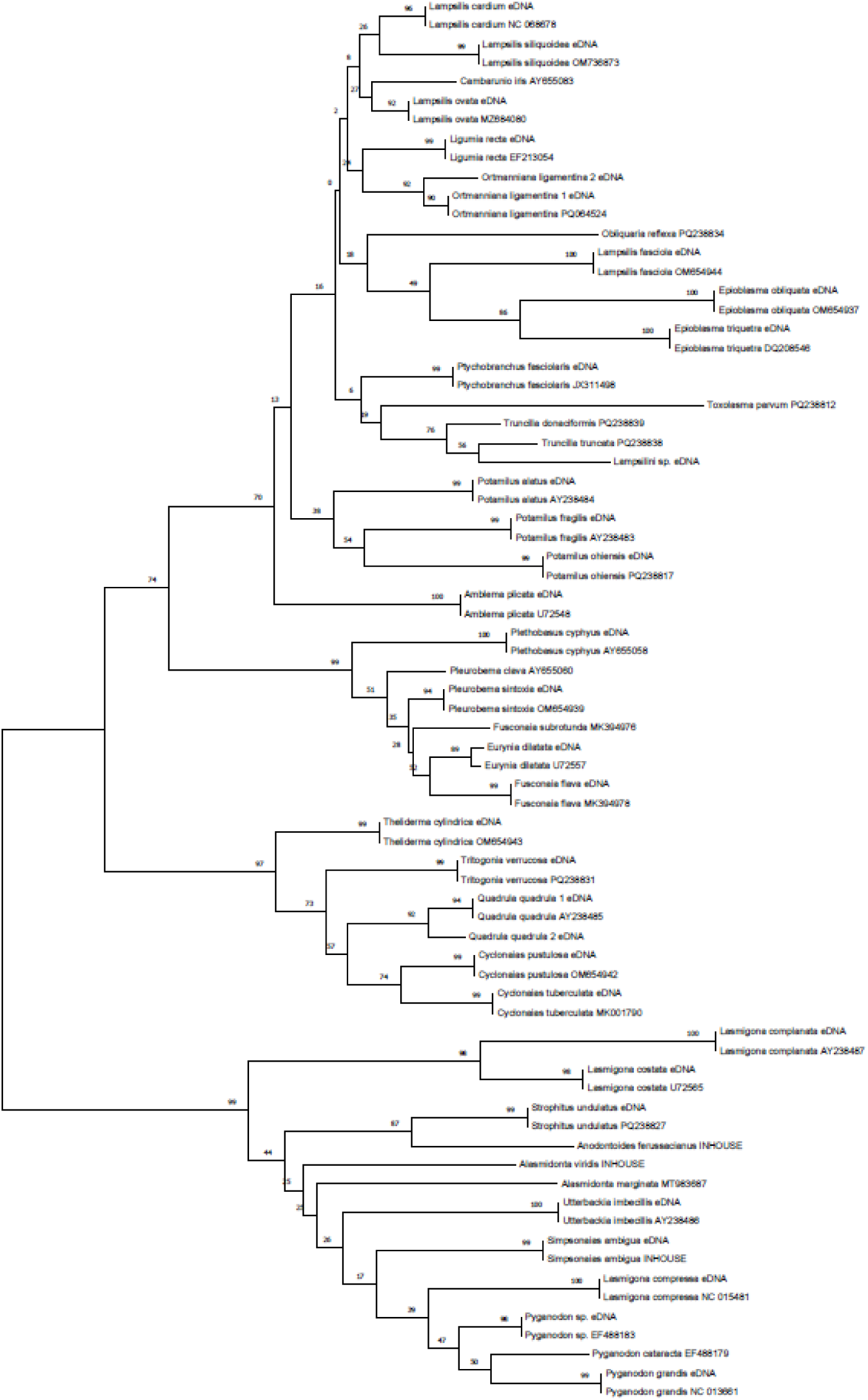
Phylogenetic tree used for taxonomic identification of the female mussel environmental DNA sequences based on the genetic match to known reference genetic data within the NCBI Genbank repository and in-house tissue sequencing.

**Supplementary Figure 4.**
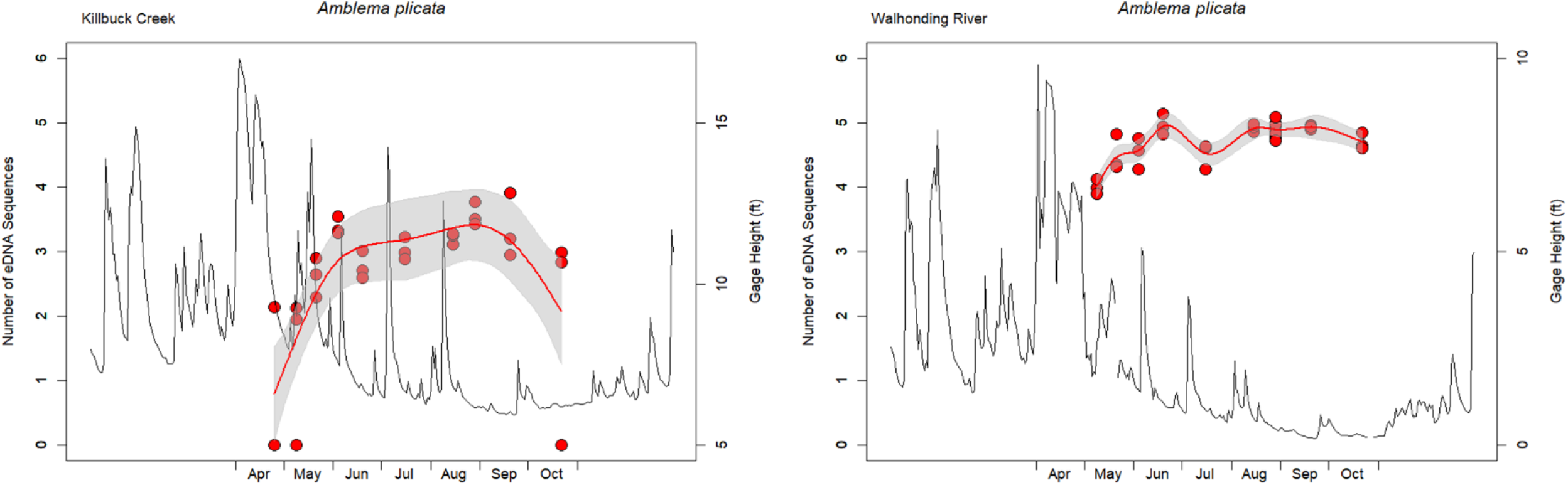
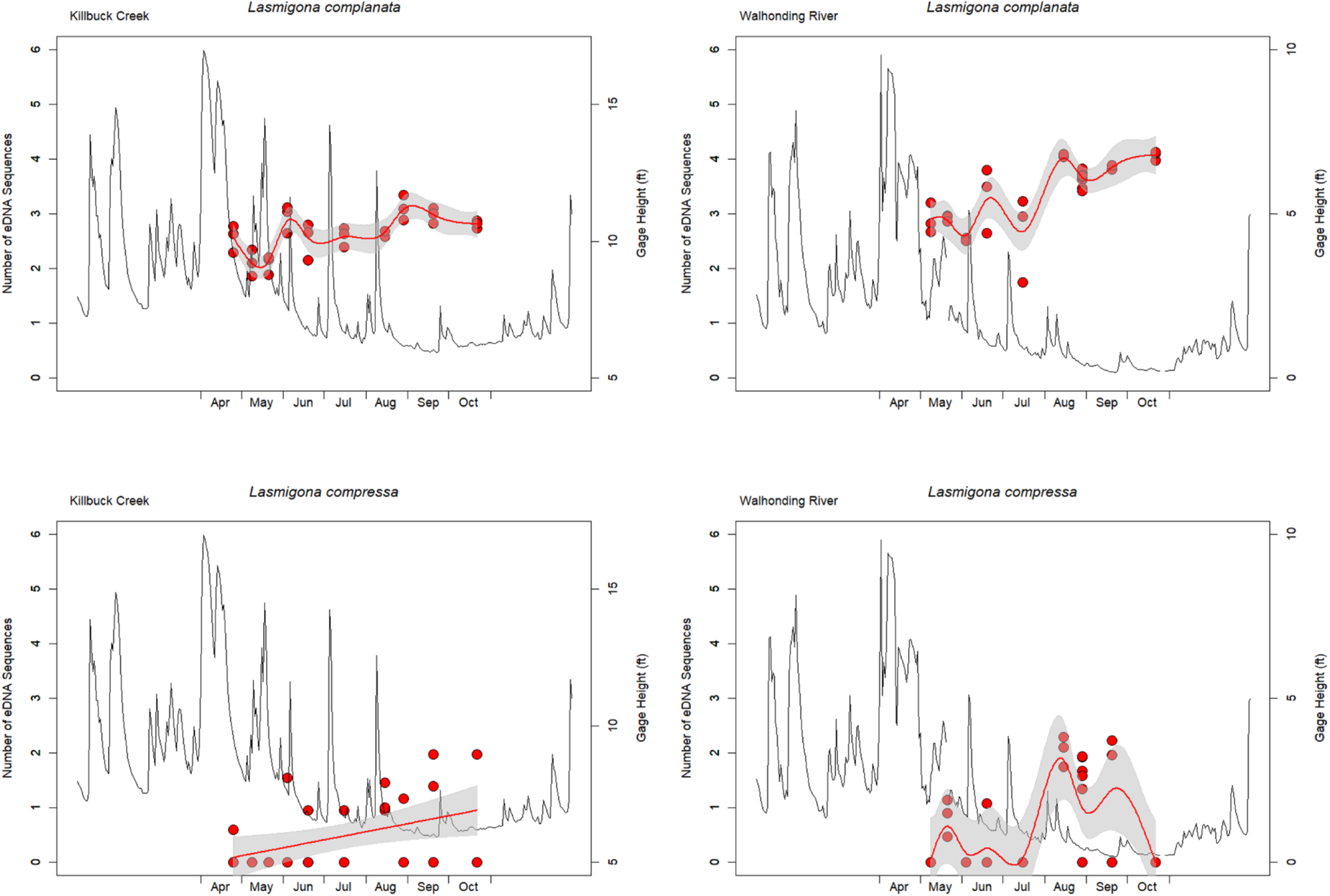
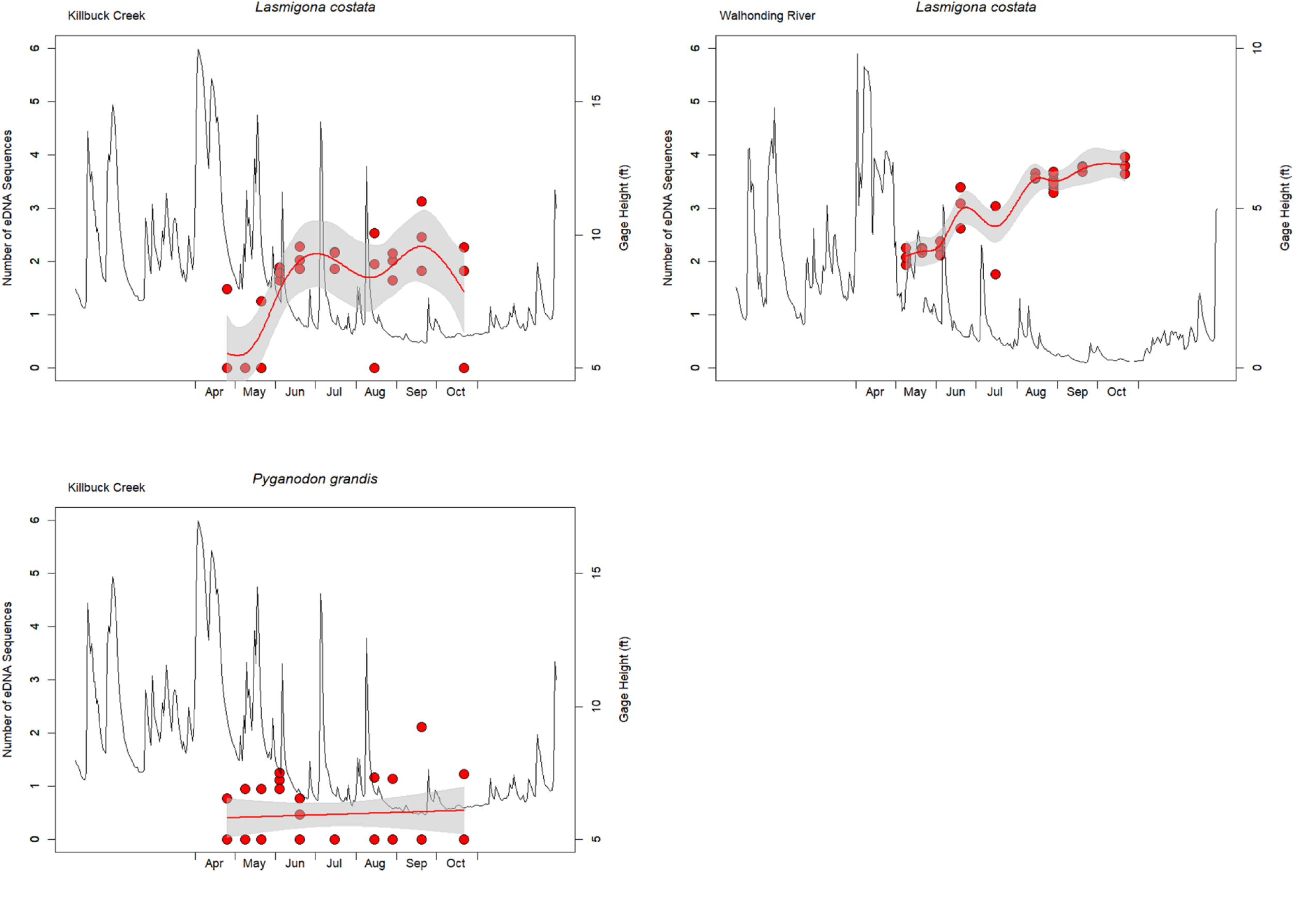
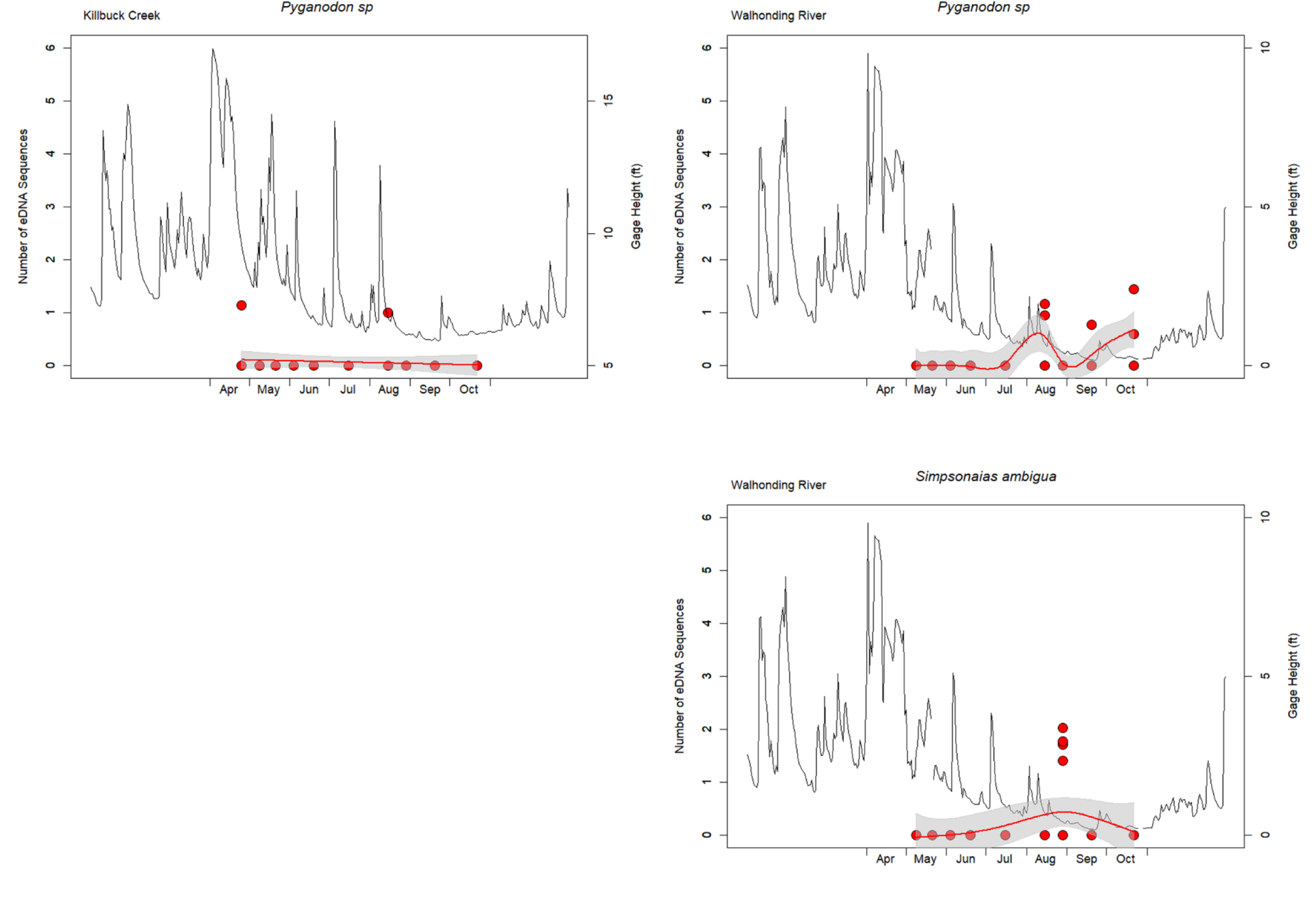
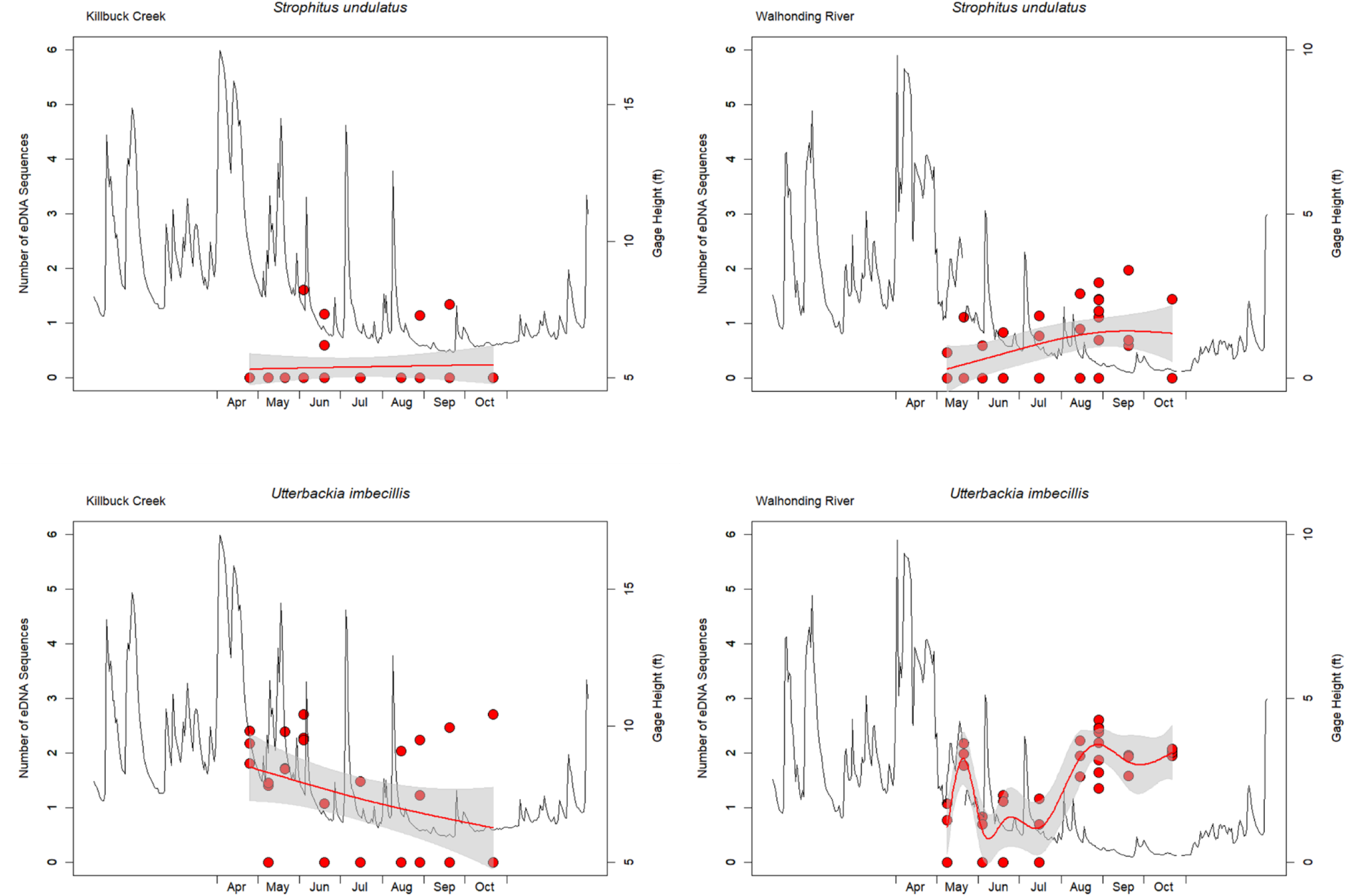
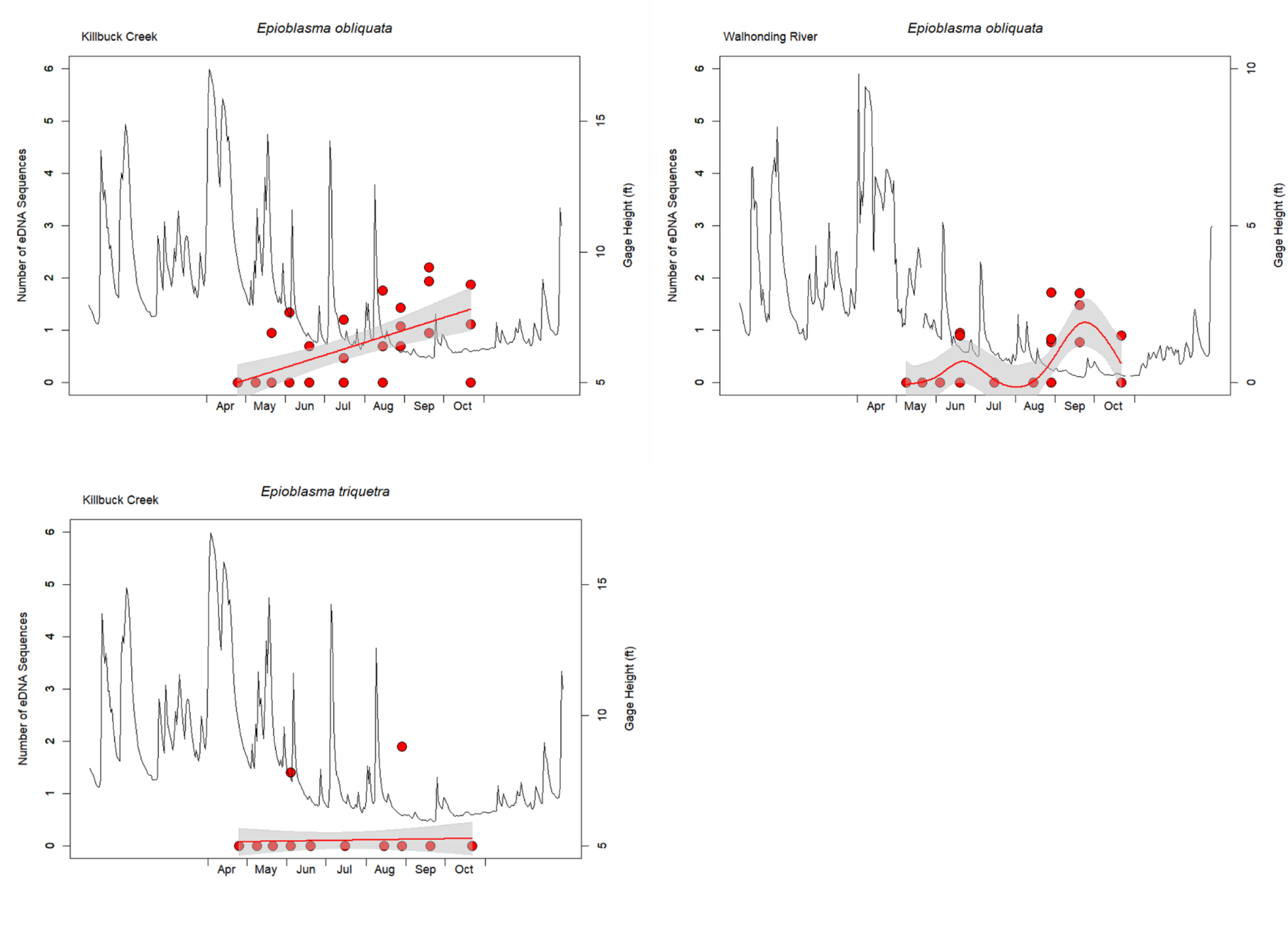
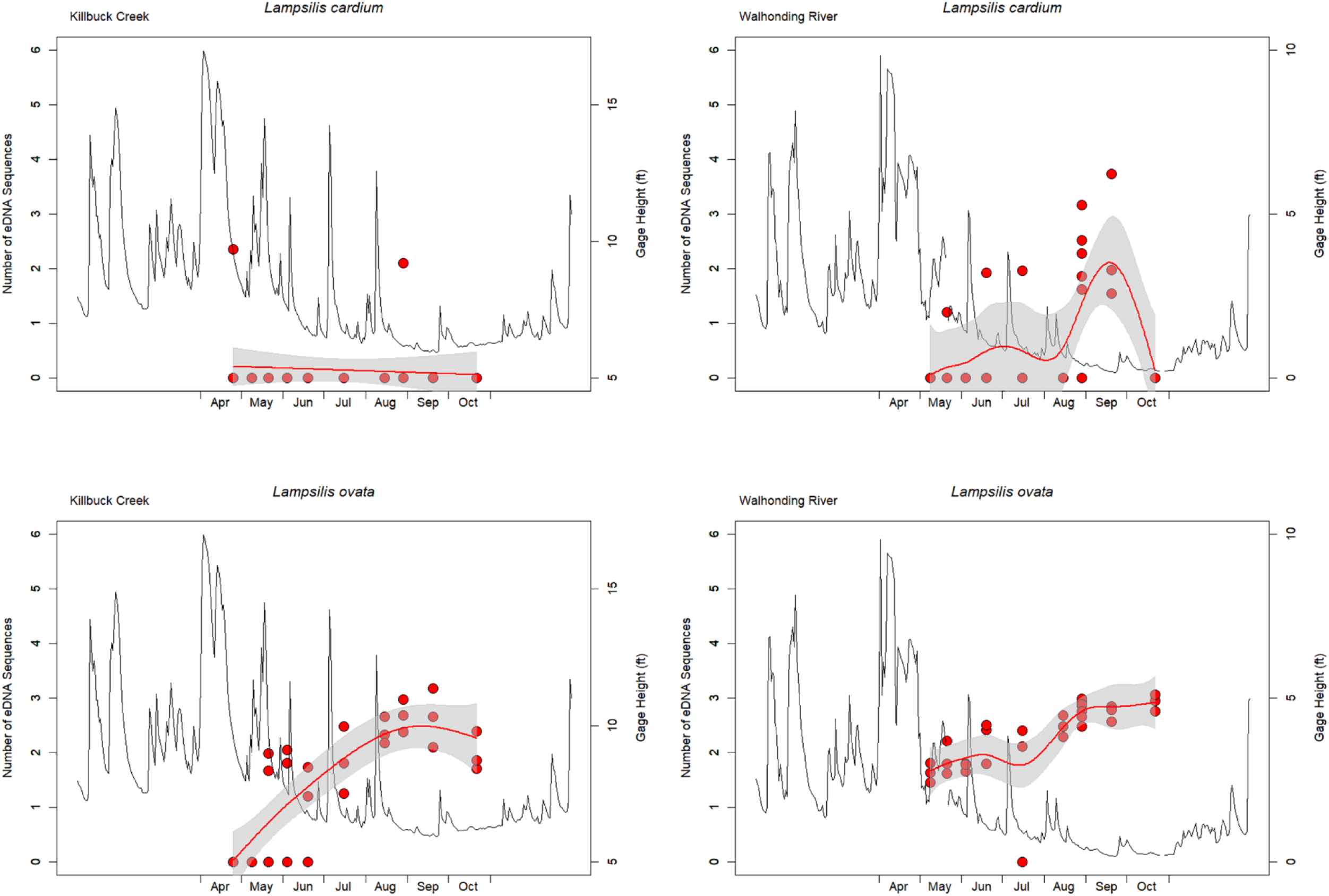
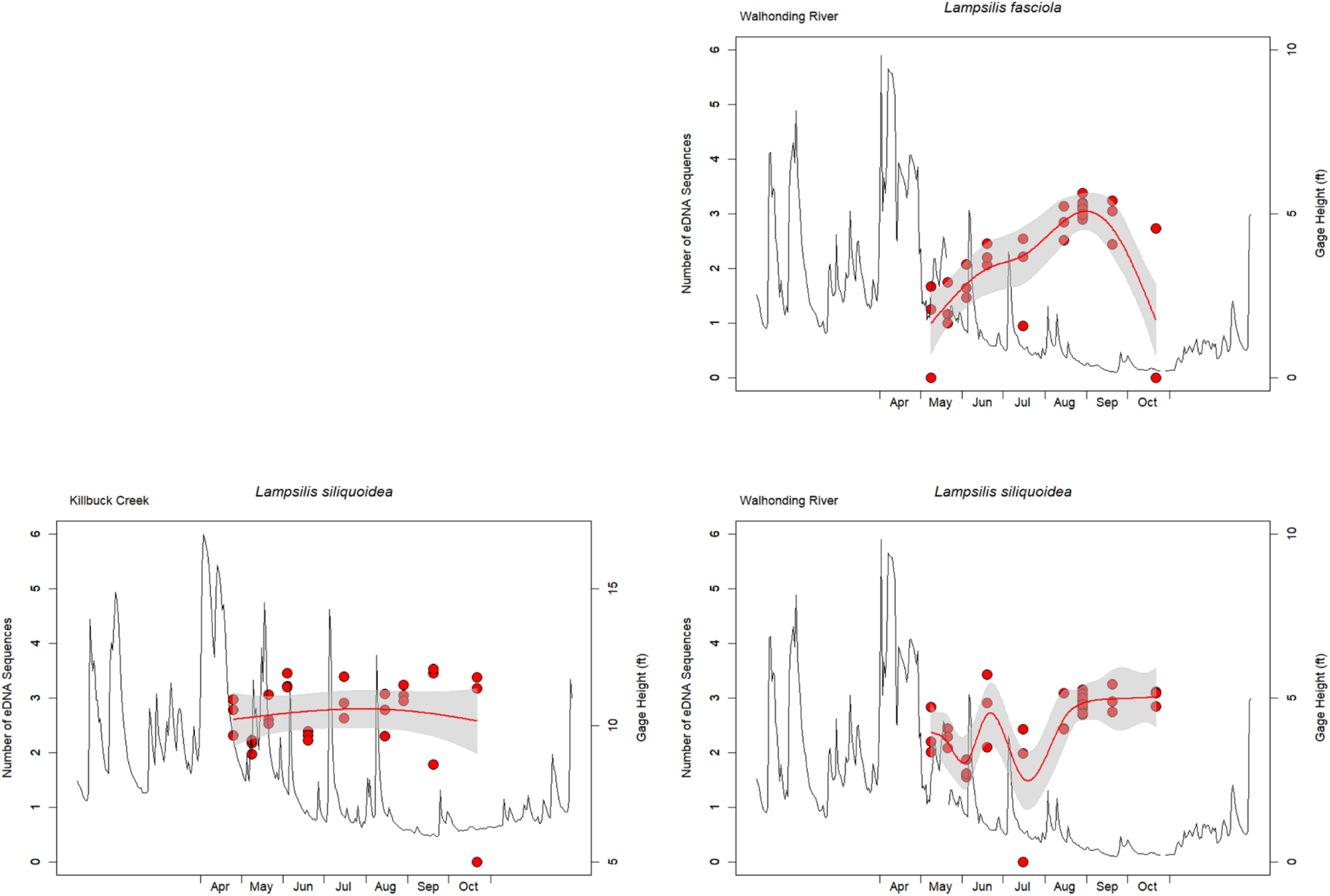
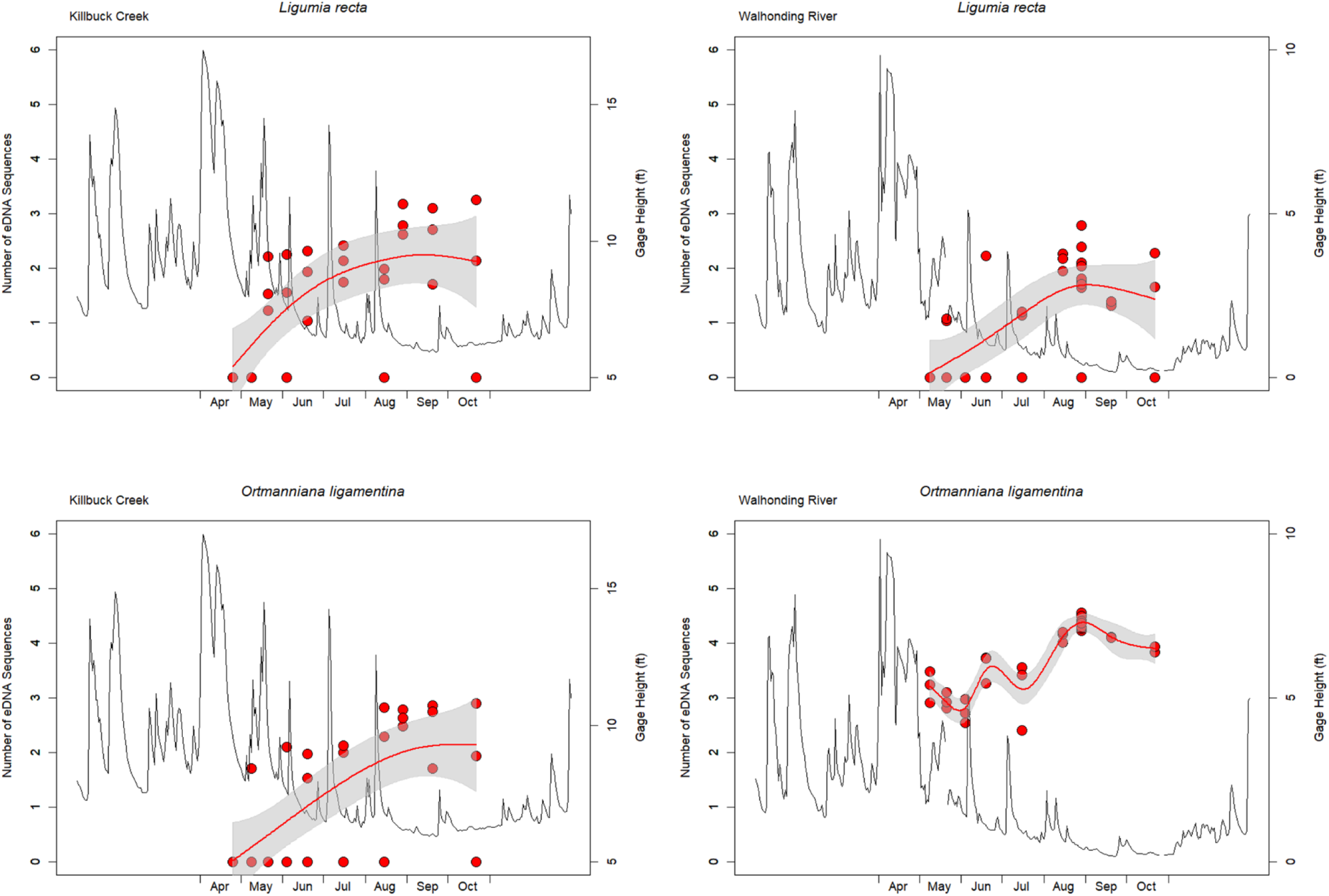
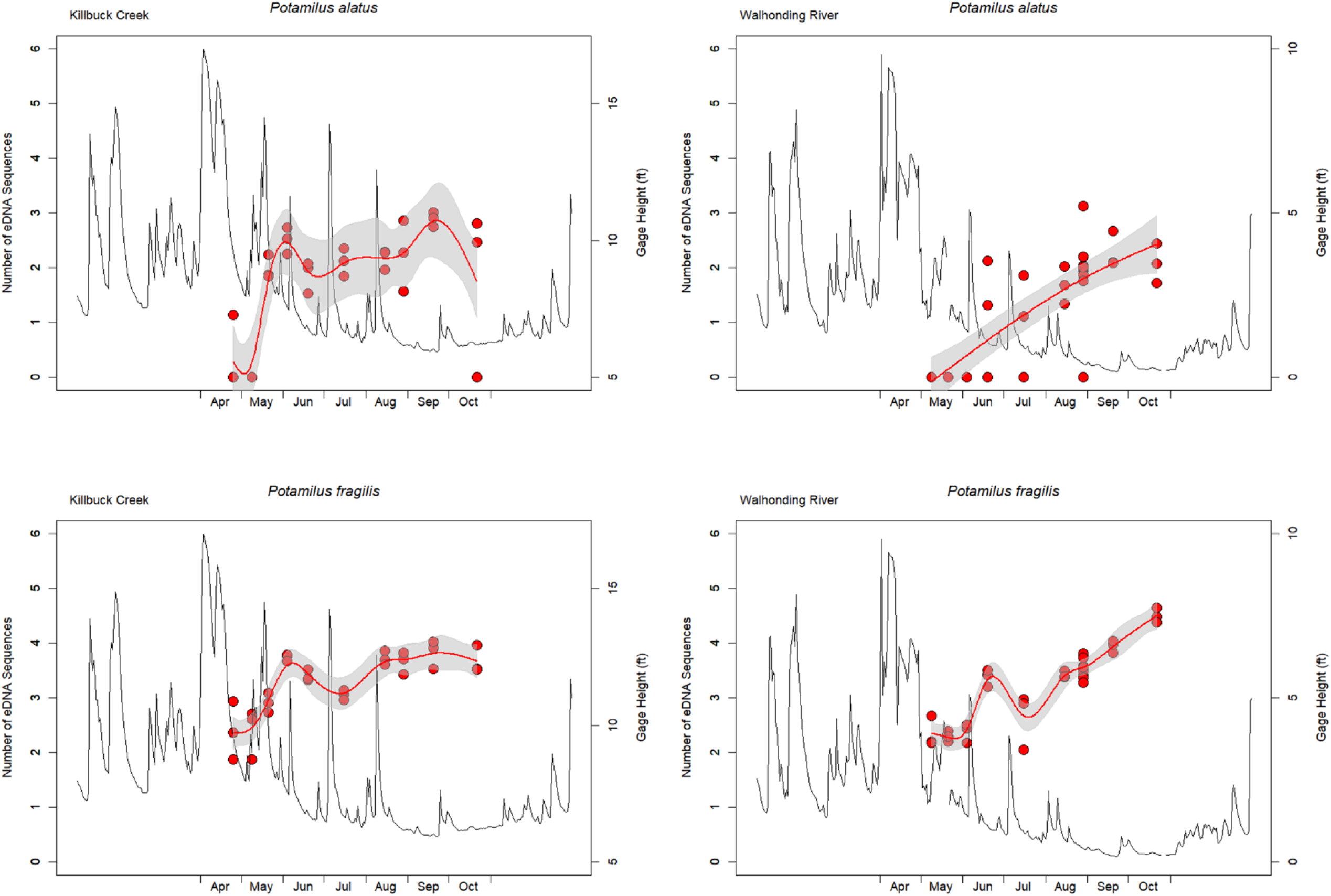
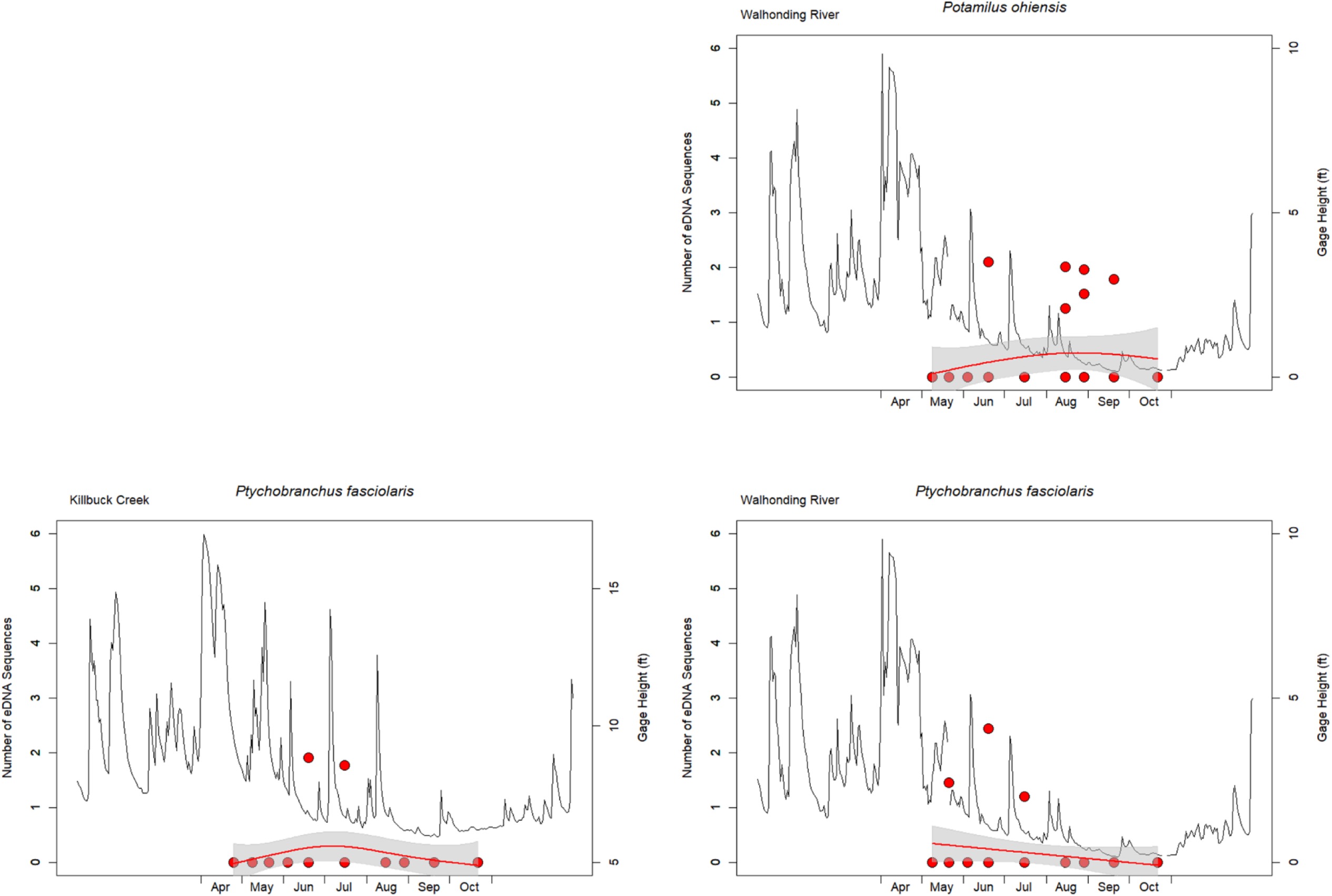
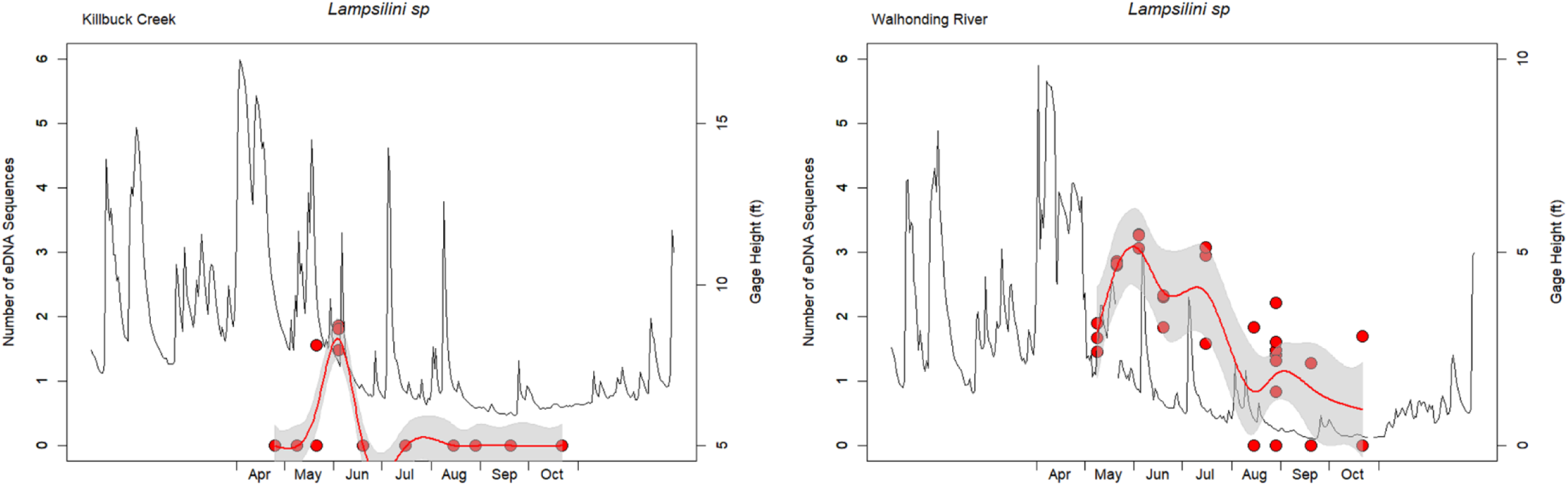
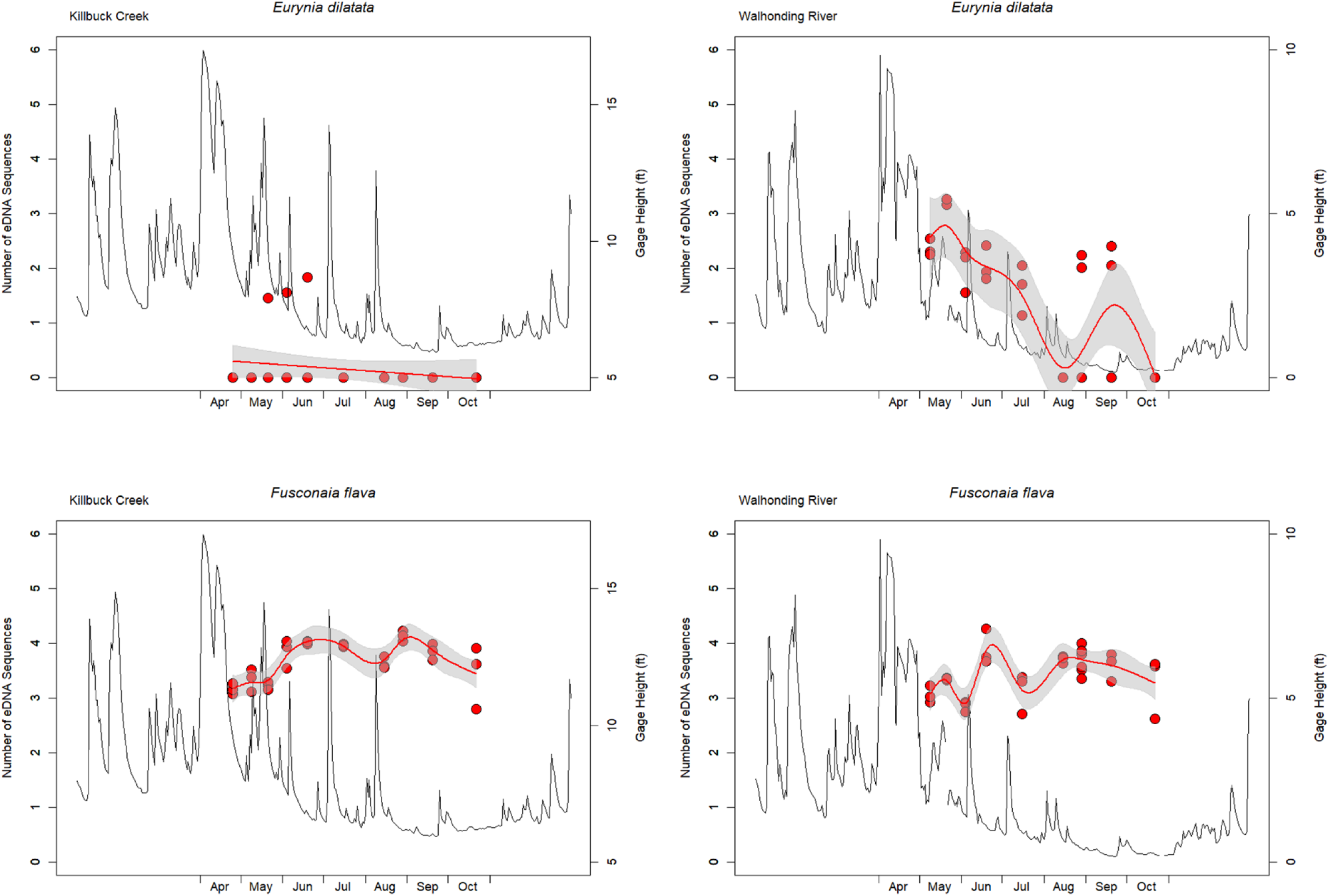
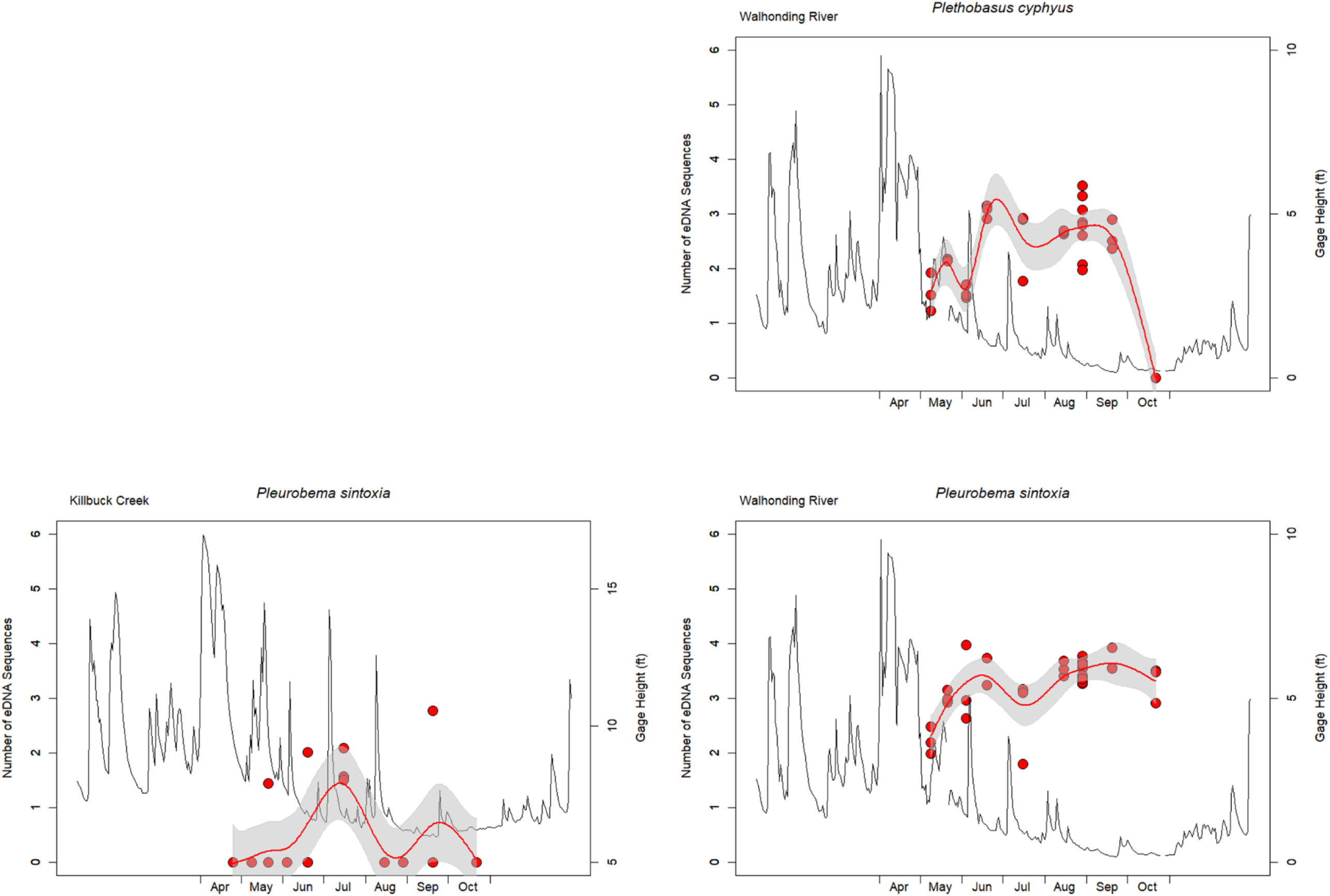
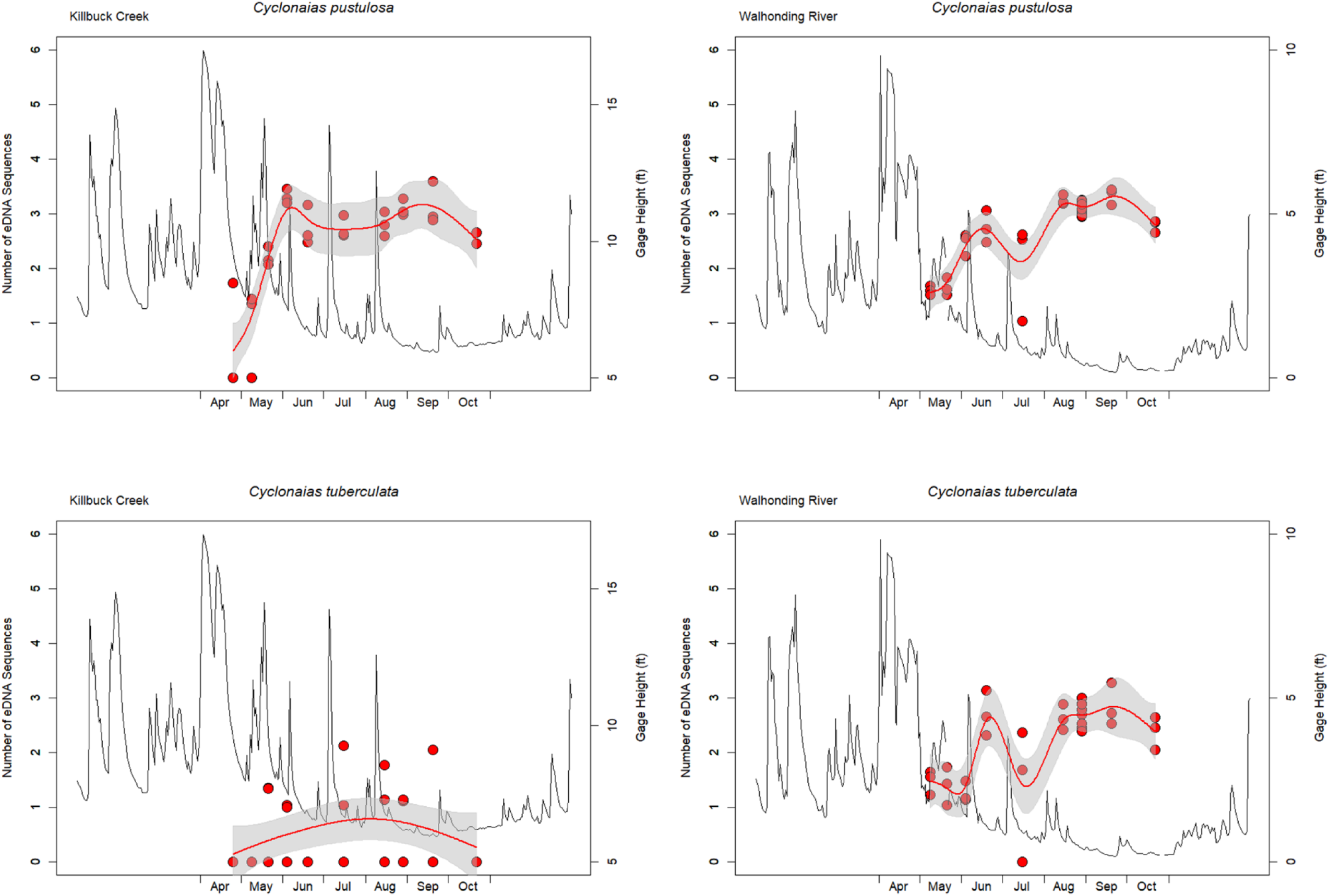
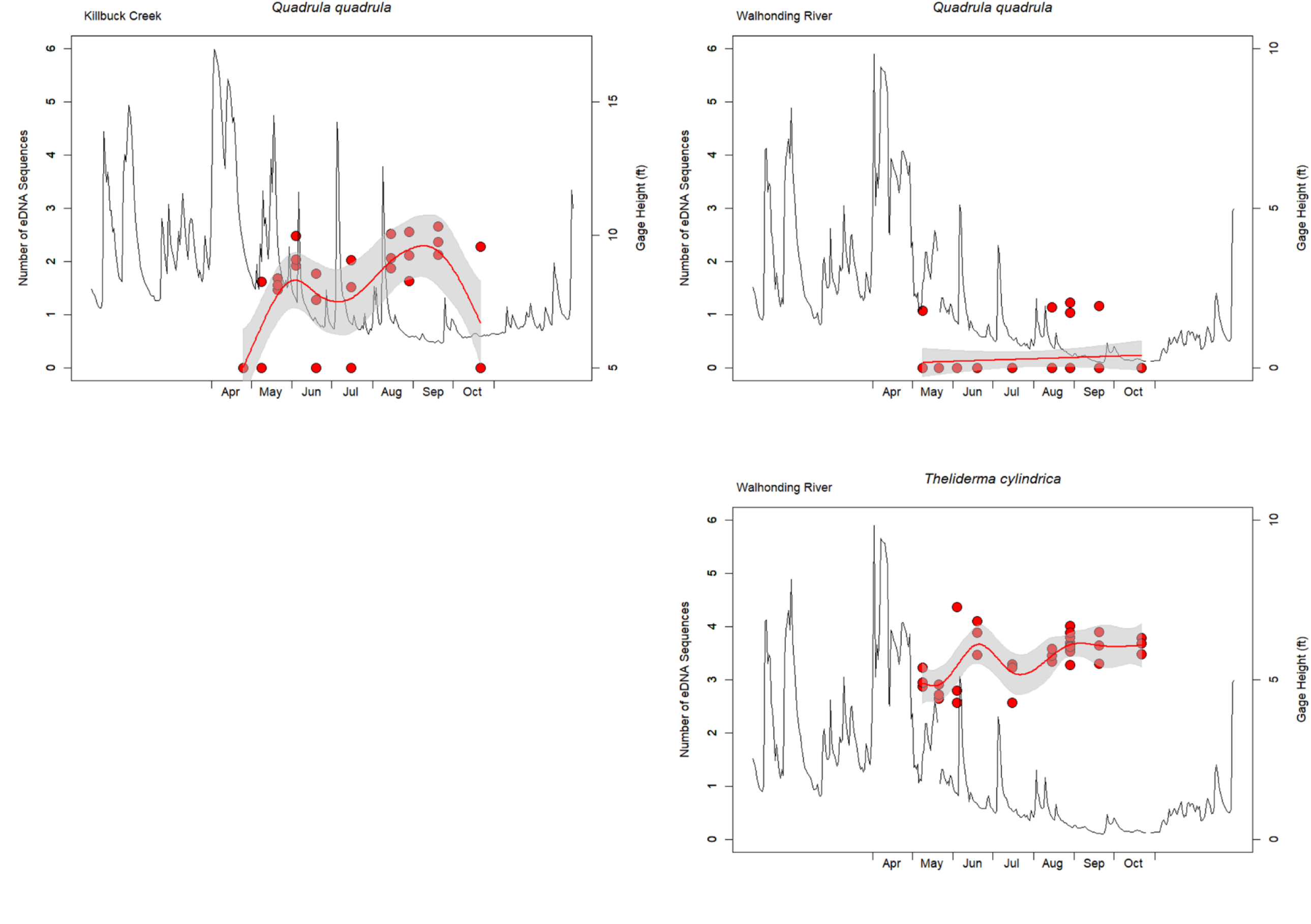
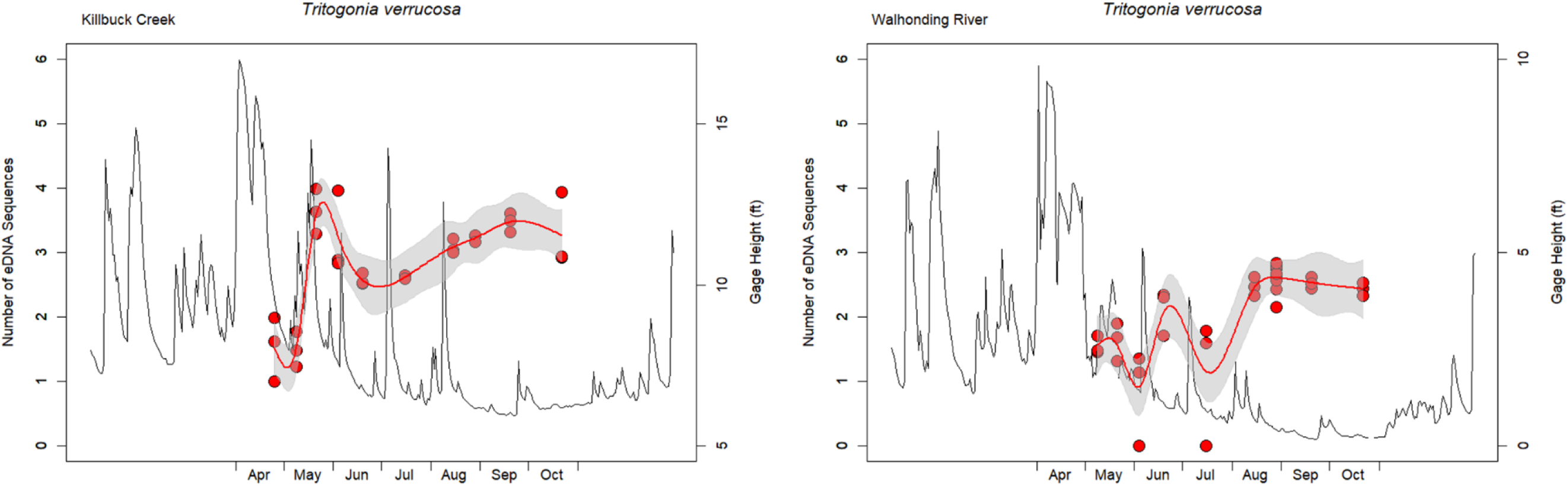
The eDNA abundance (i.e., the number of sequence reads) across all replicates on each sampling date from Killbuck Creek and Walhonding River for each species. Plots are organized by species and grouped within the major mussel tribes. The thin black line displays the USGS gage height (ft) across the sampling season.

**Supplementary Figure 5.**
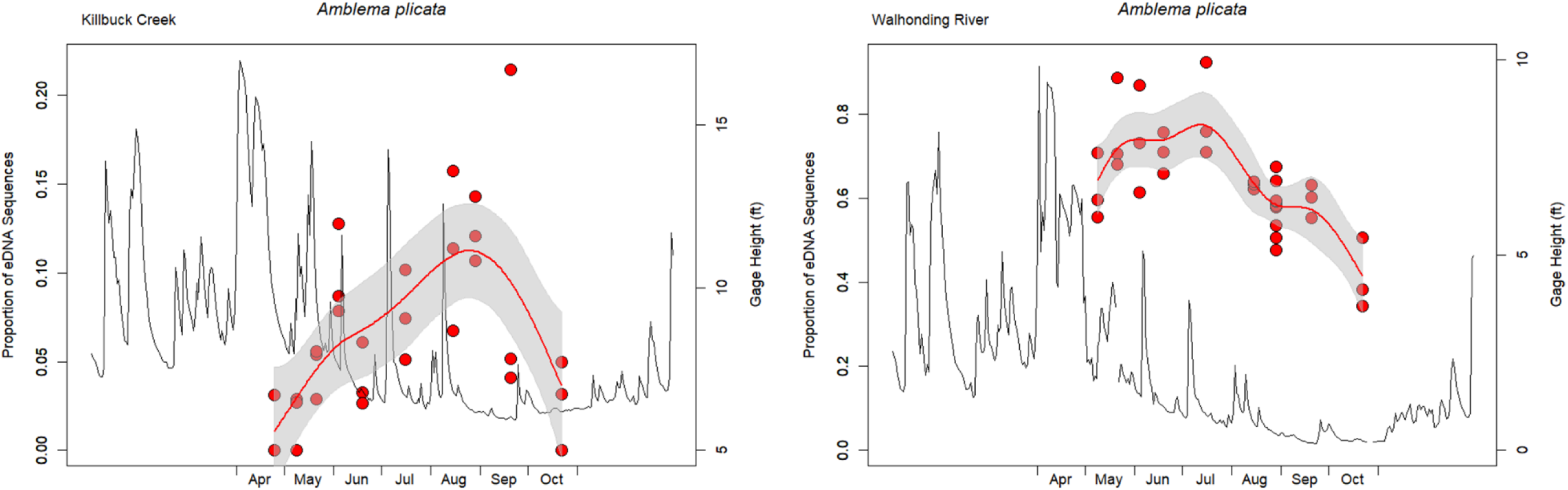
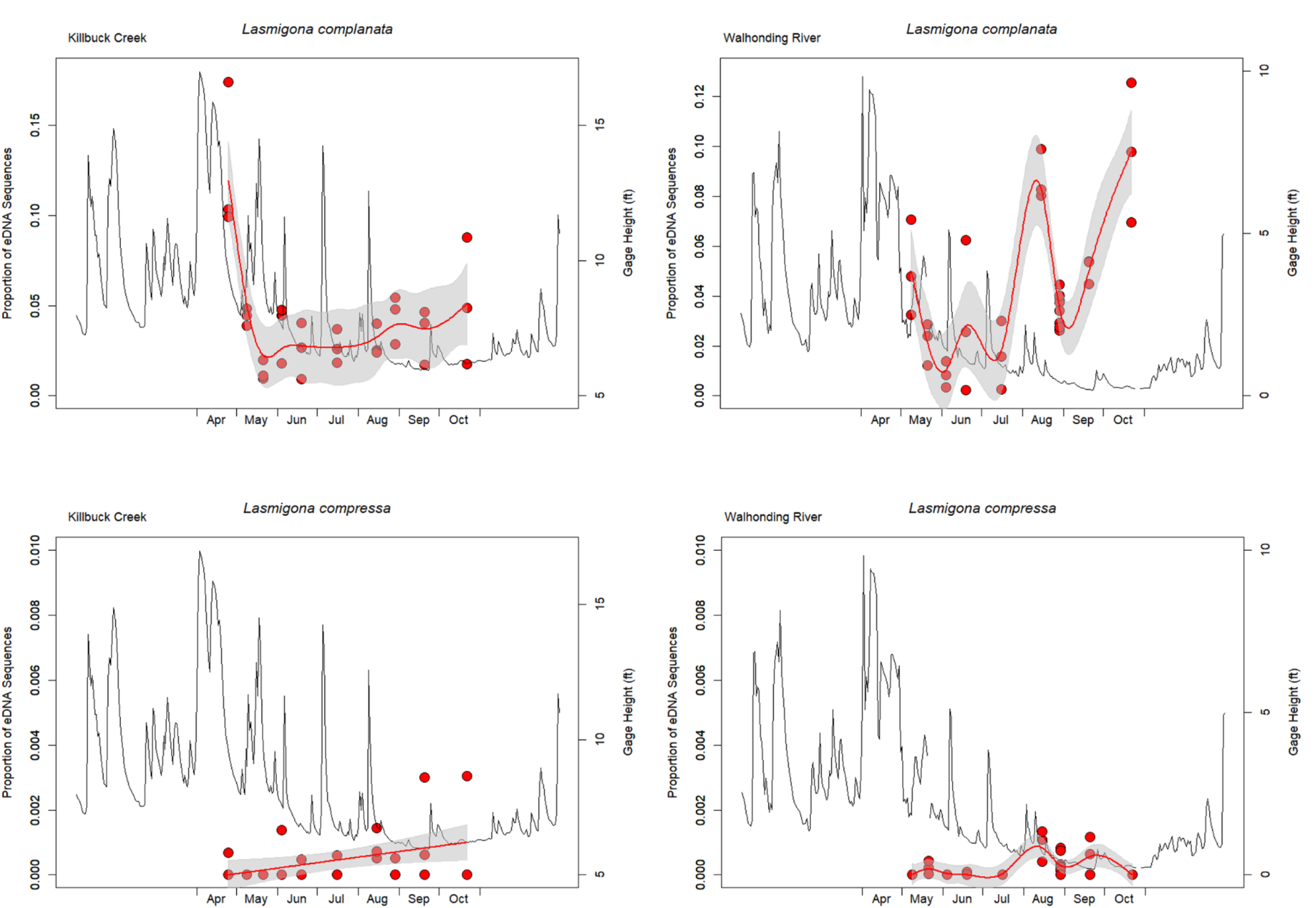
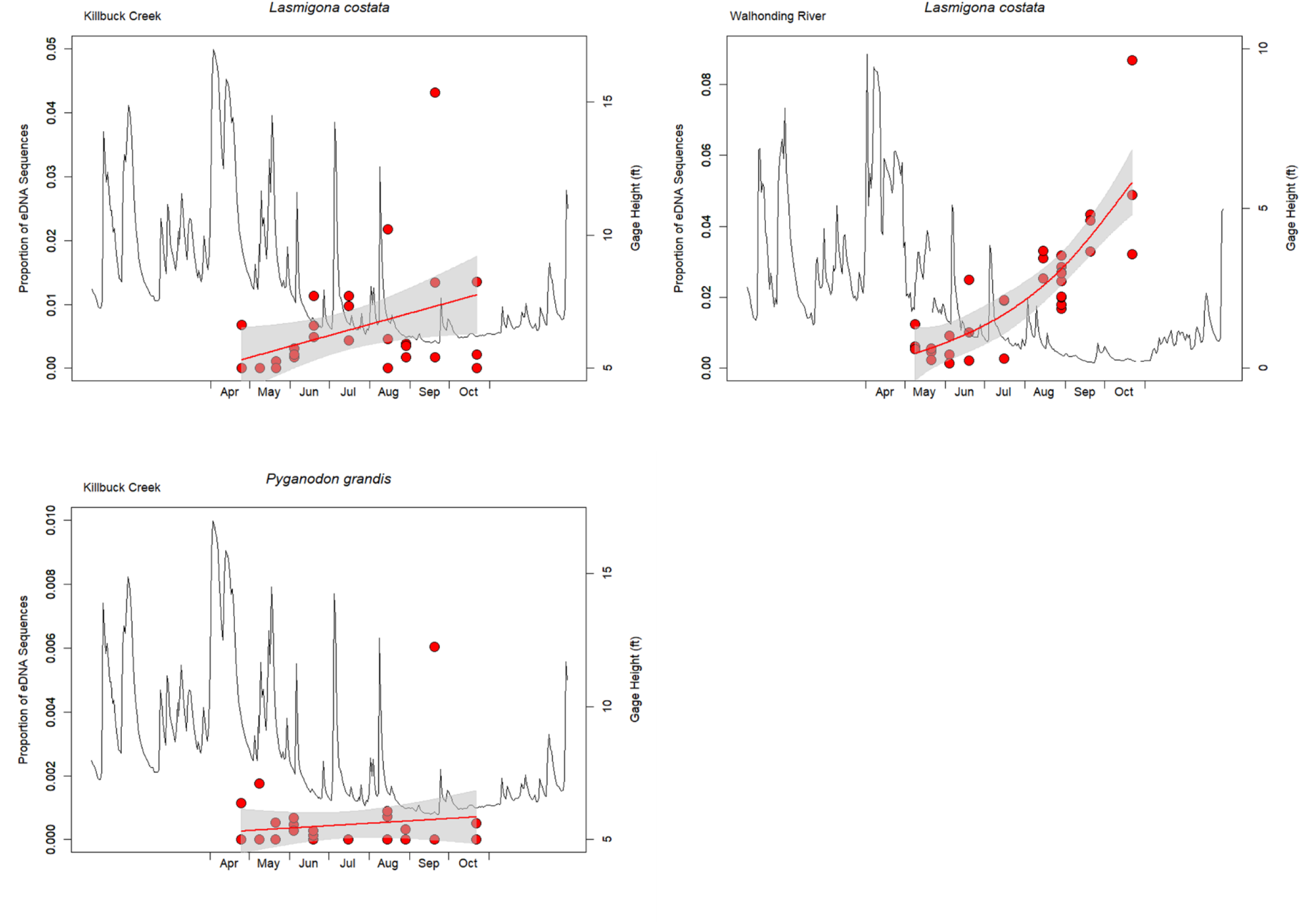
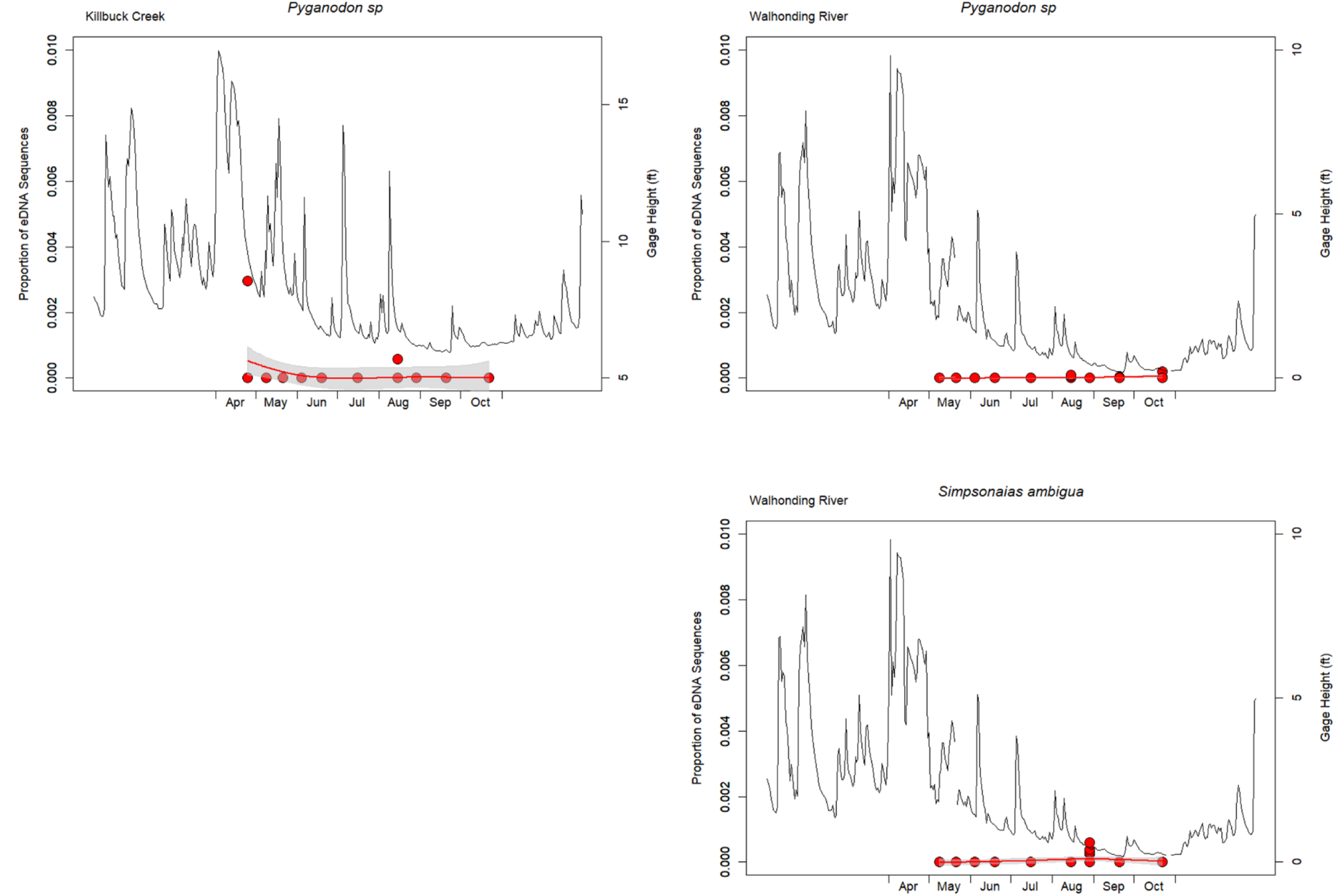
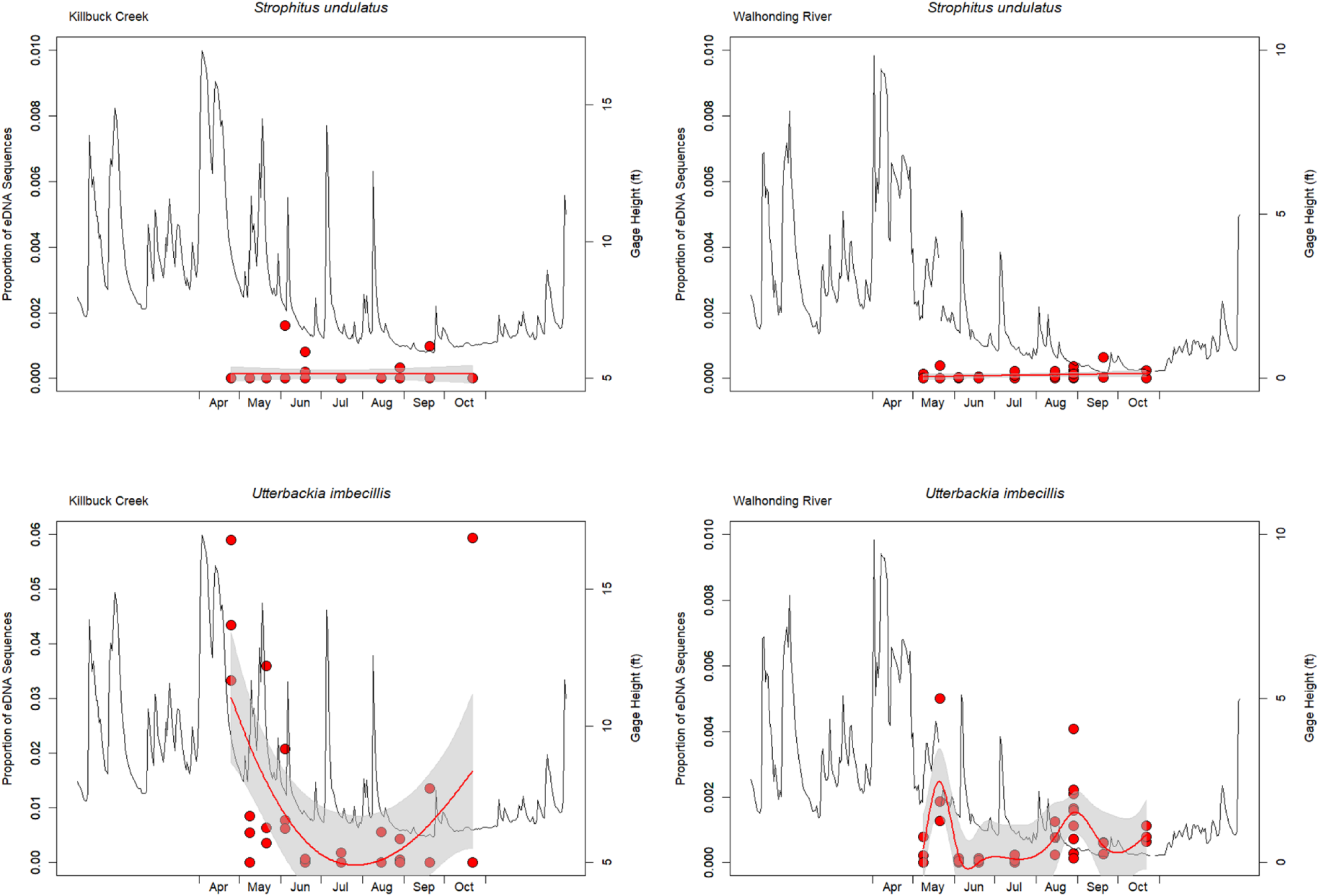
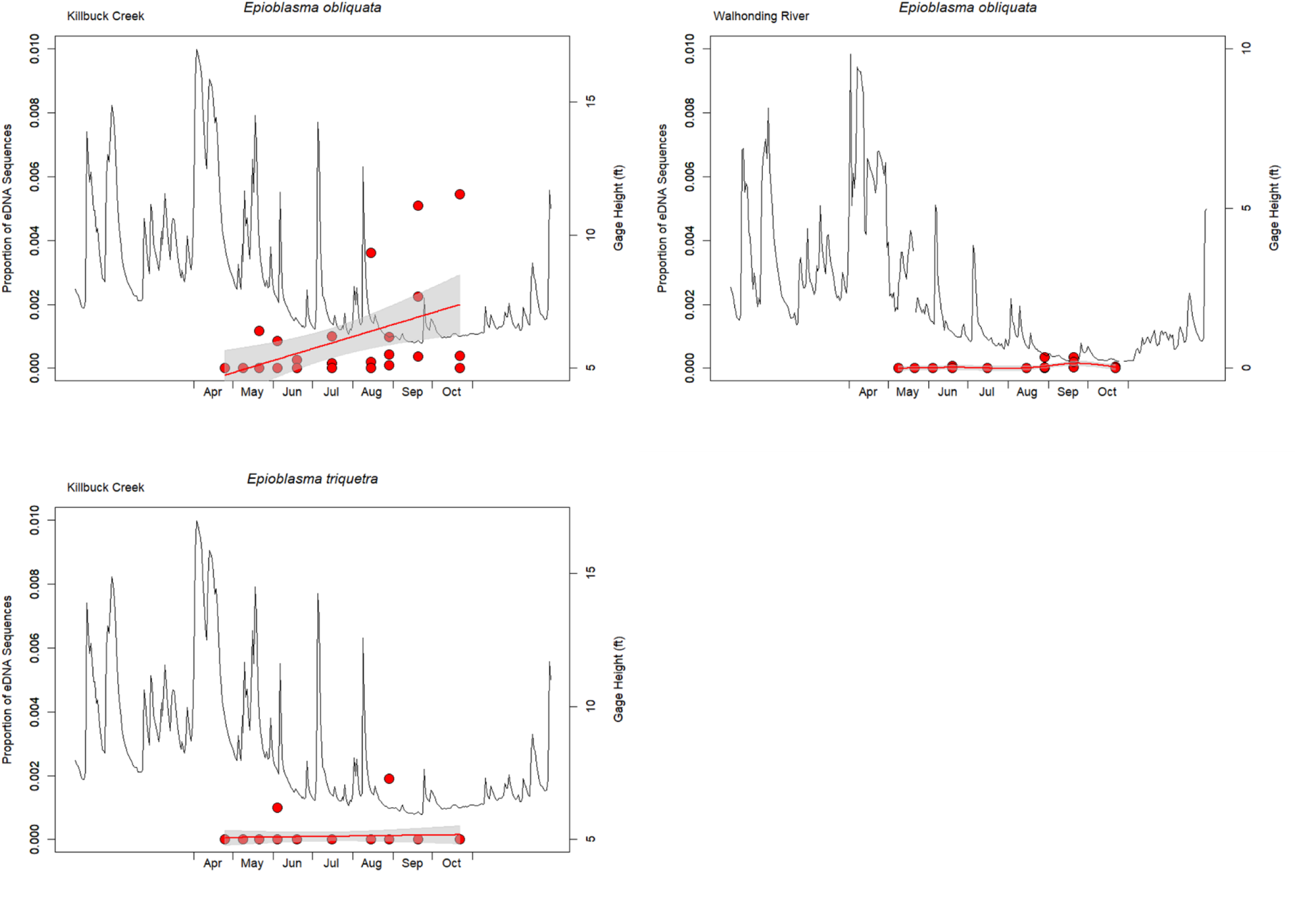
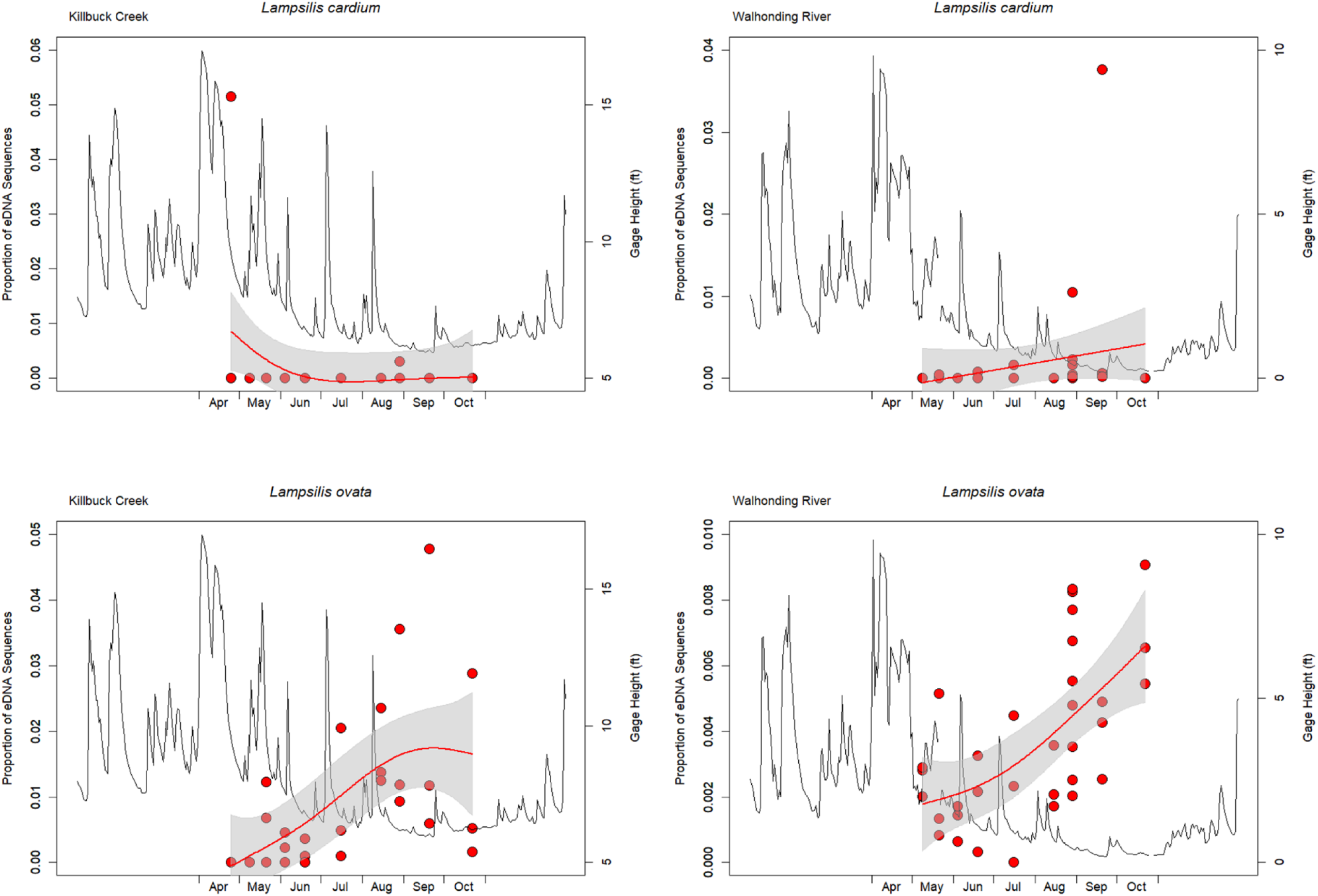
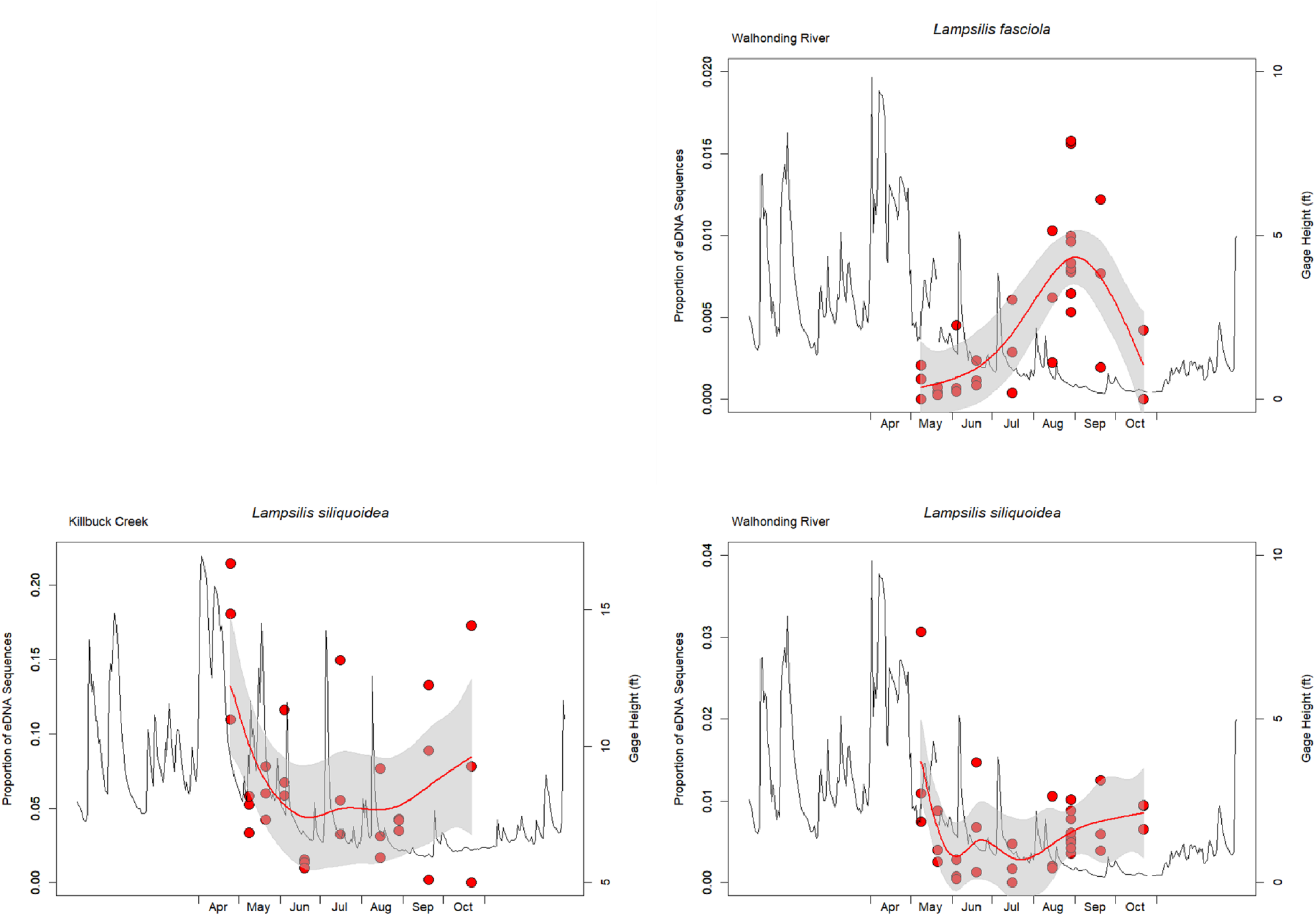
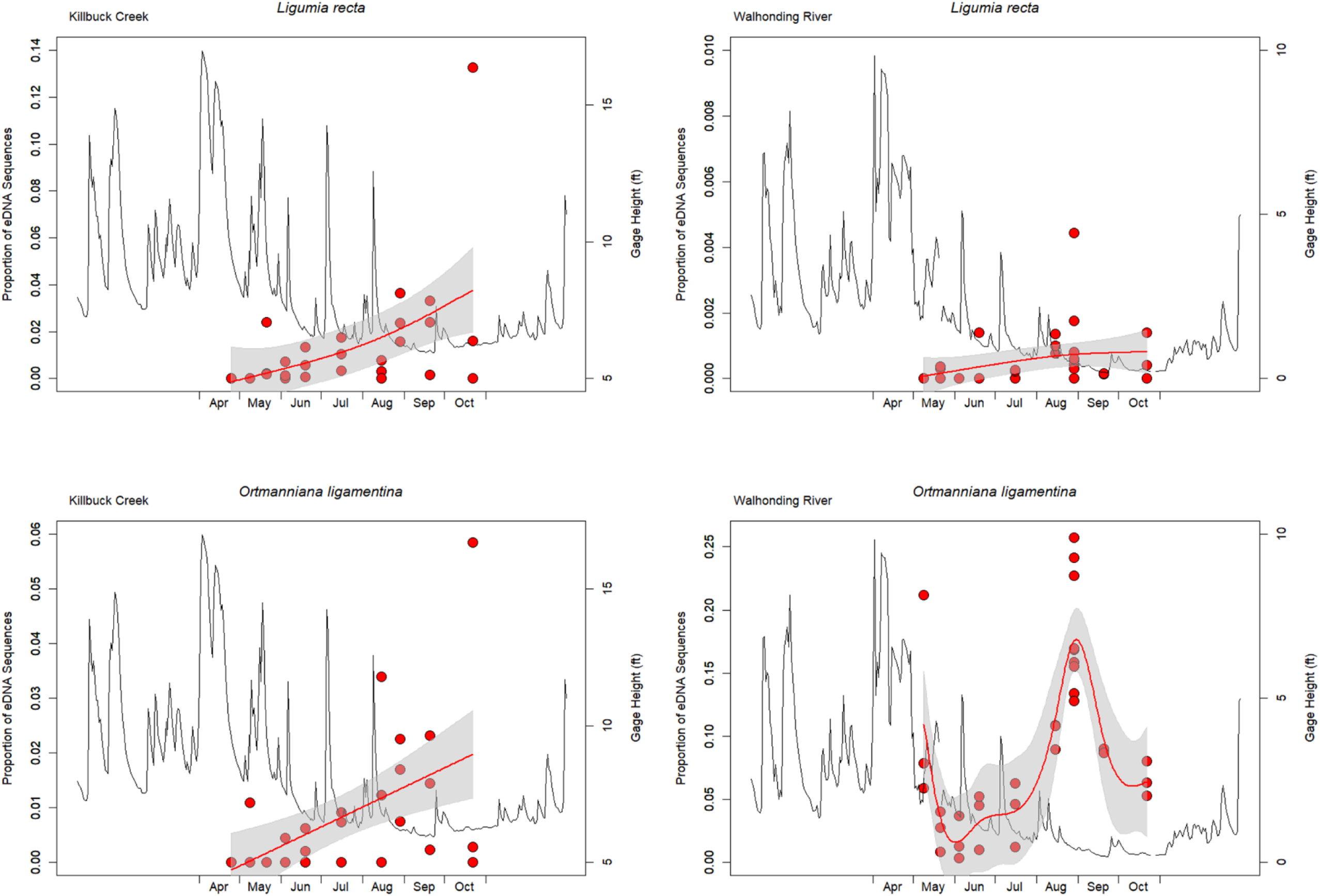
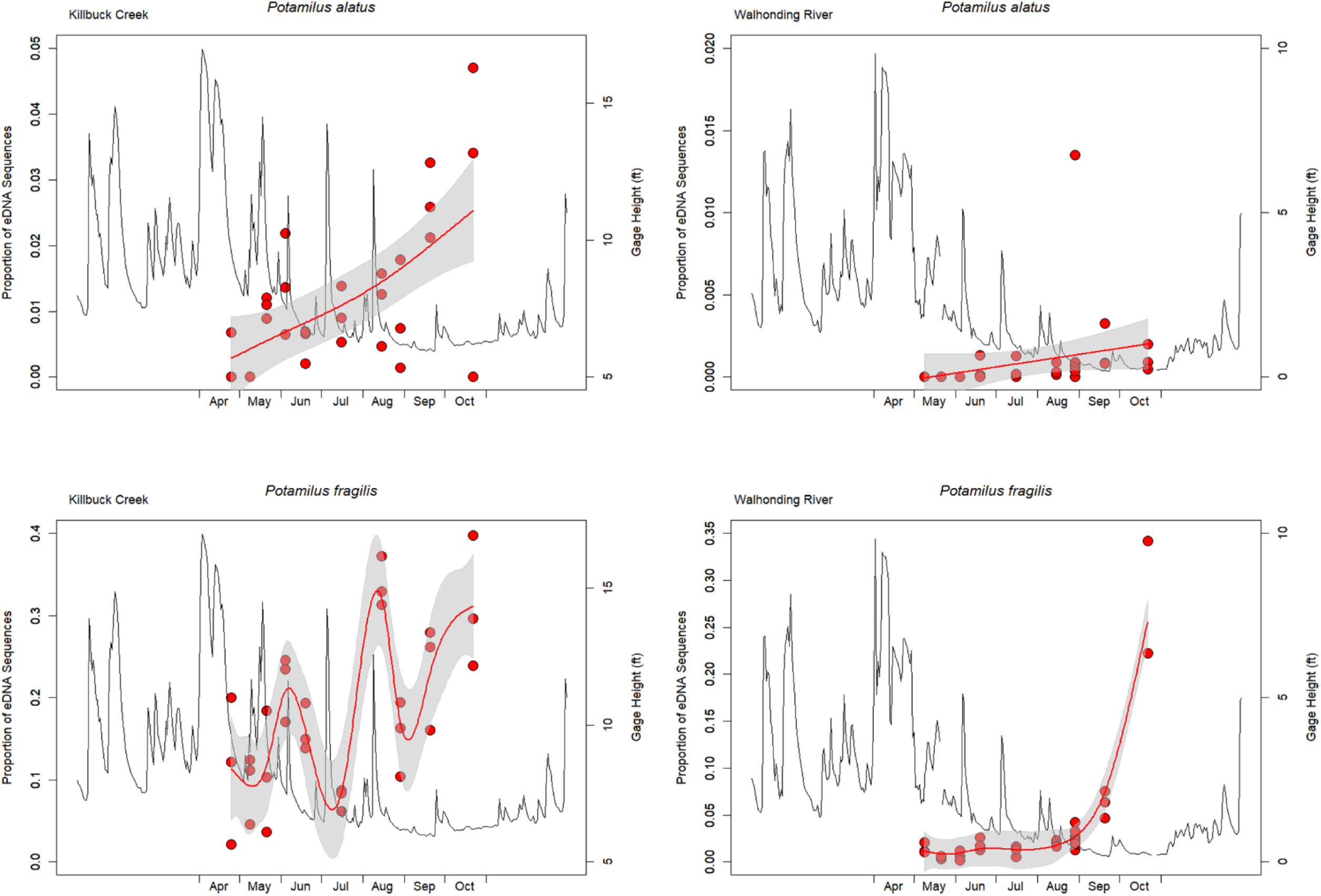
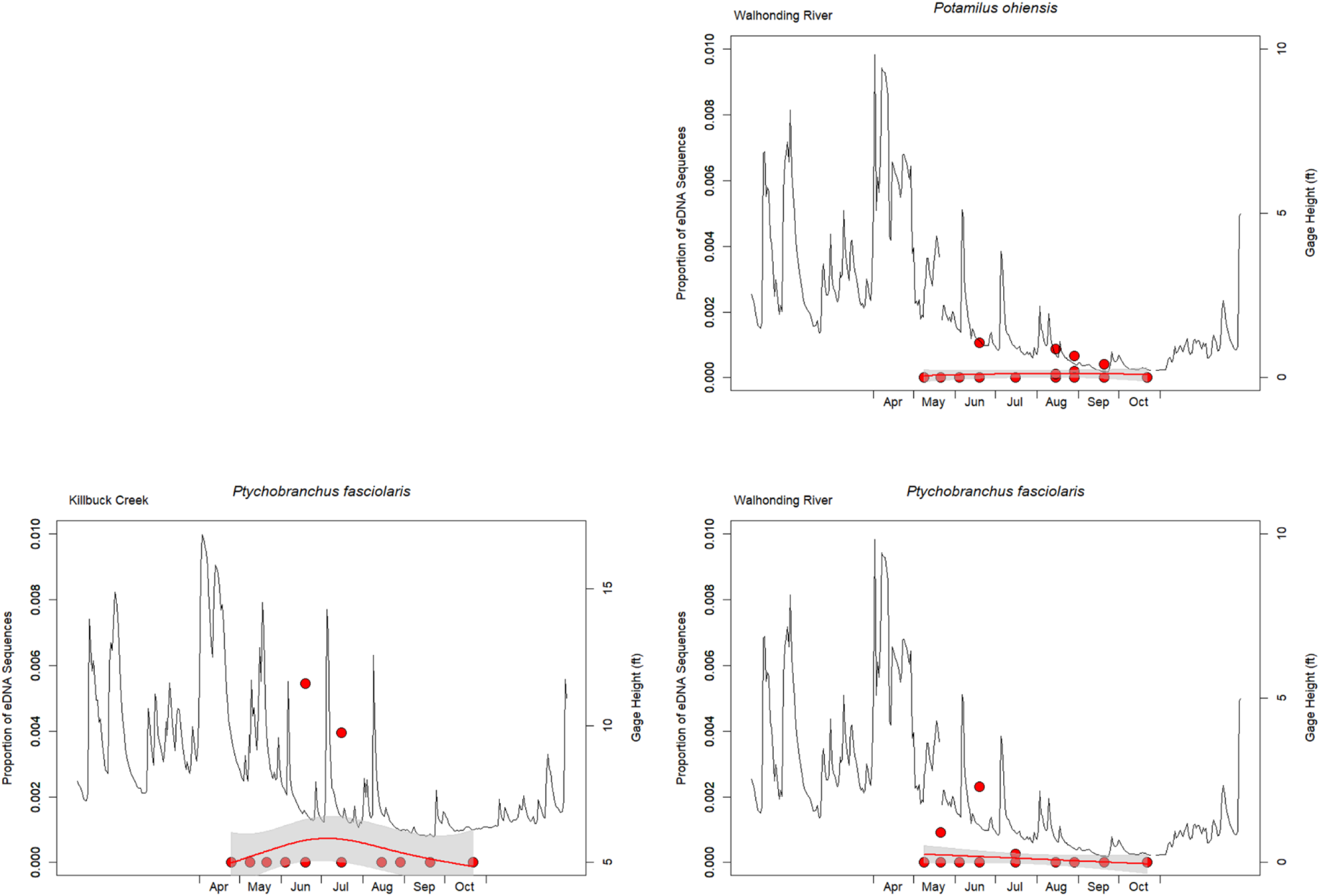
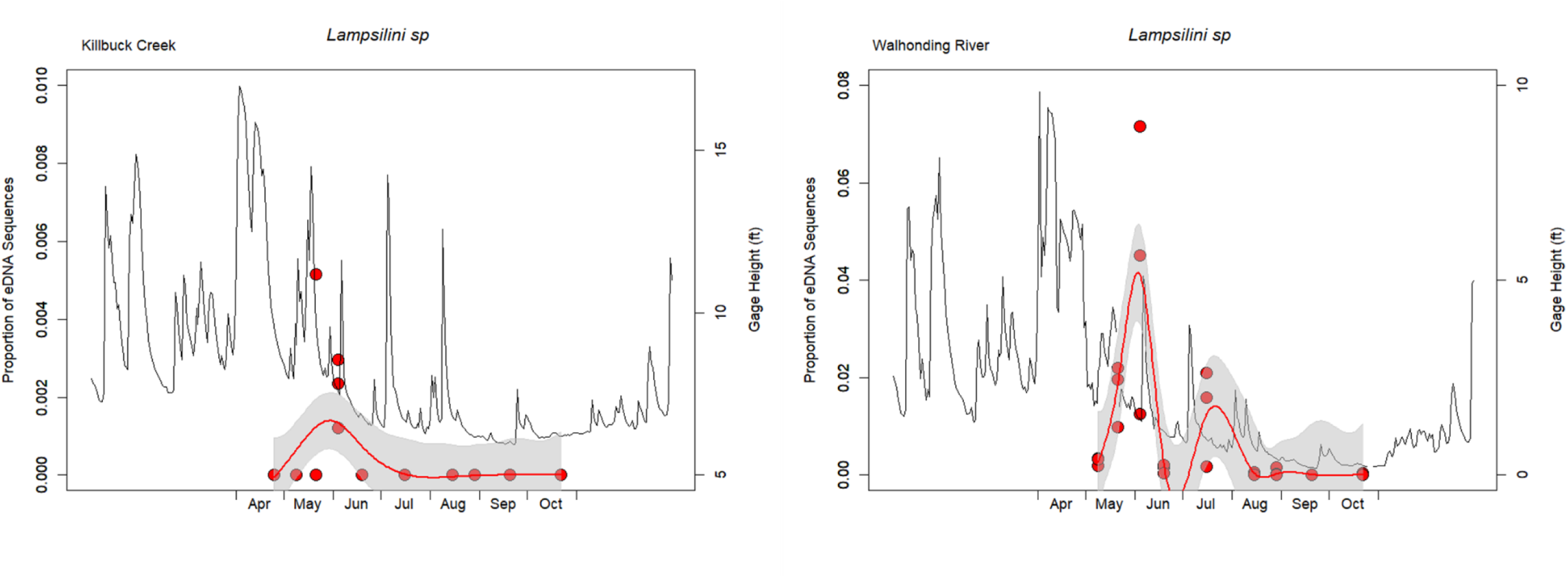
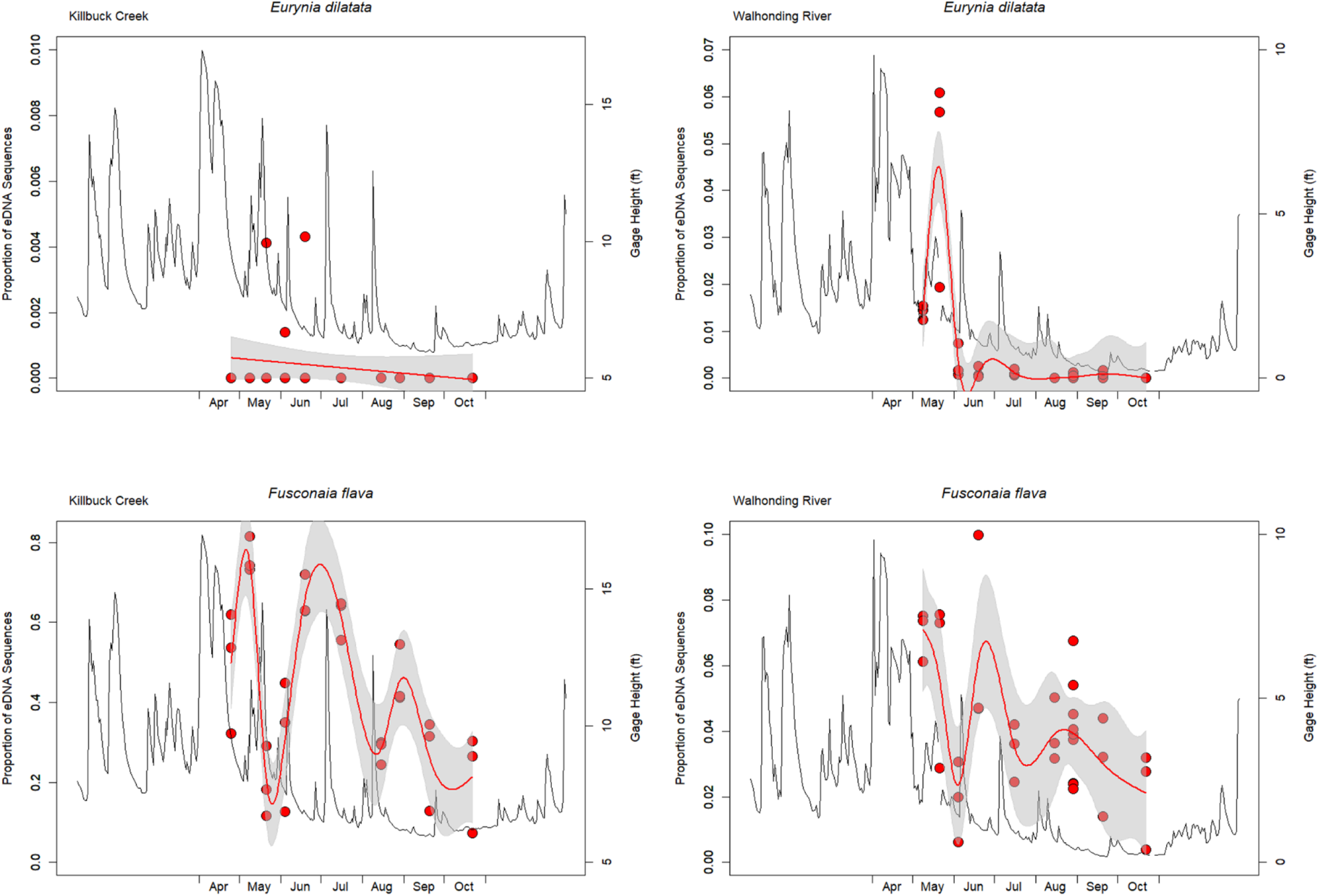
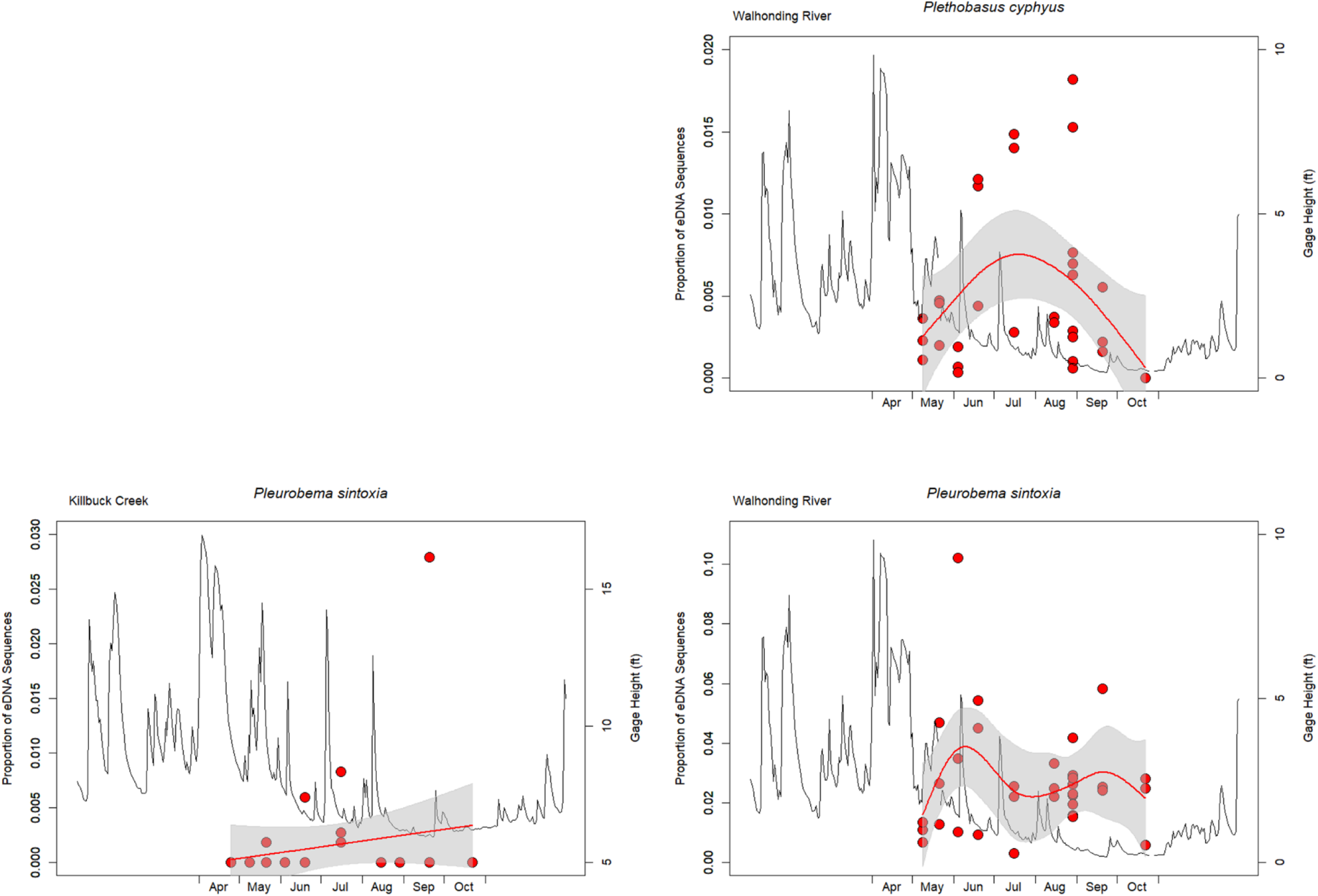
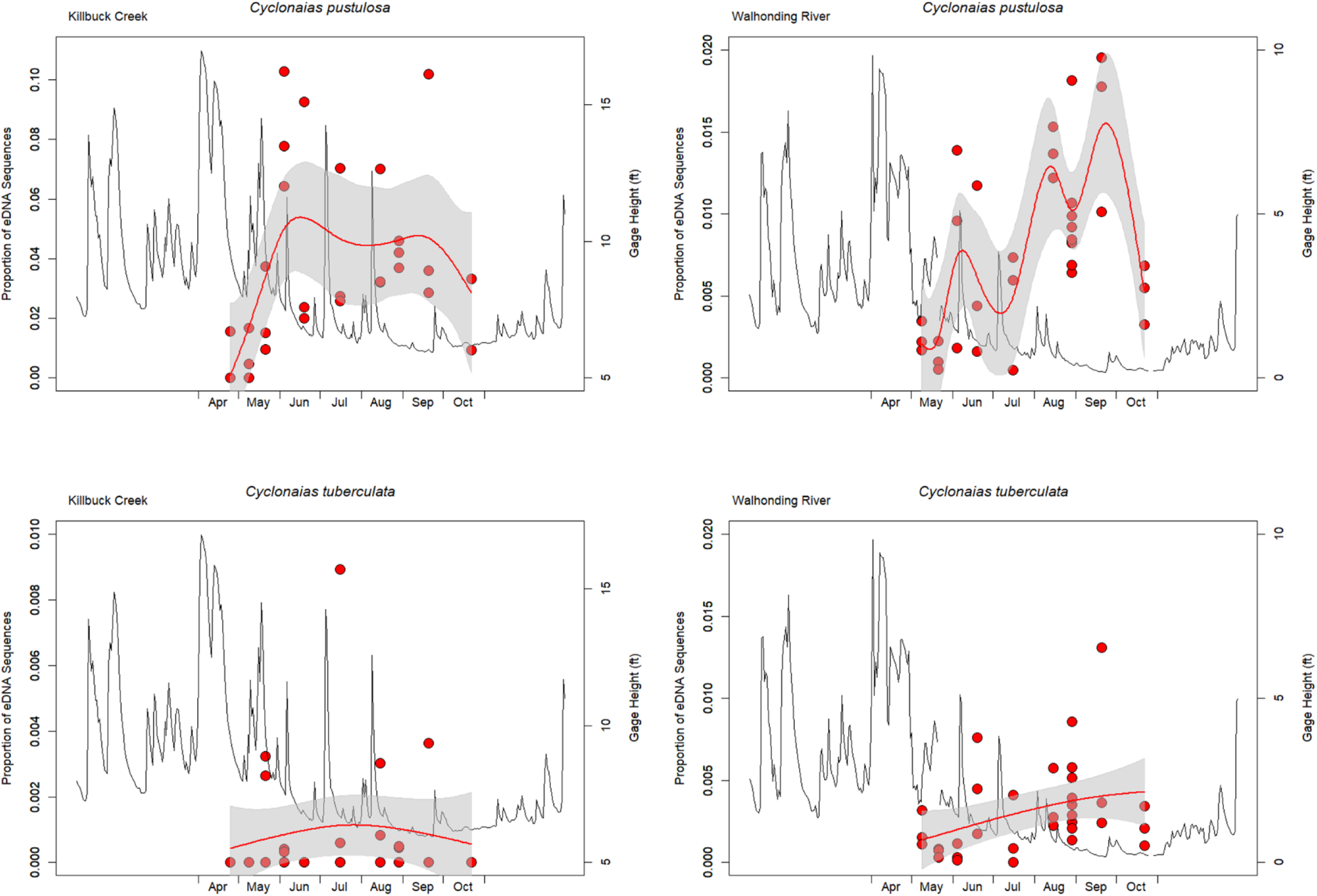
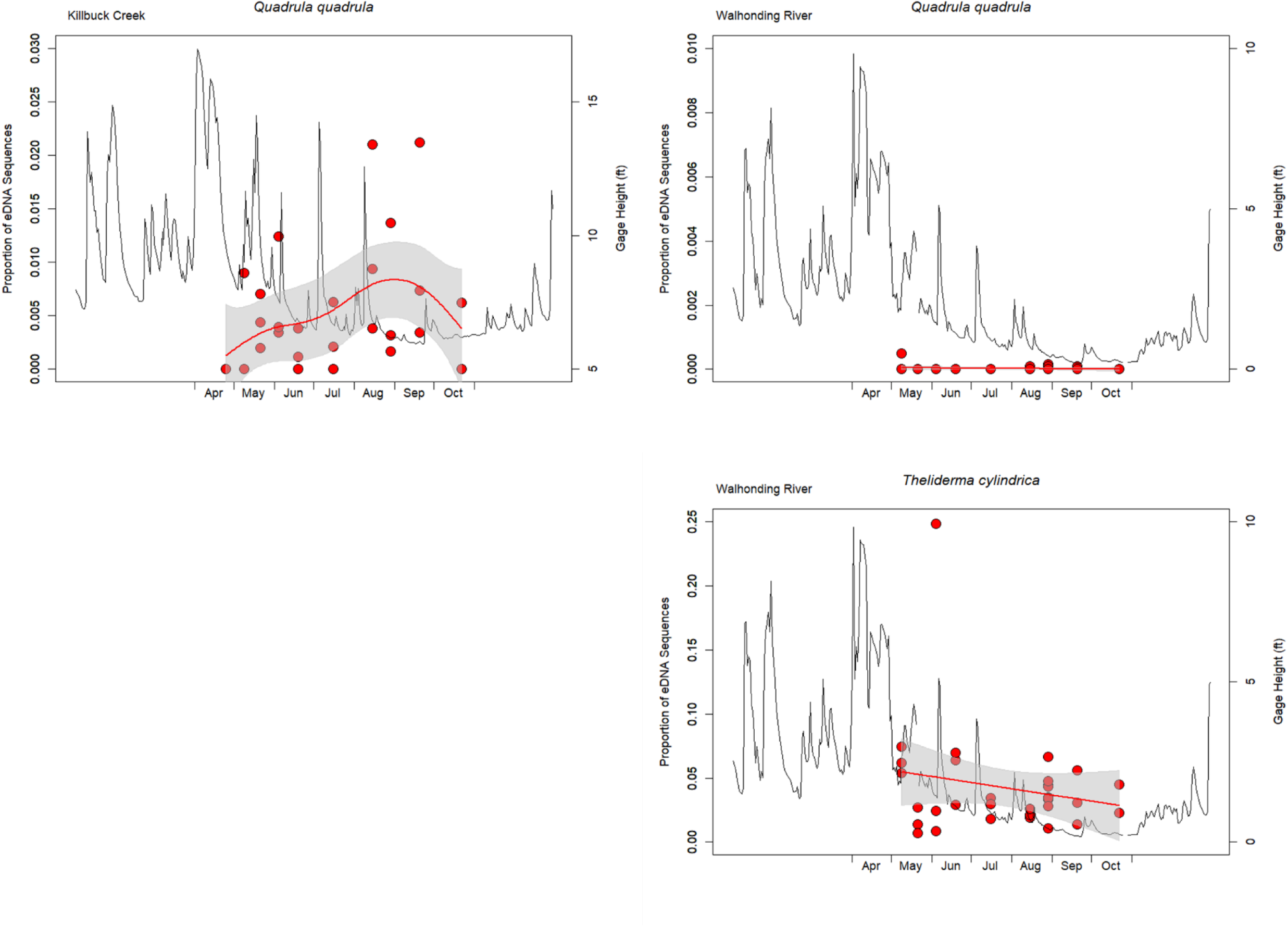
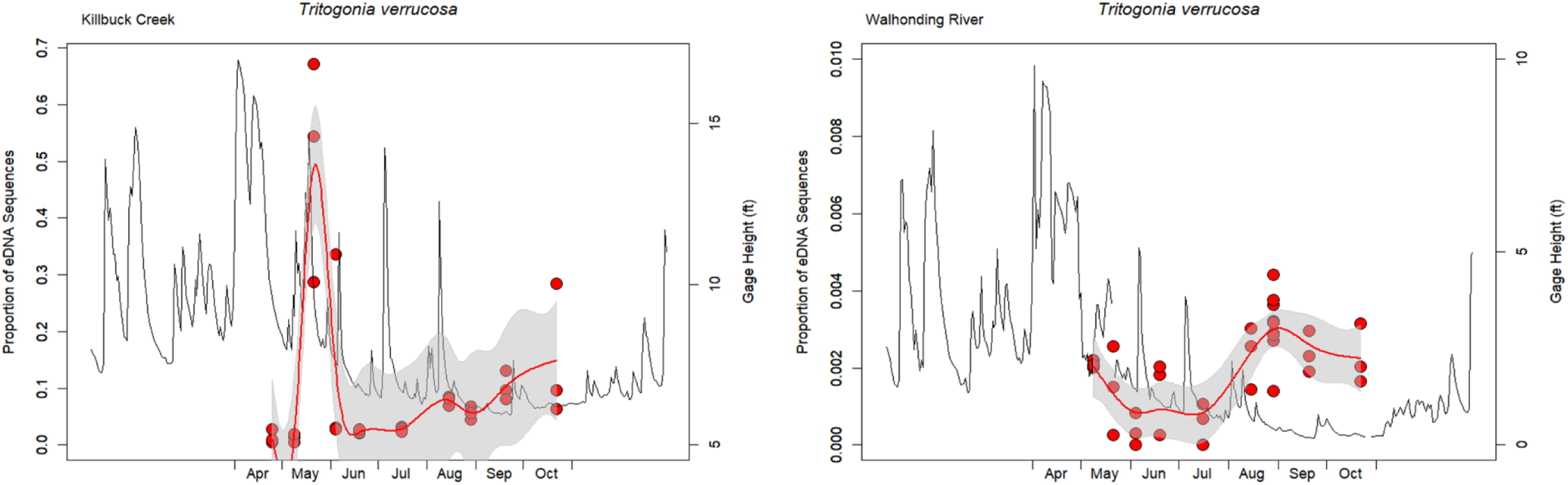
The proportion of the mussel assemblage (i.e., proportion of eDNA sequences) across all replicates on each sampling date from Killbuck Creek and Walhonding River for each species. Plots are organized by species and grouped within the major mussel tribes. The thin black line displays the USGS gage height (ft) across the sampling season.

**Supplementary Figure 6.**
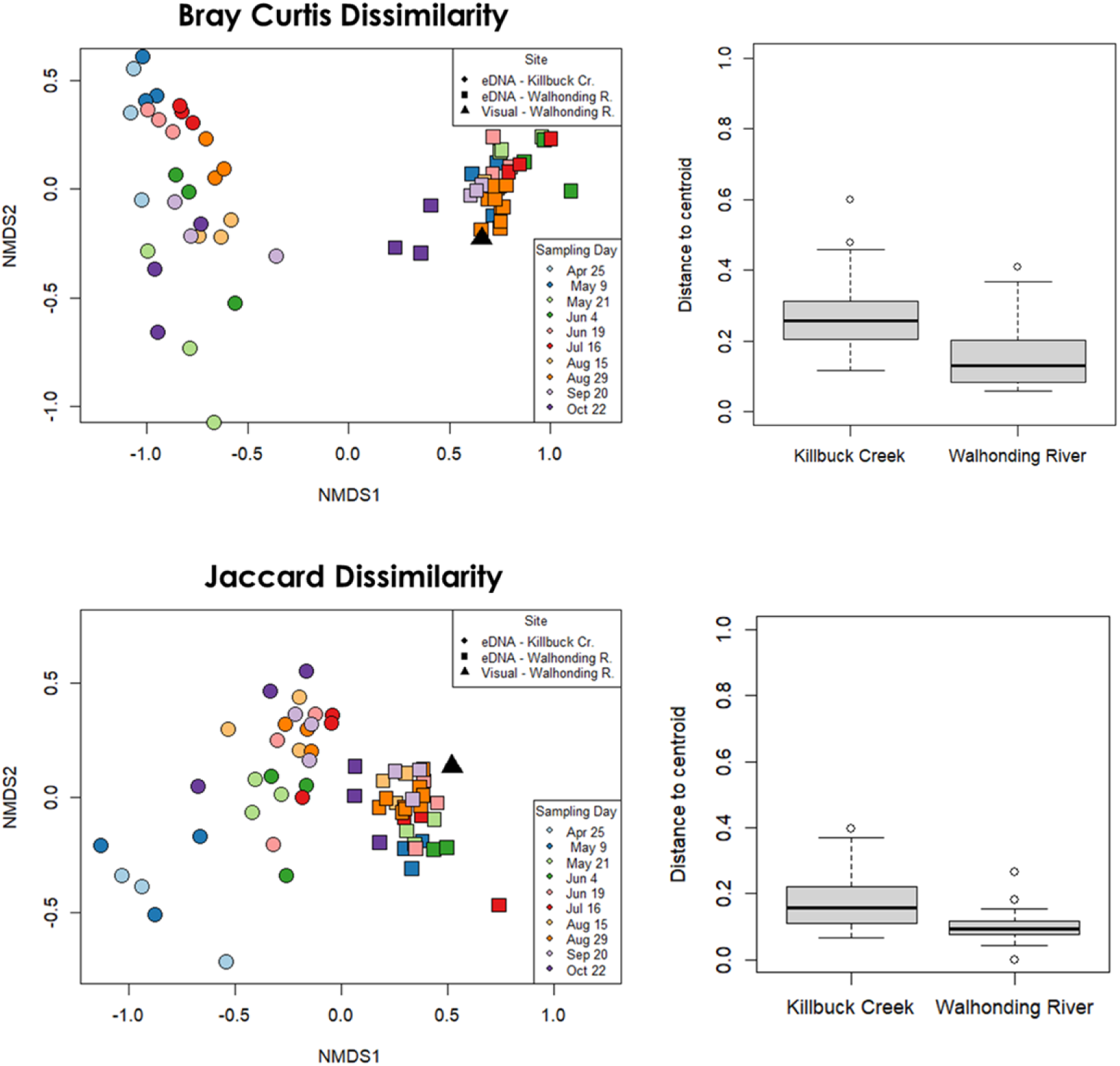
Non-metric multidimensional scaling plots for (A) Bray-Curtis Dissimilarity and (B) Jaccard Dissimilarity for the mussel assemblage detected with environmental DNA or with visual mussel survey.

**Supplementary Figure 7.**
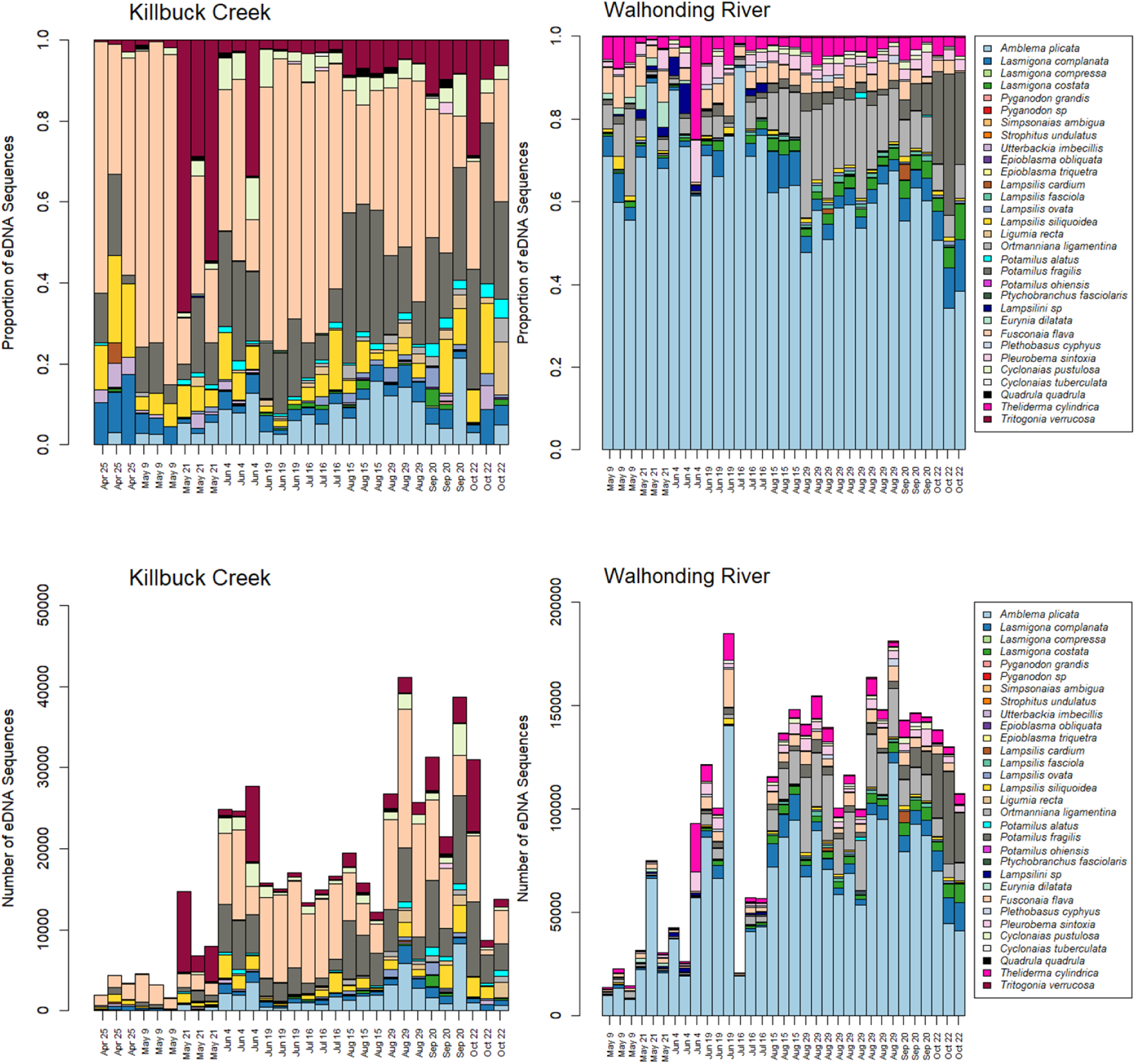
Number of freshwater mussel environmental DNA sequences and proportion of freshwater mussel environmental DNA sequences detected across sampling water replicates in Killbuck Creek and Walhonding River. The proportion of freshwater mussel abundance recorded during a visual survey in Walhonding River is shown.

**Supplementary Figure 8.**
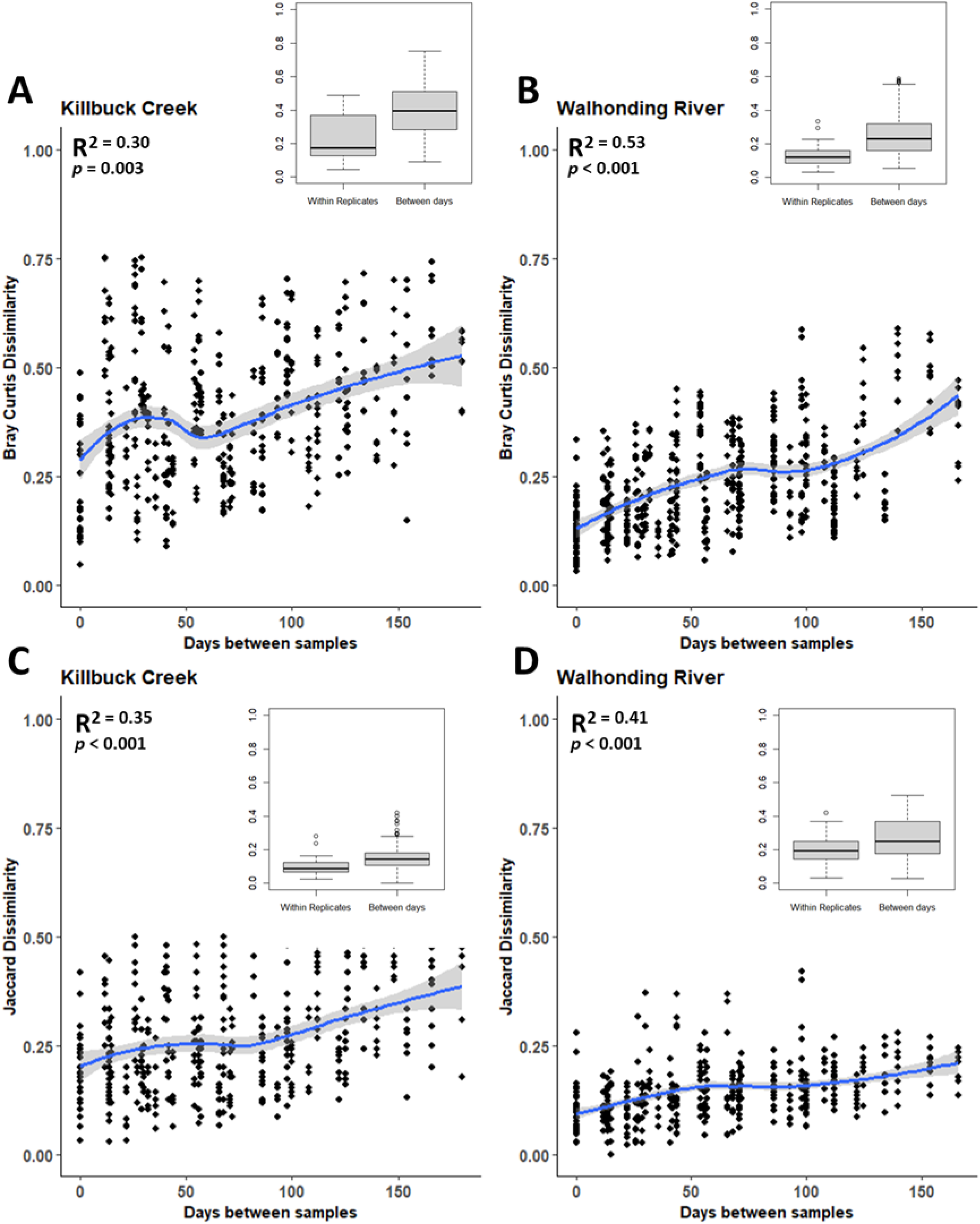
Beta diversity dissimilarities between environmental DNA samples for (A & B) Bray-Curtis Dissimilarity and (C & D) Jaccard Dissimilarity from (A & C) Killbuck Creek and (B & D) Walhonding River.

**Supplementary Figure 9.**
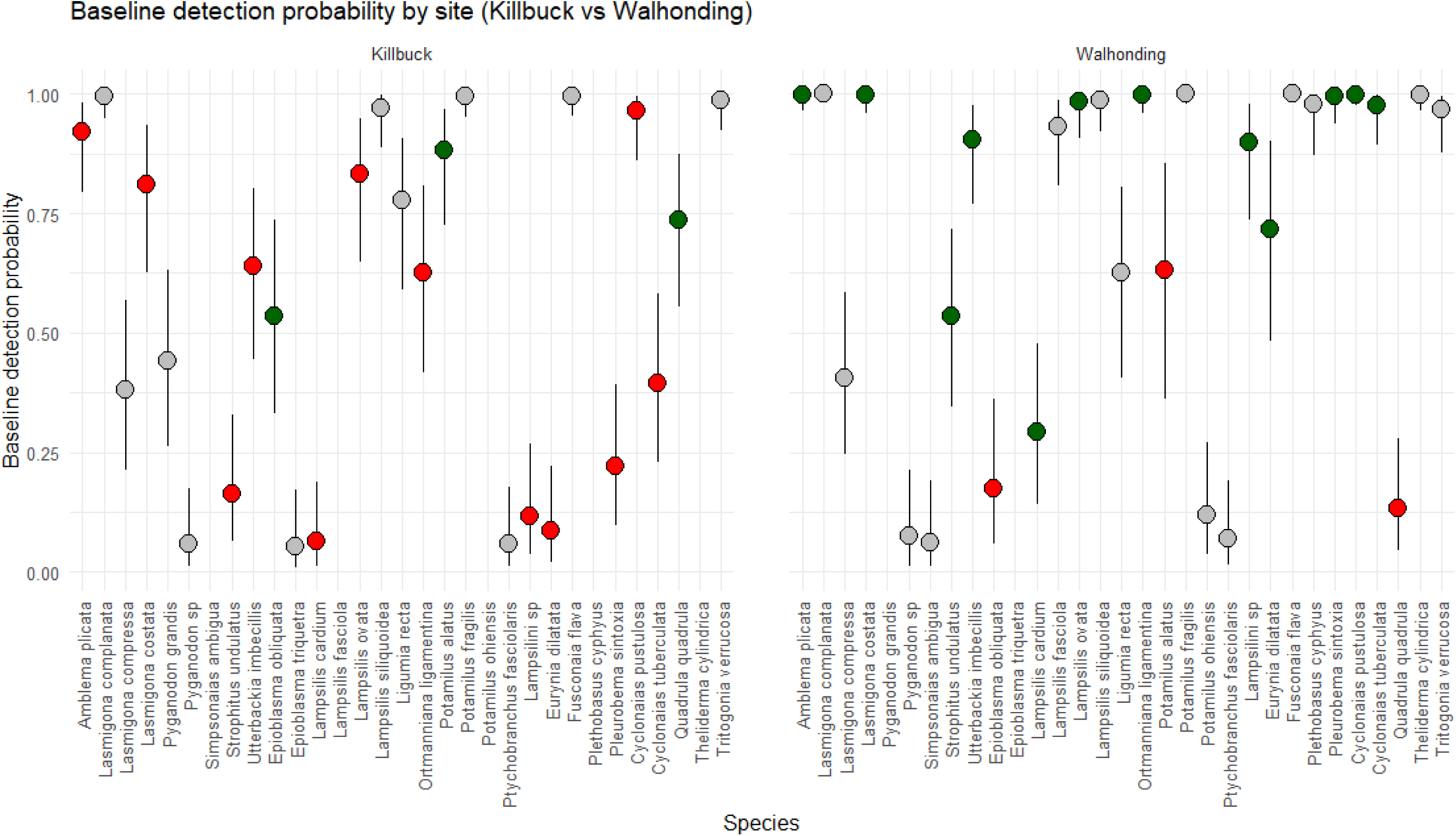
The median estimates of probability of environmental DNA detection for each detected species when collecting a single eDNA sample from (A) Killbuck Creek or (B) Walhonding River. Error bars represent credibility intervals for probability of detection estimates. Green points depict a species that had significantly greater detection probability at that corresponding site, while red depict a species that had significantly lower detection probability at that corresponding site

**Supplementary Figure 10.**
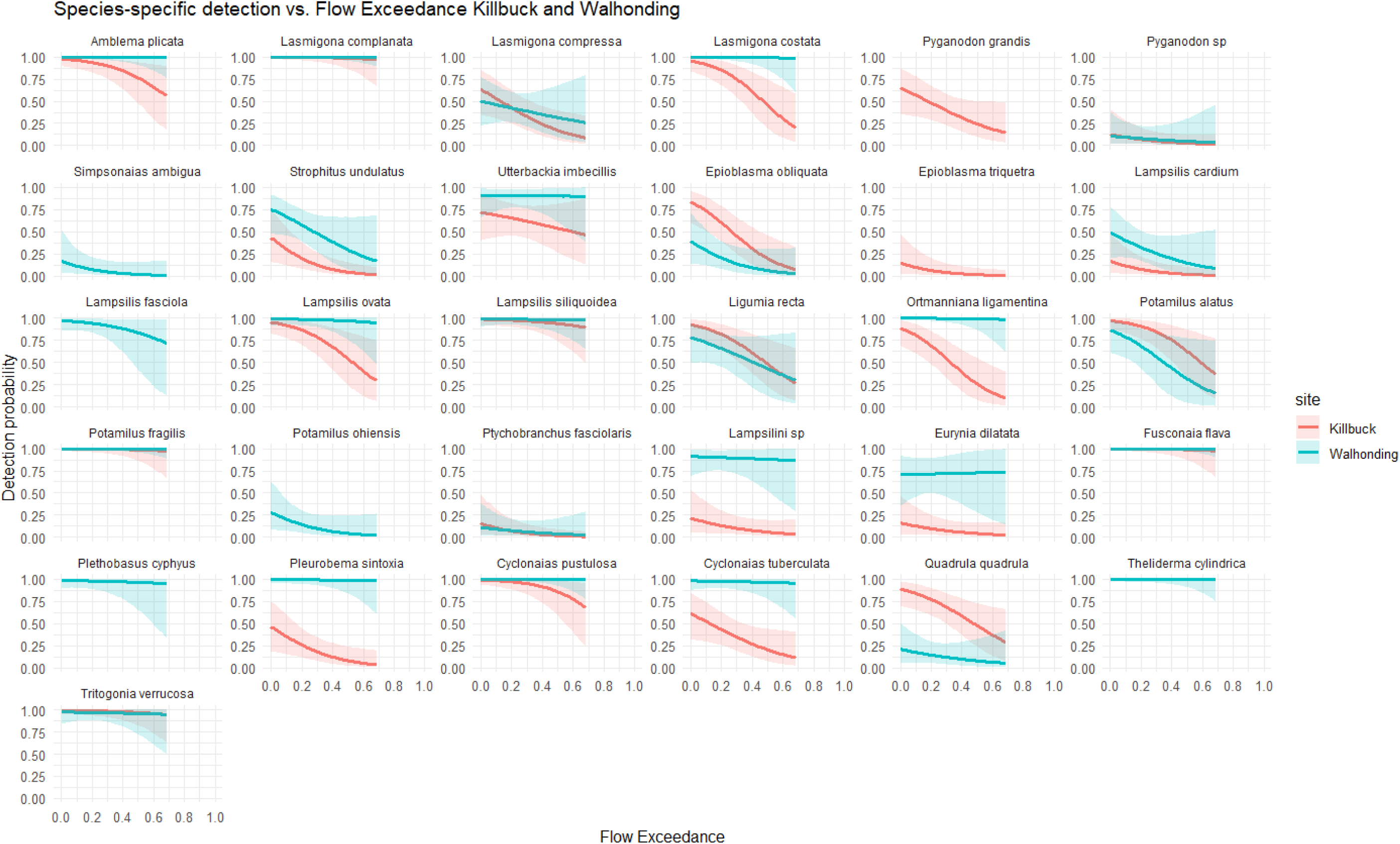
The estimated detection probability at Killbuck Creek or Walhonding River compared to flow exceedance (i.e., a measure of river discharge).

**Supplementary Figure 11.**
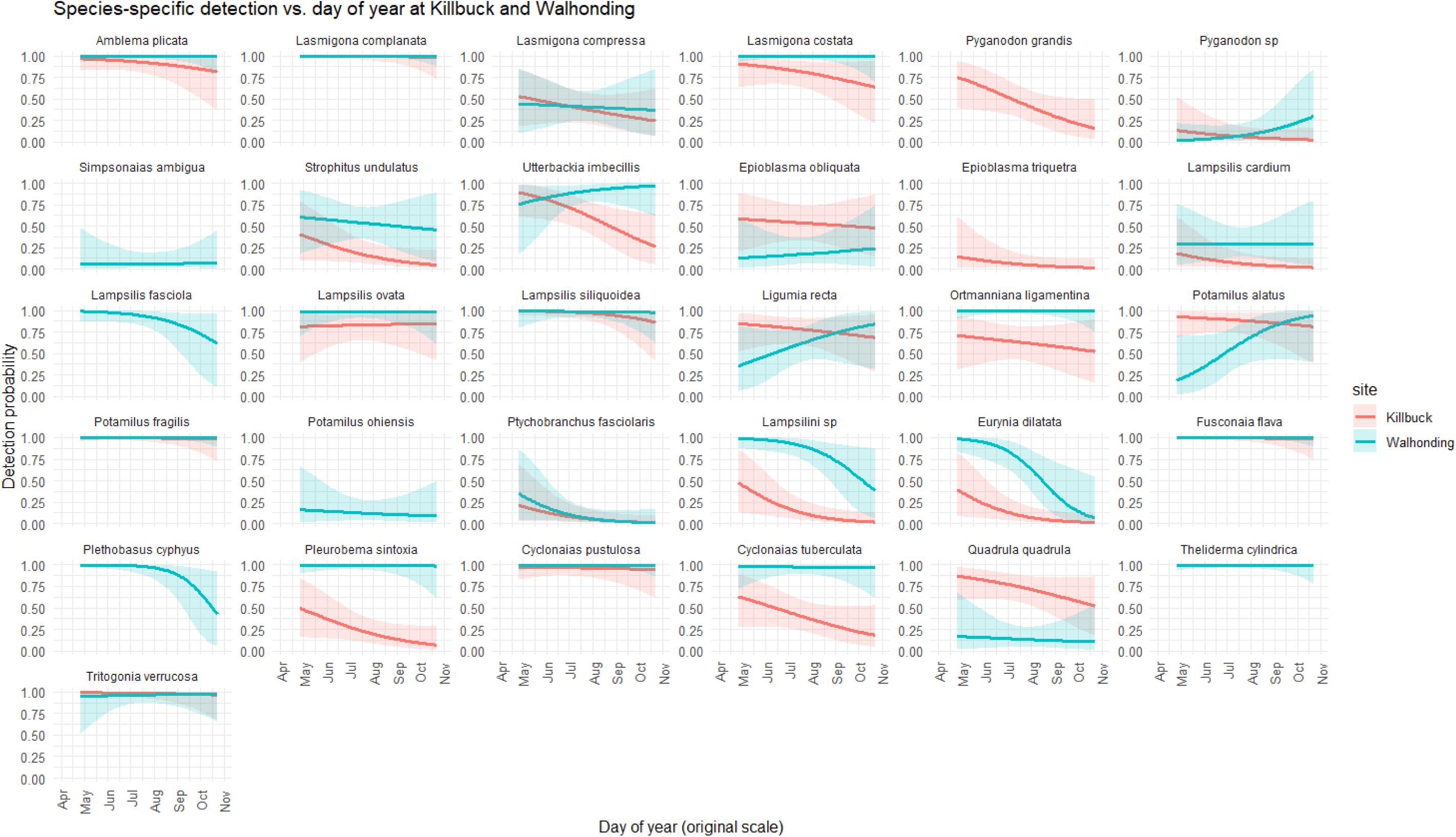
The estimated detection probability at Killbuck Creek or Walhonding River compared to the sampling season (e.g., day of the year).

**Supplementary Figure 12.**
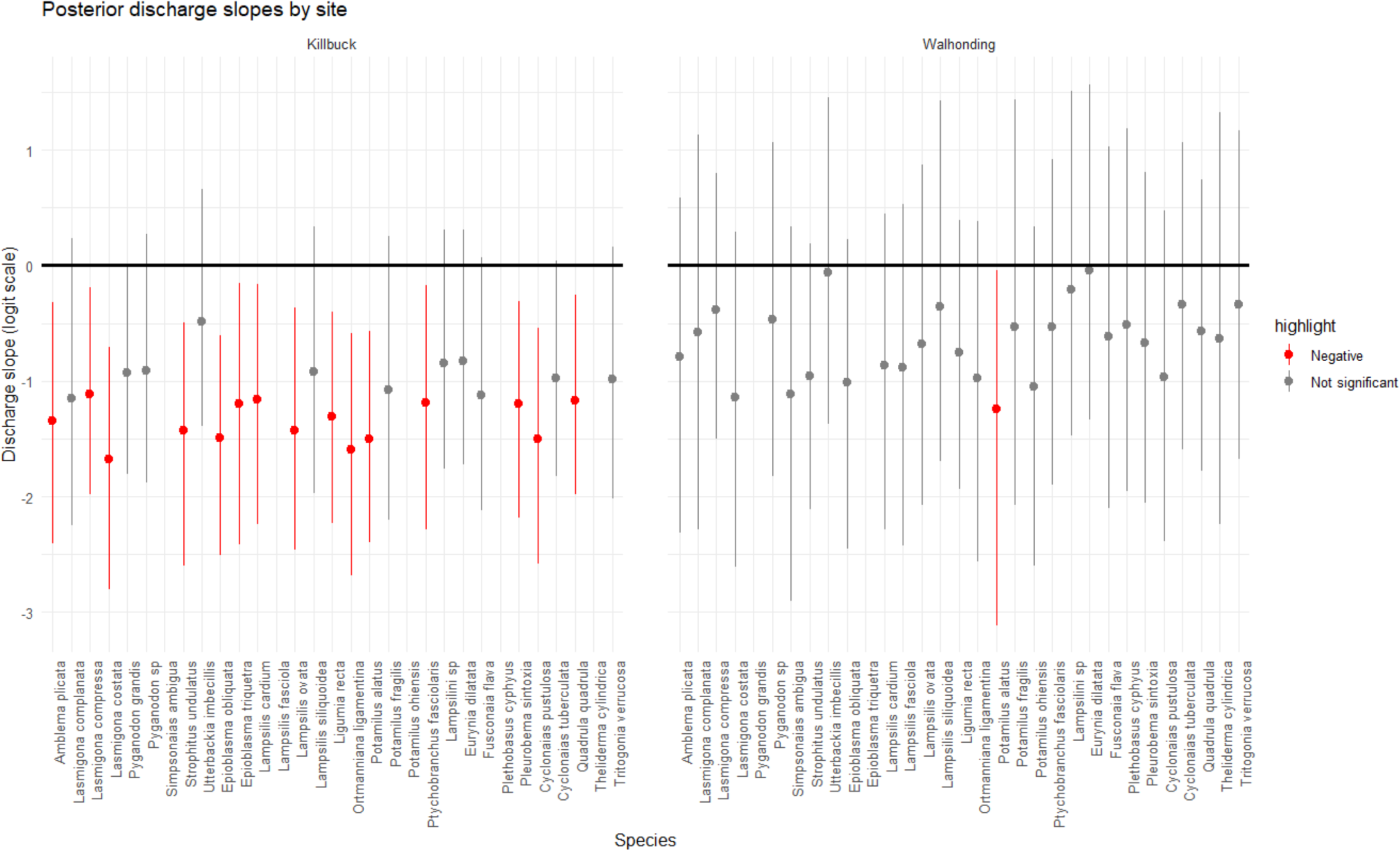
Posterior model estimates for detection probability related to flow exceedance (i.e., a measure of river discharge).

**Supplementary Figure 13.**
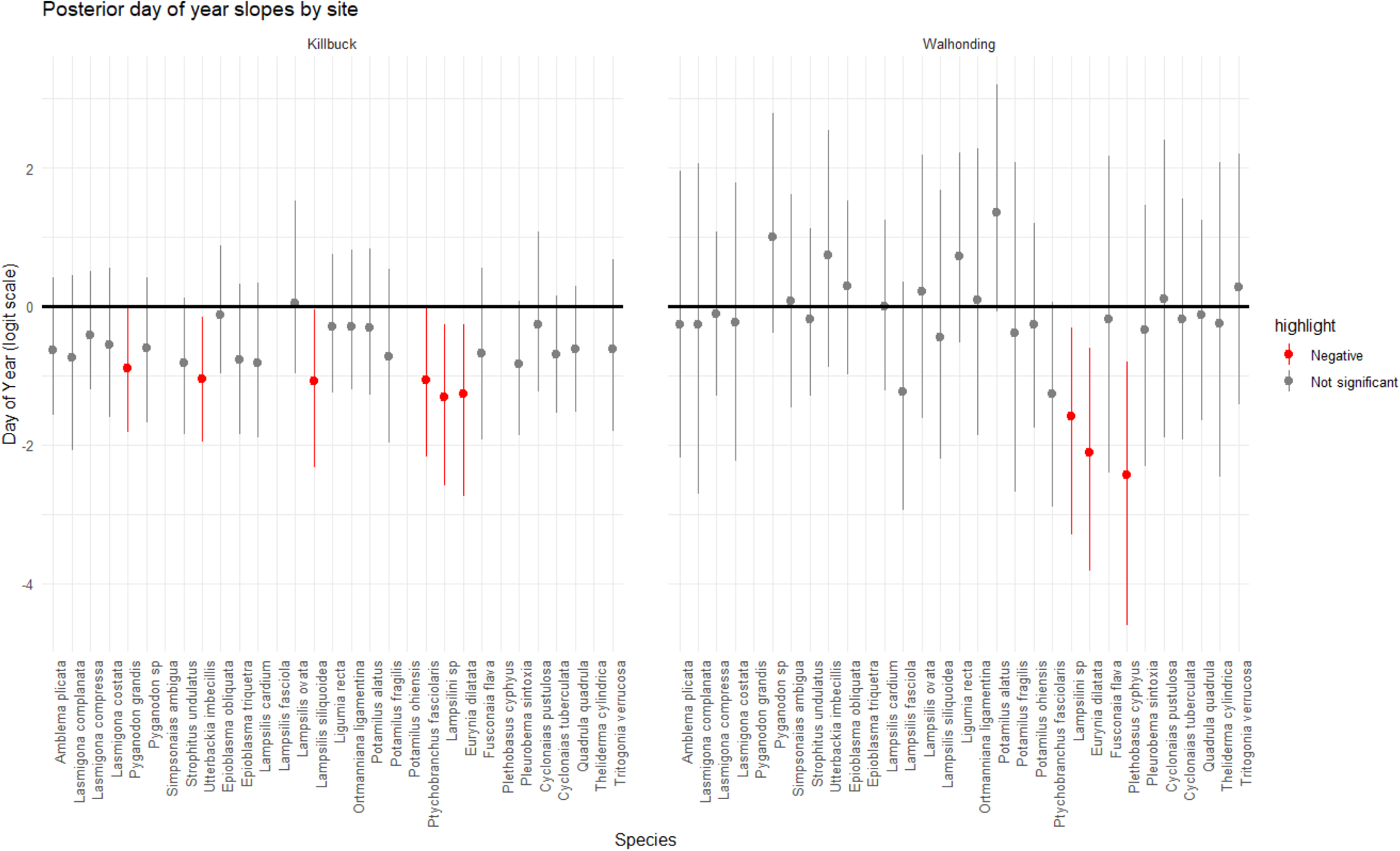
Posterior model estimates for detection probability related to sampling season (i.e., day of the year).

**Supplementary Figure 14.**
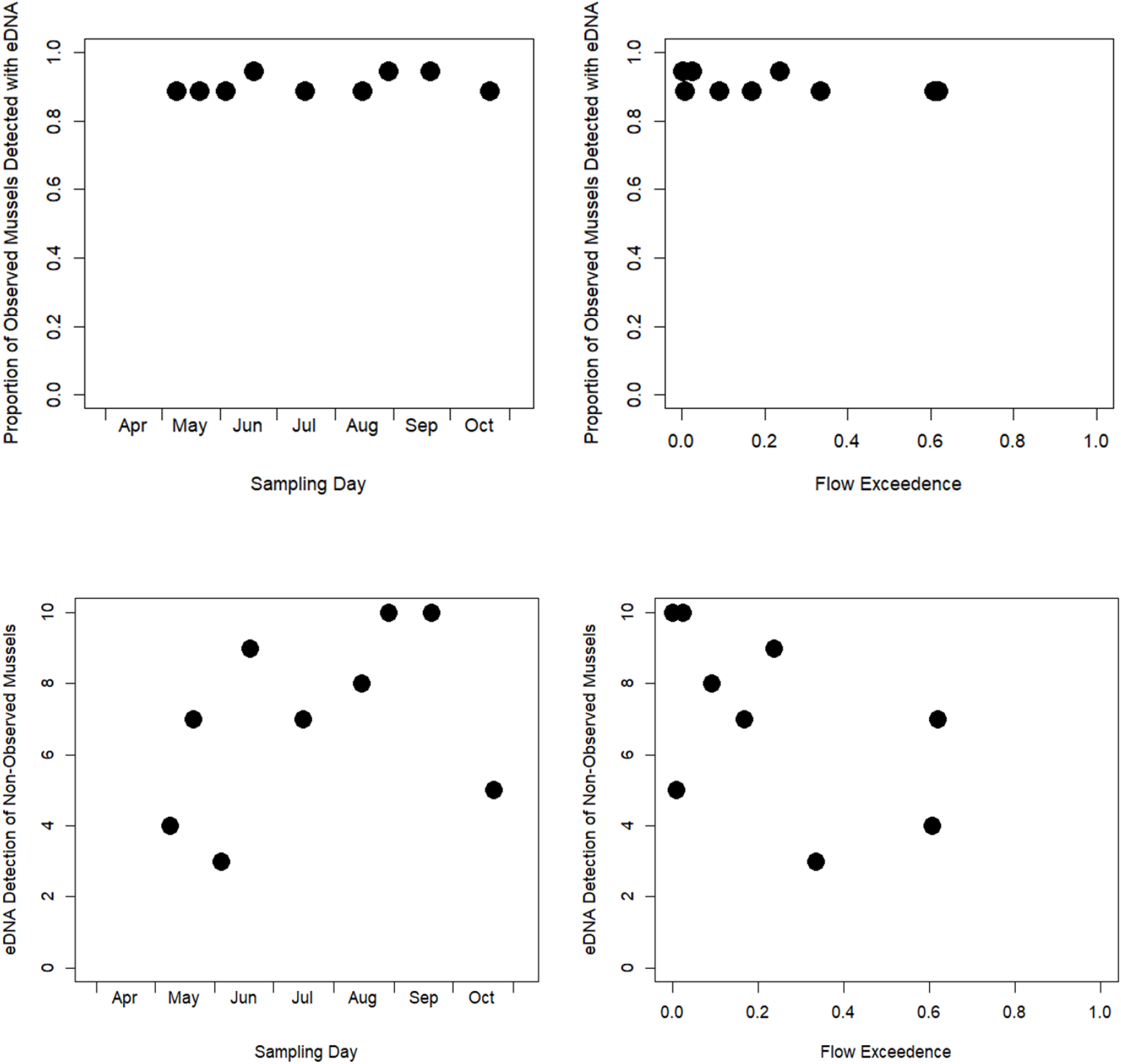
The proportion of visually observed mussel species detected with eDNA across the **(A) sampling season and (B) flow exceedances. The number of eDNA-only species detected across (C) sampling season and (D) flow exceedances.**

**Supplementary Figure 15.**
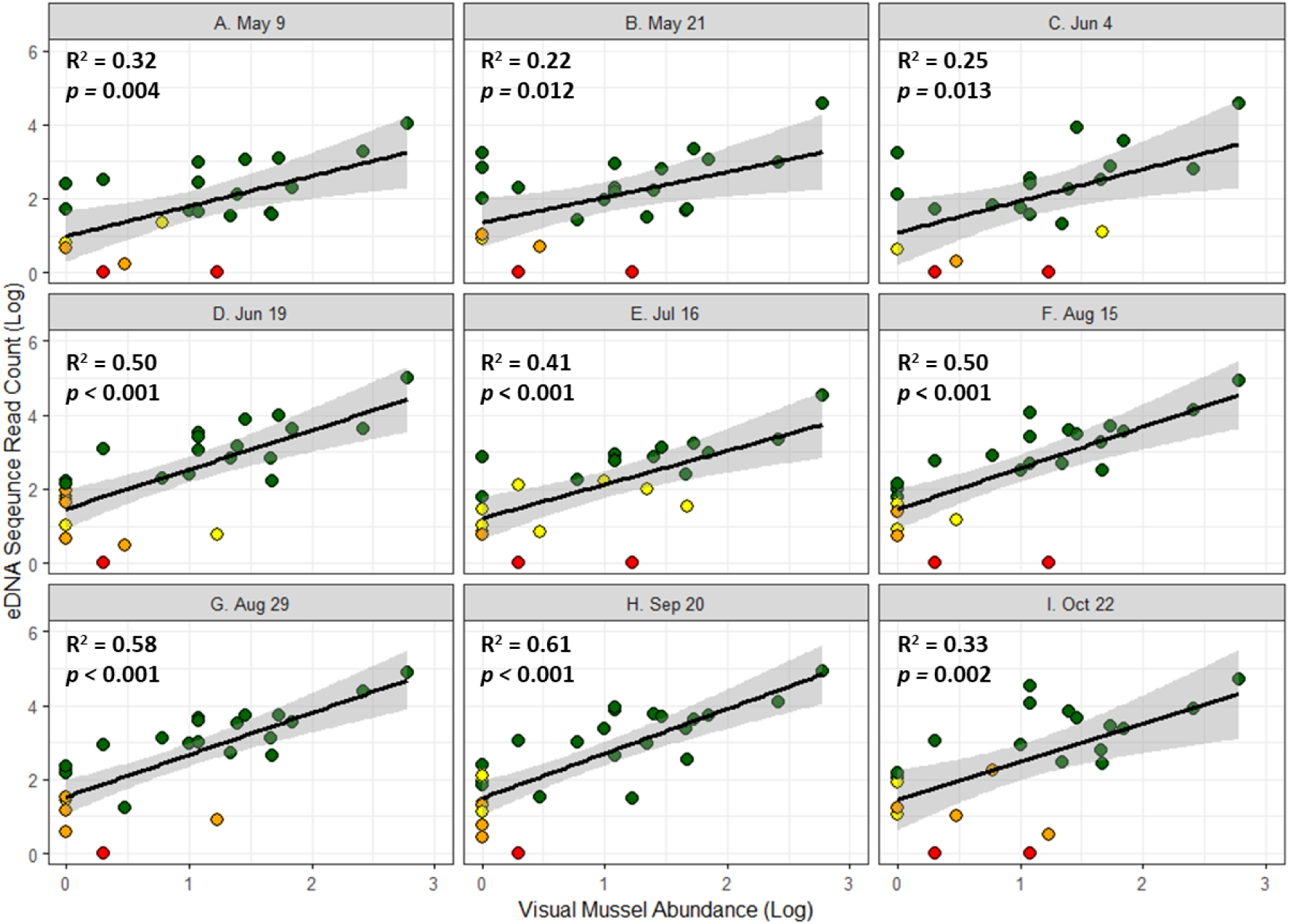
Linear regression plots comparing environmental DNA abundance (i.e., sequence read count) and visually observed mussel abundance for each individual sampling event in the Walhonding River.

**Supplementary Figure 16.**
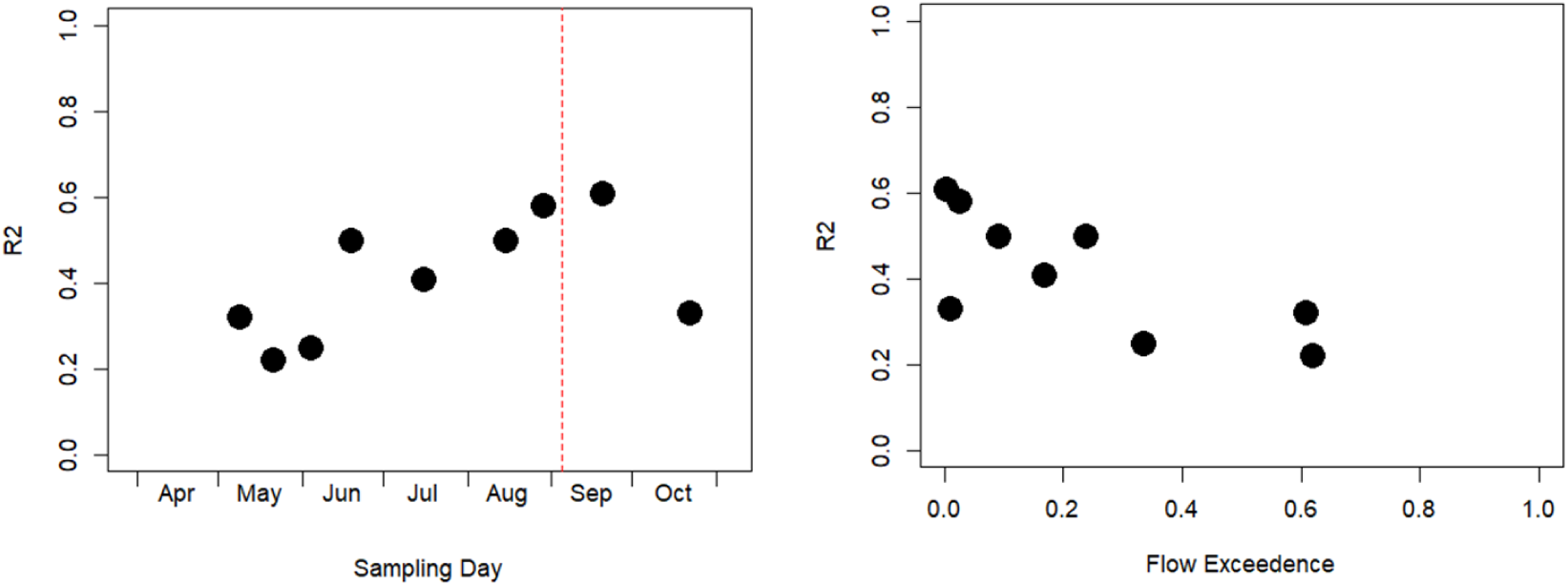
The correlation coefficient (R^2^) between environmental DNA abundance and visually observed abundance across sampling events and flow exceedances. Red dashed line indicates date of the visual mussel survey.

**Supplementary Figure 17.**
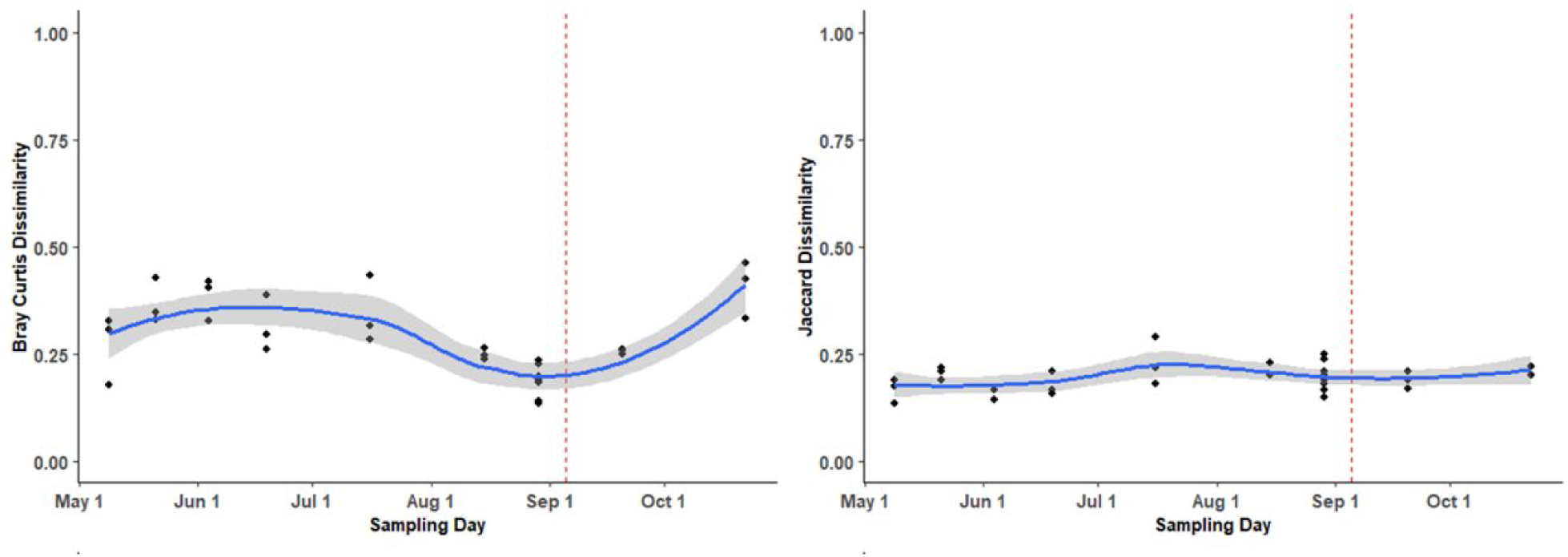
Beta diversity dissimilarity index (Bray-Curtis or Jaccard Dissimilarity) between environmental DNA and visually observed mussel assemblage compared across sampling events in the Walhonding River. Red dashed line indicates date of the visual mussel survey.

**Supplementary Figure 18.**
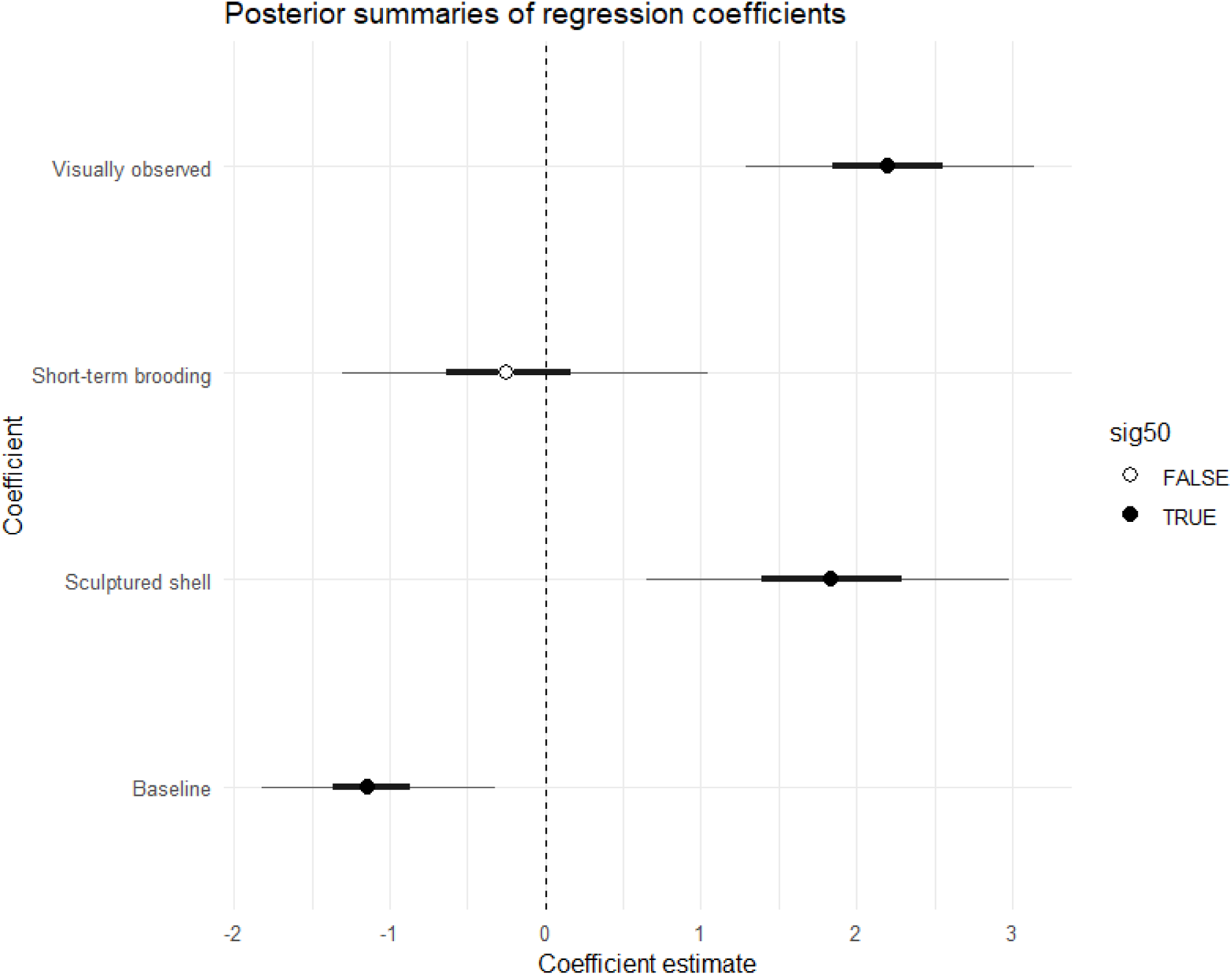
Posterior model estimates from the Walhonding River for their influence on eDNA detection probability based on if a mussel was visually observed at a site, if a species is a shot-term brooder, or if a species has a sculptured shell.

**Supplementary Table 1.** Environmental DNA sample metadata for each sampling event in Killbuck Creek and Walhonding River. A measure of flow exceedance was generated by calculating the exceedance probability on each sampling data, where the exceedance probability is the total number of days in which the gage height was higher than that on the sampling date divided by the total number of days on record for that gage.

| **Sampling Site** | **Sampling Trip** | **Replicates** | **Date** | **Julian Day of**<br>**Year** | **Gage Height**<br>**(ft)** | **Flow**<br>**Exceedance** |
| --- | --- | --- | --- | --- | --- | --- |
| Killbuck Cr. | 1 | 3 | 04252024 | 116 | 9.58 | 0.675 |
| Killbuck Cr. | 2 | 3 | 05092024 | 130 | 9.02 | 0.617 |
| Killbuck Cr. | 3 | 3 | 05212024 | 142 | 9.69 | 0.686 |
| Killbuck Cr. | 4 | 3 | 06042024 | 156 | 7.59 | 0.374 |
| Killbuck Cr. | 5 | 3 | 06192024 | 171 | 6.84 | 0.216 |
| Killbuck Cr. | 6 | 3 | 07162024 | 198 | 6.69 | 0.175 |
| Killbuck Cr. | 7 | 3 | 08152024 | 228 | 6.82 | 0.207 |
| Killbuck Cr. | 8 | 3 | 08292024 | 242 | 6.17 | 0.033 |
| Killbuck Cr. | 9 | 3 | 09202024 | 264 | 6.04 | 0.019 |
| Killbuck Cr. | 10 | 3 | 10222024 | 296 | 6.20 | 0.035 |
| Walhonding R. | 2 | 3 | 05092024 | 130 | 2.66 | 0.608 |
| Walhonding R. | 3 | 3 | 05212024 | 142 | 2.67 | 0.619 |
| Walhonding R. | 4 | 3 | 06042024 | 156 | 1.47 | 0.335 |
| Walhonding R. | 5 | 3 | 06192024 | 171 | 1.13 | 0.237 |
| Walhonding R. | 6 | 3 | 07162024 | 198 | 0.92 | 0.167 |
| Walhonding R. | 7 | 3 | 08152024 | 228 | 0.70 | 0.091 |
| Walhonding R. | 8 | 9 | 08292024 | 242 | 0.43 | 0.025 |
| Walhonding R. | 9 | 3 | 09202024 | 264 | 0.20 | 0.002 |
| Walhonding R. | 10 | 3 | 10222024 | 296 | 0.24 | 0.009 |

## References

1. Amyot, J. P., & Downing, J. A. (1991). Endo-and epibenthic distribution of the unionid mollusc *Elliptio complanata*. Journal of the North American Benthological Society, 10(3), 280–285.

2. Amyot, J. P., & Downing, J. (1997). Seasonal variation in vertical and horizontal movement of the freshwater bivalve *Elliptio complanata* (Mollusca: Unionidae). Freshwater Biology, 37(2), 345–354.

3. Beng, K. C., & Corlett, R. T. (2020). Applications of environmental DNA (eDNA) in ecology and conservation: opportunities, challenges and prospects. Biodiversity and Conservation, 29, 2089–2121.

4. Bolyen, E., Rideout, J. R., Dillon, M. R., Bokulich, N. A., Abnet, C. C., Al-Ghalith, G. A., … & Caporaso, J. G. (2019). Reproducible, interactive, scalable and extensible microbiome data science using QIIME 2. Nature biotechnology, 37(8), 852–857.

5. Callahan, B. J., McMurdie, P. J., Rosen, M. J., Han, A. W., Johnson, A. J. A., & Holmes, S. P. (2016). DADA2: High-resolution sample inference from Illumina amplicon data. Nature methods, 13(7), 581–583.

6. Camacho, C., Coulouris, G., Avagyan, V., Ma, N., Papadopoulos, J., Bealer, K., & Madden, T. L. (2009). BLAST+: architecture and applications. BMC bioinformatics, 10, 1–9.

7. Curole, J. P., & Kocher, T. D. (2005). Evolution of a unique mitotype-specific protein-coding extension of the cytochrome c oxidase II gene in freshwater mussels (Bivalvia: Unionoida). Journal of Molecular Evolution, 61(3), 381–389.

8. Curtis, A. N., Tiemann, J. S., Douglass, S. A., Davis, M. A., & Larson, E. R. (2021). High stream flows dilute environmental DNA (eDNA) concentrations and reduce detectability. Diversity and Distributions, 27(10), 1918–1931.

9. Deiner, K., & Altermatt, F. (2014). Transport distance of invertebrate environmental DNA in a natural river. PloS one, 9(2), e88786.

10. Deiner, K., Yamanaka, H., & Bernatchez, L. (2021). The future of biodiversity monitoring and conservation utilizing environmental DNA. Environmental DNA, 3(1), 3–7.

11. De Souza, L. S., Godwin, J. C., Renshaw, M. A., & Larson, E. (2016). Environmental DNA (eDNA) detection probability is influenced by seasonal activity of organisms. PloS one, 11(10), e0165273.

12. EDGE Edge Engineering & Science, LLC (2025). Purple Catspaw Monitoring – Walhonding River Purple Catspaw Monitoring Survey. Coshocton County, Ohio. Submitted to USFWS March 2025.

13. Esling, P., Lejzerowicz, F., & Pawlowski, J. (2015). Accurate multiplexing and filtering for high-throughput amplicon-sequencing. Nucleic acids research, 43(5), 2513–2524. 10.1093/nar/gkv107

14. Fediajevaite, J., Priestley, V., Arnold, R., & Savolainen, V. (2021). Meta-analysis shows that environmental DNA outperforms traditional surveys, but warrants better reporting standards. Ecology and Evolution, 11(9), 4803–4815.

15. Fiske, I., & Chandler, R. (2011). Unmarked: an R package for fitting hierarchical models of wildlife occurrence and abundance. Journal of statistical software, 43, 1–23.

16. FMCS Freshwater Mollusk Conservation Society. 2021. The 2021 checklist of freshwater bivalves (Mollusca: Bivalvia: Unionida) of the United States and Canada. Considered and approved by the Bivalve Names Subcommittee December 2020.

17. Gusman, A., Lecomte, S., Stewart, D. T., Passamonti, M., & Breton, S. (2016). Pursuing the quest for better understanding the taxonomic distribution of the system of doubly uniparental inheritance of mtDNA. PeerJ, 4, e2760.

18. Haag, W. R. (2012). North American freshwater mussels: natural history, ecology, and conservation. Cambridge University Press.

19. Hornbach, D. J., & Deneka, T. (1996). A comparison of a qualitative and a quantitative collection method for examining freshwater mussel assemblages. Journal of the North American Benthological Society, 15(4), 587–596.

20. Jerde, C. L. (2021). Can we manage fisheries with the inherent uncertainty from eDNA?. Journal of fish biology, 98(2), 341–353.

21. Laschever, E., Kelly, R., Hoge, M., & Lee, K. (2023). The next generation of environmental monitoring: Environmental DNA in agency practice. Columbia Journal of Environmental Law, 48(S), 51–51.

22. MacKenzie, D. I., Nichols, J. D., Lachman, G. B., Droege, S., Andrew Royle, J., & Langtimm, C. A. (2002). Estimating site occupancy rates when detection probabilities are less than one. Ecology, 83(8), 2248–2255.

23. Marshall, N. T., Symonds, D. E., Dean, C. A., Schumer, G., & Fleece, W. C. (2022). Evaluating environmental DNA metabarcoding as a survey tool for unionid mussel assessments. Freshwater Biology, 67(9), 1483–1507.

24. Marshall, N. T., & Fleece, W. C. (2025). Assessment of Environmental DNA Survey Design for the Detection of Freshwater Unionid Mussels. Environmental DNA, 7(4), e70152.

25. Martin, M. (2011). Cutadapt removes adapter sequences from high-throughput sequencing reads. EMBnet. journal, 17(1), 10–12.

26. O’Donnell, J. L., Kelly, R. P., Lowell, N. C., & Port, J. A. (2016). Indexed PCR primers induce template-specific bias in large-scale DNA sequencing studies. PloS one, 11(3), e0148698.

27. ODNR Division of Wildlife and USFWS Ohio Ecological Services Field Office. 2025. Ohio Mussel Survey Protoco.l April 2025. 63 pp. https://dam.assets.ohio.gov/image/upload/ohiodnr.gov/documents/wildlife/permits/dow-protocol-ohio-mussel-survey.pdf

28. Perles, S. J., Christian, A. D., & Berg, D. J. (2003). Vertical migration, orientation, aggregation, and fecundity of the freshwater mussel *Lampsilis siliquoidea*. The Ohio Journal of Science. v103, n4 (September, 2003), 73-78.

29. Prié, V., Valentini, A., Lopes-Lima, M., Froufe, E., Rocle, M., Poulet, N., … & Dejean, T. (2021). Environmental DNA metabarcoding for freshwater bivalves biodiversity assessment: methods and results for the Western Palearctic (European sub-region). Hydrobiologia, 848, 2931–2950.

30. Prié, V., Clément, L., Gaboriaud, C., Gailledrat, M., Naudon, D., Bourri, R., Lopes-Lima, M., & Valentini, A. (2025). From the Balkans to France: the threatened range-restricted freshwater mussel Unio carneus revealed as an introduced species by environmental DNA.

31. Reid, S. M. (2016). Search effort and imperfect detection: Influence on timed-search mussel (Bivalvia: Unionidae) surveys in Canadian rivers. Knowledge and Management of Aquatic Ecosystems, (417), 17.

32. Richardson, R. T. (2022). Controlling critical mistag-associated false discoveries in metagenetic data. Methods in Ecology and Evolution, 13(5), 938–944. 10.1111/2041-210X.13838

33. Ruppert, K. M., Kline, R. J., & Rahman, M. S. (2019). Past, present, and future perspectives of environmental DNA (eDNA) metabarcoding: A systematic review in methods, monitoring, and applications of global eDNA. Global Ecology and Conservation, 17, e00547.

34. Sanchez, B., & Schwalb, A. N. (2021). Detectability affects the performance of survey methods: a comparison of sampling methods of freshwater mussels in Central Texas. Hydrobiologia, 848(12), 2919–2929.

35. Schill, W. B., & Galbraith, H. S. (2019). Detecting the undetectable: Characterization, optimization, and validation of an eDNA detection assay for the federally endangered dwarf wedgemussel, Alasmidonta heterodon (Bivalvia: Unionoida). Aquatic conservation, 29(4).

36. Schwalb, A. N., & Pusch, M. T. (2007). Horizontal and vertical movements of unionid mussels in a lowland river. Journal of the North American Benthological Society, 26(2), 261–272.

37. Sellers, G. S., Jerde, C. L., Harper, L. R., Benucci, M., Di Muri, C., Li, J., Peirson, G., Walsh, K., Hatton-Ellis, T., Duncan, W., Duguid, A., Ottewell, D., Willby, N., Law, A., Bean, C.W., Winfield, I.J., Read, D.S., Handley, L.L., & Hänfling, B. (2024). Optimising species detection probability and sampling effort in lake fish eDNA surveys. Metabarcoding and Metagenomics, 8, 121–143.

38. Shogren, A. J., Tank, J. L., Egan, S. P., Bolster, D., & Riis, T. (2019). Riverine distribution of mussel environmental DNA reflects a balance among density, transport, and removal processes. Freshwater Biology, 64(8), 1467–1479.

39. Smith, D. R., Villella, R. F., & Lemarié, D. P. (2001). Survey protocol for assessment of endangered freshwater mussels in the Allegheny River, Pennsylvania. Journal of the North American Benthological Society, 20(1), 118–132.

40. Smith, D. R. (2006). Survey design for detecting rare freshwater mussels. Journal of the North American Benthological Society, 25(3), 701–711.

41. Sternhagen, E. C., Davis, M. A., Larson, E. R., Pearce, S. E., Ecrement, S. M., Katz, A. D., & Sperry, J. H. (2024). Comparing cost, effort, and performance of environmental DNA sampling and trapping for detecting an elusive freshwater turtle. Environmental DNA, 6(2), e525.

42. Stoeckle, B. C., Beggel, S., Kuehn, R., & Geist, J. (2021). Influence of stream characteristics and population size on downstream transport of freshwater mollusk environmental DNA. Freshwater Science, 40(1), 191–201.

43. Taberlet, P., Bonin, A., Zinger, L., & Coissac, E. (2018). Environmental DNA: For biodiversity research and monitoring. Oxford University Press.

44. Takahara, T., Yamagishi, S., Shimoda, R., Nagata, A., Sakata, M. K., Doi, H., & Minamoto, T. (2025). Seven-year changes in eDNA concentrations of two dominant submerged macrophytes in Lake Shinji: Effects of salinity. Estuarine, Coastal and Shelf Science, 315, 109165.

45. Tillotson, M. D., Kelly, R. P., Duda, J. J., Hoy, M., Kralj, J., & Quinn, T. P. (2018). Concentrations of environmental DNA (eDNA) reflect spawning salmon abundance at fine spatial and temporal scales. Biological conservation, 220, 1–11.

46. Troth, C. R., Sweet, M. J., Nightingale, J., & Burian, A. (2021). Seasonality, DNA degradation and spatial heterogeneity as drivers of eDNA detection dynamics. Science of the Total Environment, 768, 144466.

47. Van Driessche, C., Everts, T., Neyrinck, S., & Brys, R. (2023). Experimental assessment of downstream environmental DNA patterns under variable fish biomass and river discharge rates. Environmental DNA, 5(1), 102–116.

48. Vaughn, C. C., Taylor, C. M., & Eberhard, K. J. (1997). A comparison of the effectiveness of timed searches vs. quadrat sampling in mussel surveys. In Proceedings of an Upper Mississippi River Conservation Committee (UMRCC) Symposium, 16–18 October 1995 St. Louis, Missouri (pp. 157-162).

49. Villella, R. F., Smith, D. R., & Lemarie, D. P. (2004). Estimating survival and recruitment in a freshwater mussel population using mark-recapture techniques. The American midland naturalist, 151(1), 114–133.

50. USFWS United States Fish and Wildlife Service. (2025). ECOS Environmental Conservation Online System. https://ecos.fws.gov/ecp/. Accessed in November 2025.

51. Wacker, S., Fossøy, F., Larsen, B. M., Brandsegg, H., Sivertsgård, R., & Karlsson, S. (2019). Downstream transport and seasonal variation in freshwater pearl mussel (Margaritifera margaritifera) eDNA concentration. Environmental DNA, 1(1), 64–73.

52. Waits, E. R., Smith, L. M., Patnode, K. A., Clayton, J. L., McGregor, M. A., & Bergdale, A. J. (2025). Development and validation of environmental DNA assays for the detection of endangered and threatened freshwater mussels. Conservation Genetics Resources, 1-8.

53. Watters, G. T., O’Dee, S. H., & Chordas III, S. (2001). Patterns of vertical migration in freshwater mussels (Bivalvia: Unionoida). Journal of Freshwater Ecology, 16(4), 541–549.

54. Watters, G. T., Hoggarth, M. A., & Stansberry, D. H. (2009). The freshwater mussels of Ohio. The Ohio State University Press.

55. Wilcox, T. M., McKelvey, K. S., Young, M. K., Sepulveda, A. J., Shepard, B. B., Jane, S. F., … & Schwartz, M. K. (2016). Understanding environmental DNA detection probabilities: A case study using a stream-dwelling char Salvelinus fontinalis. Biological Conservation, 194, 209–216.

56. Wisniewski, J. M., Rankin, N. M., Weiler, D. A., Strickland, B. A., & Chandler, H. C. (2013). Occupancy and detection of benthic macroinvertebrates: a case study of unionids in the lower Flint River, Georgia, USA. Freshwater Science, 32(4), 1122–1135.

